# Stochastic bacterial population dynamics prevent the emergence of antibiotic resistance from single cells

**DOI:** 10.1101/458547

**Authors:** Helen K. Alexander, R. Craig MacLean

## Abstract

A better understanding of how antibiotic exposure impacts the evolution of resistance is crucial for designing more sustainable treatment strategies. The conventional approach to relating antibiotic dose to resistance evolution within a bacterial population is to measure the range of concentrations over which resistant strain(s) are selectively favoured over a sensitive strain – the “mutant selection window”. Here, we instead investigate how antibiotic concentration impacts the initial establishment of resistance from single cells, mimicking the clonal expansion of a resistant lineage following mutation or horizontal gene transfer. Using two *Pseudomonas aeruginosa* strains carrying distinct resistance plasmids, we show that single resistant cells have <5% probability of outgrowth at antibiotic concentrations as low as 1/8^th^ of the resistant strain’s minimum inhibitory concentration. This low probability of establishment is due to detrimental effects of antibiotics on resistant cells, coupled with the inherently stochastic nature of cell division and death on the single-cell level, which leads to loss of many nascent resistant lineages. Our findings suggest that moderate doses of antibiotics, within the traditional mutant selection window, may be more effective at preventing *de novo* emergence of resistance than predicted by deterministic approaches.

**Significance statement:** The emergence of antibiotic resistance poses a critical threat to the efficacy of antibiotic treatments. A resistant bacterial population must originally arise from a single cell that mutates or acquires a resistance gene. This single cell may, by chance, fail to successfully reproduce before it dies, leading to loss of the nascent resistant lineage. Here we show that antibiotic concentrations that selectively favour resistance are nonetheless sufficient to reduce the chance of outgrowth from a single cell to a very low probability. Our findings suggest that lower antibiotic concentrations than previously thought may be sufficient to prevent, with high probability, emergence of resistance from single cells.

## Introduction

Antibiotics have had a huge impact on human health by reducing the burden associated with bacterial infections, and the use of antibiotics now underpins many areas of medicine. Unfortunately, antibiotic treatment is also associated with the evolution of resistance [1], resulting in poorer patient outcomes [2]. A better understanding of how antibiotic dosing affects resistance evolution could aid the design of more effective treatment strategies that suppress pathogenic bacteria without driving the emergence of resistance.

To date, *in vitro* work addressing how antibiotics affect evolution of resistance has focused on identifying the range of antibiotic concentrations at which resistant mutants are selectively favoured by antibiotic treatment, known as the “mutant selection window” (MSW) [3, 4, 5, 6]. Here we will refer to any strain with reduced susceptibility relative to a wild-type (“sensitive”) strain simply as “resistant”, as is common in evolutionary microbiology literature (e.g. [7, 8, 9]), as opposed to defining resistance with respect to clinical breakpoints. Originally, the lower boundary of the MSW was approximated by the minimum inhibitory concentration of the sensitive strain (MIC_S_) [3, 4, 5, 6], i.e. the lowest antibiotic concentration that abolishes its growth in a standardized assay (such as [10]). However, more recent work has emphasized that a resistant strain can be selectively favoured down to a minimal selective concentration (MSC) that is often well below MIC_S_ [11, 12, 13, 7, 8]. The upper boundary of the MSW is conventionally defined by the lowest antibiotic concentration that prevents growth of all mutant subpopulations [3, 4]. This upper bound is often equated with the minimum inhibitory concentration of the most resistant single-point mutant [5] or of a specific resistant strain under study [13, 8], which we denote MIC_R_. The MSW therefore ranges between antibiotic concentrations that are so low that they are unlikely to have any clinical benefit (below MIC_S_) and very high concentrations (up to MIC_R_) that may be difficult to achieve in practice because of physiological constraints on the accumulation of antibiotics in tissues (pharmacokinetics) and toxic side-effects of antibiotics [14, 15].

Selection operates efficiently when both sensitive and resistant populations are large, resulting in an increase in frequency of the fitter strain in an antibiotic dose-dependent manner. Correspondingly, the mutant selection window is typically measured by direct competition between large numbers of cells (typically >10^4^ colony-forming units, CFU) of both resistant and sensitive strains across a gradient of antibiotic concentrations (e.g. [13]). This provides a powerful approach to measure the impact of antibiotic dose on selection for established resistant strains, i.e. those that are already reasonably prevalent. However, *de novo* emergence of resistant strains, when not initially present, should be subject to stochastic processes [16] that are not reflected by the MSW, nor captured by this experimental design. First, resistance must stochastically arise in a sensitive cell by mutation, or by acquisition of a resistance gene through horizontal gene transfer. Next, the single resistant cell thus generated must survive and successfully divide to produce daughter cells that likewise survive, and so on to generate a large number of resistant descendant cells. The latter process, which we will refer to throughout as “establishment” of resistance [16], will be our focus here. Importantly, due to the stochastic nature of cell divisions and deaths on the individual level, establishment is not guaranteed, even under conditions in which the resistant strain is selectively favoured [17]. Moreover, if antibiotics even partially inhibit the resistant strain below its MIC_R_, the chance that a resistant cell dies or fails to divide, and thus the risk that a resistant lineage is stochastically lost, should increase in the presence of antibiotics at concentrations within the MSW. Despite the substantial body of work addressing the selection of resistance, very little work has addressed the stochastic establishment phase (see however [18, 19, 20]).

We set out to quantify stochastic establishment using *in vitro* experiments with *Pseudomonas aeruginosa*, an important opportunistic pathogen that evolves resistance at an exceptionally high rate during infections [1, 21]. To isolate the establishment phase, we inoculated hundreds of cultures, each with a very small number of resistant cells (on average, approx. 1-3), and assessed culture growth. We tested two strains carrying non-conjugative plasmids (Rms149 and PAMBL2) that confer resistance to streptomycin and meropenem, respectively, across a gradient of the corresponding antibiotic concentrations within the MSW. By fitting mathematical models to these data, we estimated the probability of establishment, i.e. of detectable culture growth due to clonal expansion from a single resistant cell, as a function of antibiotic concentration. Our key finding is that the establishment probability of resistant cells drastically declines at concentrations well below the MIC of the resistant strain, reaching ≲5% at 1/8^th^ of MIC_R_ in both systems. These concentrations lie well above the corresponding MIC values of the sensitive strain and within the conventional mutant selection window. Our results highlight that antibiotic selection pressure is not a sufficient condition for *de novo* emergence of resistance starting from single cells. Accounting for the demographic stochasticity inherent to the outgrowth of mutant lineages substantially narrows the window of concentrations at which resistant mutants are likely to establish, suggesting that moderate antibiotic dosing may be an effective strategy to prevent the emergence of resistance.

## Results

### Establishment of resistance is inhibited by sub-MIC_R_ antibiotic concentrations

To elucidate the direct impact of antibiotics on resistant cells, we first investigated establishment of a resistant strain in the absence of a sensitive strain. We first focused on the streptomycin-resistant PA01:Rms149 strain. To estimate its probability of establishment, defined as outgrowth of a detectable (i.e. large) population from a single cell, we conducted large-scale “seeding” experiments (see also [20]). In this assay (**Fig. 1**), a highly diluted overnight culture of the resistant strain is inoculated into fresh media in a large number of replicate cultures. The high dilution factors yield average inoculum sizes of <1 to ∼3 cells per culture. Importantly, however, the actual number of cells inoculated into each replicate culture is random, and can be described by a Poisson distribution (**SI Appendix, Fig. S1**). One implication of this protocol is that many cultures are not inoculated with any cells, while others receive more than one cell; our modelling approach will account for this variation statistically. We inoculated parallel replicate cultures in streptomycin-free media and at a range of streptomycin concentrations below the MIC of the resistant strain, denoted MIC_R_, as measured using standard protocols [10] (**SI Appendix, Table S1**). We then scored the number of replicate cultures showing growth based on reaching a threshold optical density (OD) of 0.1 within 3 days post-inoculation.

**Figure 1:**
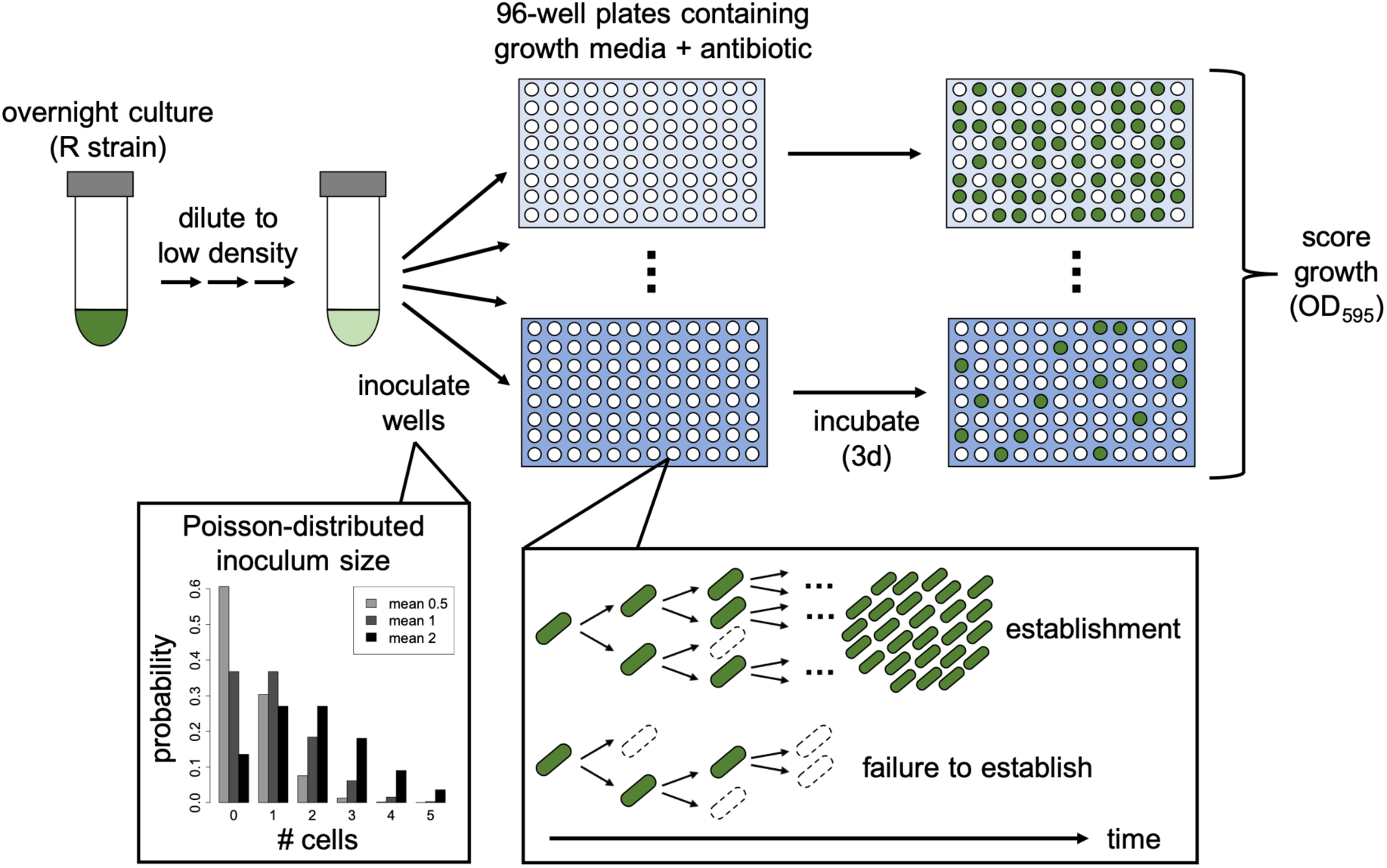
Design of seeding experiments to estimate establishment probability. An overnight culture of the resistant strain is highly diluted and used to inoculate 96-well plates containing growth media (LB broth) with antibiotic at various concentrations (shades of blue). The number of cells inoculated per well follows a Poisson distribution (examples plotted for mean inoculum size of 0.5, 1, or 2 cells per well). Within these culture wells, stochastic population dynamics imply that each inoculated cell may either produce a large number of descendants (*establishment*) or produce no/few descendants that ultimately die out (*failure to establish*). Plates are incubated for 3 days and optical density is measured to score growth in wells (OD_595_ > 0.1; dark green). The number of replicate cultures showing growth is used to estimate the per-cell establishment probability at each antibiotic concentration by fitting a mathematical model.

A culture could fail to grow either because the inoculum did not contain any cells, or because every cell in the inoculum failed to give rise to a surviving lineage. To infer the probability that a single cell yields detectable population growth (i.e. the per-cell establishment probability), we fit a mathematical model, accounting for both the random inoculum size and demographic stochasticity, to the observed number of replicate cultures showing growth (**Materials and Methods**). All probabilities are normalized by the result in streptomycin-free media, which corresponds to scaling inoculum size by the mean number of cells that establish in benign conditions (which we call the “effective” inoculum size). Thus, relative establishment probability 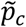 equals one by definition in streptomycin-free conditions, while we expect 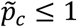 with streptomycin treatment; however, values larger than one can arise due to sampling error.

Our seeding experiments revealed that the probability of establishment of a single resistant cell declines with increasing streptomycin concentration (**Fig. 2** and **SI Appendix, Table S2**). While exposure to the lowest tested concentrations of streptomycin (up to 1/32 × MIC_R_) had no detectable impact on establishment, 1/16 × MIC_R_ was already sufficient for significant declines, to 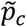 of 55-73% (maximum likelihood estimates in two independent experiments). At 1/8 × MIC_R_, 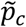 dropped to just 3-5%. These results suggest that a *de novo* resistant mutant would only rarely establish at antibiotic concentrations that are well below its MIC_R_, i.e. within the MSW.

**Figure 2:**
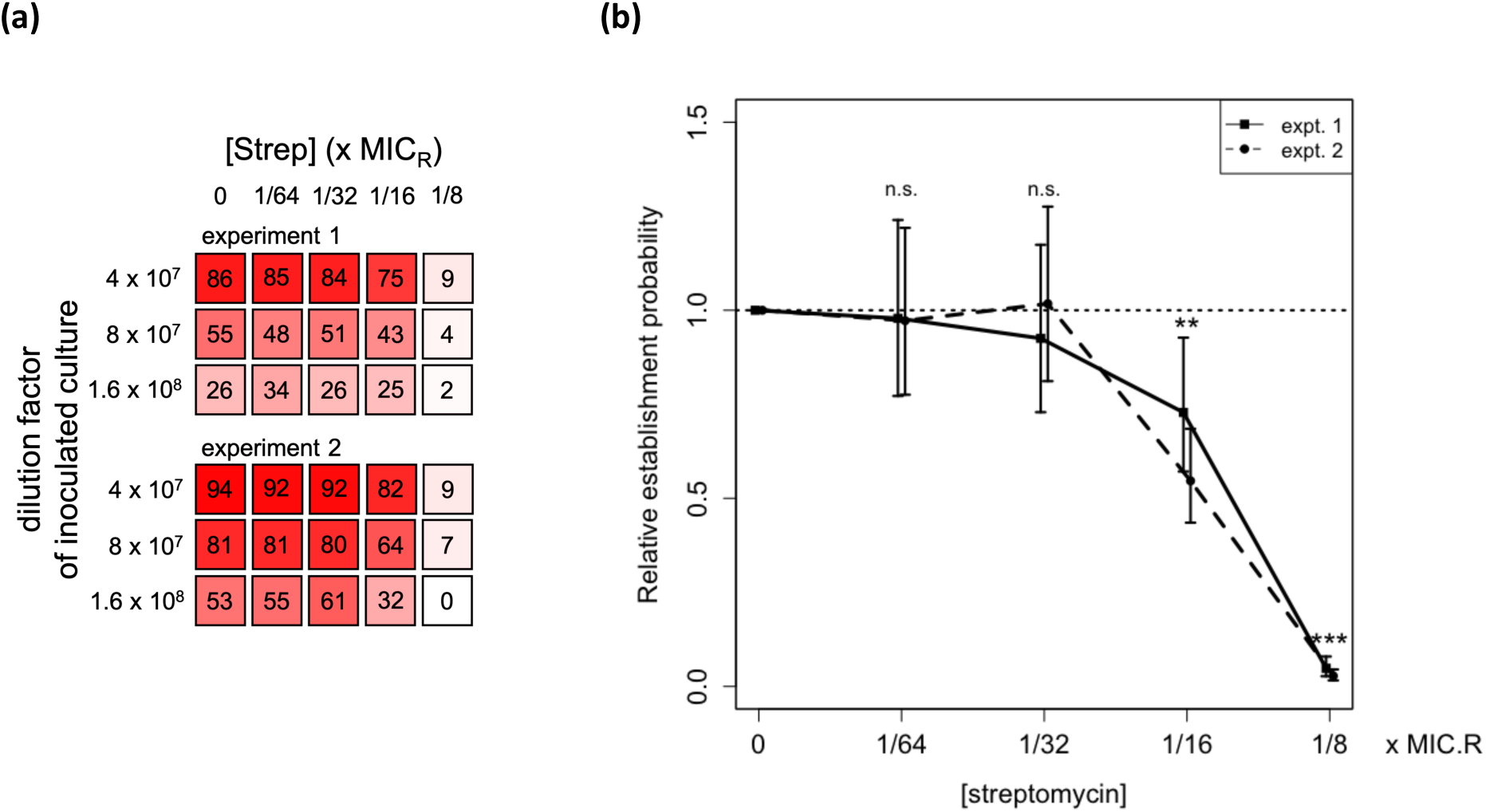
Establishment probability of single PA01:Rms149 streptomycin-resistant cells, estimated from seeding experiments. (a) Visual representation of the growth data, indicating the number of replicate cultures (out of 96) that grew in each test condition up to 3d post-inoculation. **(b) Estimated relative per-cell establishment probability 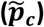**, scaled by the probability in streptomycin-free medium, as a function of streptomycin concentration, scaled by the standard MIC value of the resistant strain (MIC_R_ = 2048μg/ml; **Table S1**). Results are shown for two separate experiments. Plotted points indicate the maximum likelihood estimate of 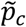 and error bars indicate the 95% confidence interval, using the fitted model selected by the likelihood ratio test (experiment 1: Model B’, fixed environmental effect; experiment 2: Model C’, the null model [**Eqn. 1**]. Both of these models pool data across three inoculation densities; see **Suppl. Text**, section 10, for details). Significance of the streptomycin effect is determined by fitting a generalized linear model to the population growth data (n.s.: not significant, *p* > 0.05; ** *p*=0.01 in expt. 1, *p*=2e-7 in expt. 2, and *p*=2e-8 pooling both experiments; *** *p* < 2e-16 in both experiments; see **Suppl. Text**, section 14.1, for full results).

### MIC depends on inoculum size

The frequent failure of the resistant strain to grow in our seeding experiments at concentrations well below its MIC is, at face value, surprising. We hypothesized that these results could be explained by the difference in inoculum size between these assays. Specifically, standard MIC values are assessed from an inoculation density of 5×10^5^ CFU/mL [10], which corresponds to an inoculum size of 10^5^ CFU per 200μl culture on our microtitre plates. In contrast, our seeding experiments used an inoculum size on the order of 1 CFU per culture. MIC for many antibiotics has been observed to increase with higher-than-standard inoculation densities (CFU/ml) [22, 23, 24] which corresponds to higher absolute inoculum size (CFU) for a fixed culture volume. Although less well-explored, it has also occasionally been noted that MIC can decrease when lower absolute inoculum sizes are used [25, 26].

To test the hypothesis that inoculum size influences MIC in the present system, we conducted a modified MIC assay using the PA01:Rms149 strain with inoculum sizes ranging over three orders of magnitude, from approximately 10^2^ to 10^5^ CFU per culture (corresponding to inoculation densities of 5×10^2^ up to the standard 5×10^5^ CFU/ml). We found that MIC indeed increases with inoculum size (**Fig. 3a**). This pattern arises regardless of whether growth is scored at 20h, as per the standard MIC assay protocol [10], or up to 3d post-inoculation, as in our seeding experiments, although the number of cultures showing detectable growth, and thus the measured MIC, tends to increase over time (**SI Appendix, Fig. S2**).

**Figure 3:**
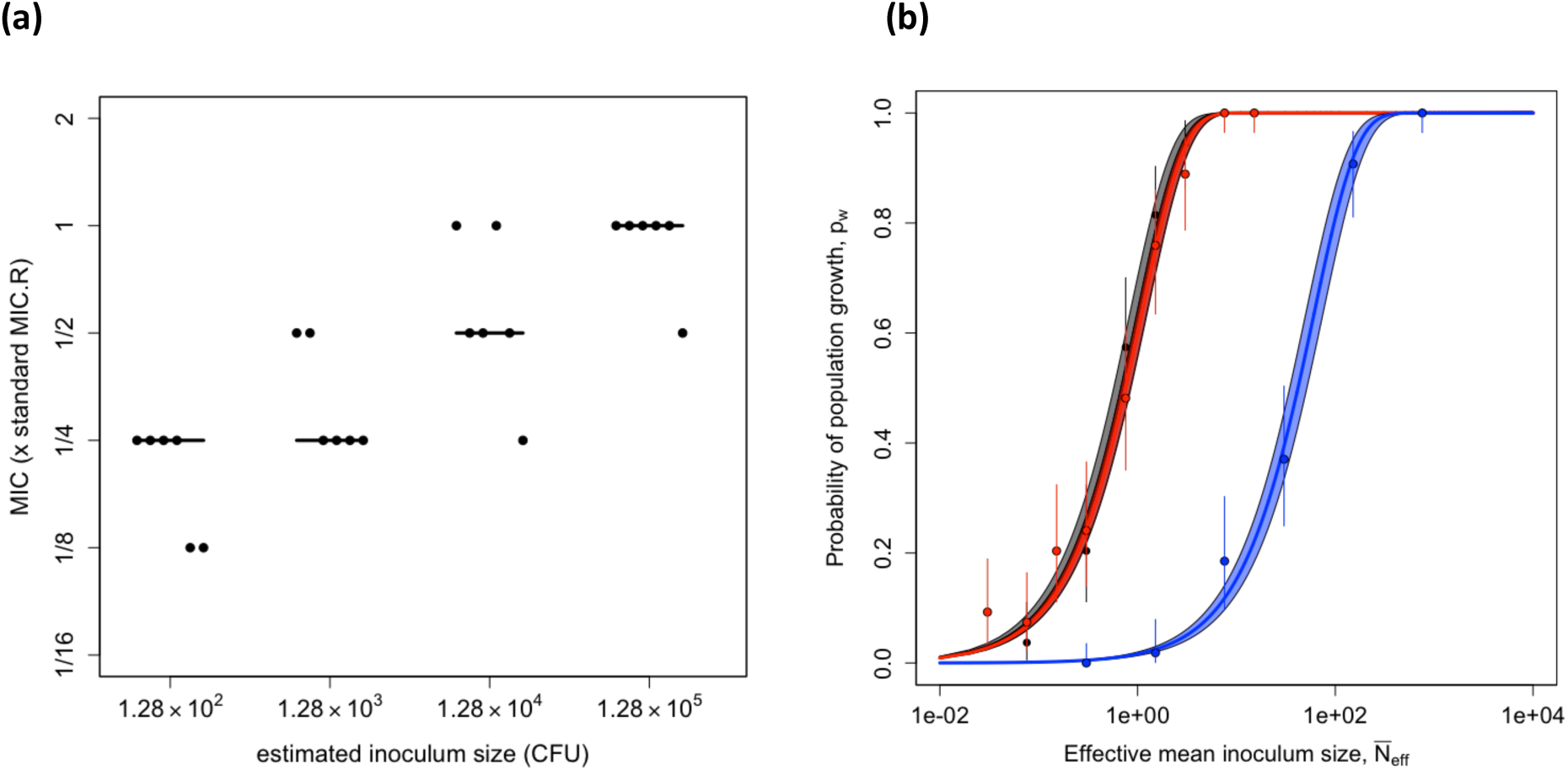
Inoculum size effects on MIC and probability of population growth of the resistant PA01:Rms149 strain in streptomycin. (a) MIC as a function of inoculum size. Cultures were inoculated with PA01:Rms149 at four different inoculum sizes. MIC was evaluated as the minimal tested streptomycin concentration that prevented detectable growth up to 3d post-inoculation; a qualitatively similar pattern arose if growth was evaluated at 20h (**Fig. S2**). The y-axis is scaled by the MIC of this strain at standard inoculation density (MIC_R_). The points represent six replicates at each inoculum size, with the line segments indicating their median. **(b) Null model of the inoculum size effect (Eqn. 1) fit to culture growth data**. Probability of population growth (*p*_*w*_) is plotted as a function of effective mean inoculum size (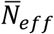, calibrated by the results in streptomycin-free media; see **Fig. S4**). Black: streptomycin-free; red: streptomycin at 1/16 × MIC_R_; blue: 1/8 × MIC_R_. These results are based on growth in streptomycin up to 5d post-inoculation; see **Suppl. Text**, section 15, for results at 3d post-inoculation. Points indicate the proportion of replicate cultures showing growth, i.e. the maximum likelihood estimate (MLE) of *p*_*w*_ in the full model, with error bars indicating the 95% confidence interval (CI). The solid line shows the best fit of the null model (i.e. **Eqn. 1** parameterized with the MLE of 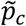) and the shaded area corresponds to the 95% CI. According to the likelihood ratio test, the null model deviance from the full model is not significant at any streptomycin concentration (streptomycin-free: *p*=0.55; 1/16 × MIC_R_: *p*=0.28; 1/8 × MIC_R_: *p*=0.71; see **Suppl. Text**, section 15, for full results).

Since all cultures contained the same volume in the above experiment, this pattern could be due to changes either in absolute inoculum size (i.e. CFU) or in inoculation density (i.e. CFU per unit volume). These two possibilities are not typically distinguished in the literature; however, they lead to distinct interpretations. If demographic stochasticity is the dominant force, we expect absolute numbers to matter, whereas if interactions among cells (e.g. competition or cooperation) affect establishment, cell density per unit volume could be more important. To disentangle these two factors, we repeated the MIC assay co-varying inoculation density and culture volume. This experiment confirmed that absolute inoculum size has a strong effect on MIC. In contrast, inoculation density per unit volume does not have a significant effect within the range that we tested, after controlling for absolute cell numbers (**SI Appendix, Fig. S3**).

### Population growth can be explained by an independent chance of each cell to establish

Taken together, our seeding experiments and MIC assays reveal that the absolute number of cells in the inoculum has a strong effect on whether the culture eventually shows detectable growth. The simplest explanation for this result is that population growth can be attributed to the stochastic outgrowth of one or more lineages, each initiated by a single cell in the inoculum, acting independently. This independence assumption yields a “null model” that mathematically describes the effect of inoculum size on the probability of outgrowth of a detectable population (**Materials and Methods, Eqn. 1**). Here the probability of establishment of each cell in the inoculum 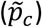 is a scaling parameter, which does not depend on inoculum size. Note that this null model would not hold if interactions among cells substantially influenced their chances of successful replication. For example, if cells secrete an enzyme that breaks down an antibiotic extracellularly, then the establishment probability of each cell could increase with inoculum size. On the other hand, if cells compete for limiting resources or secrete toxins, the per-cell establishment probability could decrease with inoculum size.

To formally test the null model, we again conducted seeding experiments with the PA01:Rms149 strain, but now using many different inoculum sizes, spanning approximately three orders of magnitude. We tested two streptomycin concentrations (1/16 and 1/8 × MIC_R_) for which growth often failed from a single cell, but succeeded from standard inoculum size in MIC assays. In parallel, we tested growth in streptomycin-free media in order to estimate the effective mean inoculum size (**SI Appendix, Fig. S4**). This left one free parameter, the per-cell relative establishment probability 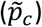, to fit at each streptomycin concentration.

We found good agreement between the null model and our experimental data at all tested streptomycin concentrations, consistent with the hypothesis that cells establish independently (main experiment, **Fig. 3b**, and repeat experiments, **SI Appendix, Fig. S5**). More precisely, the null model did not show significant deviance from the observed proportion of populations that grew (according to the likelihood ratio test), and thus we accept it as a parsimonious explanation for the data. Furthermore, we obtain estimates of relative establishment probability, 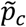, at 1/16 and 1/8 × MIC_R_ similar to those from the previous seeding experiments (**SI Appendix, Table S2**).

To summarize, the probability of culture growth at any given streptomycin concentration depends on inoculum size, according to a simple quantitative relationship. Our experimental data are consistent with a simple model of cells behaving independently, such that a fixed per-cell establishment probability can explain our growth data across inoculum sizes. That is, cells are not “more susceptible” to streptomycin at lower inoculum sizes, but rather, culture growth is less likely to be observed simply because fewer cells are available to establish, and not all cells succeed. In turn, the minimal concentration of streptomycin required to prevent growth in some proportion of replicate cultures (i.e. the observed MIC) increases with inoculum size.

### Sub-MIC_R_ streptomycin concentrations induce resistant cell death and extend lag phase

We hypothesized that resistant cells sometimes failed to establish in our seeding experiments because exposure to streptomycin compromised cell division rate and/or viability. As a simple test of this idea, we measured the relative abundance of dead cells in cultures of the resistant strain grown at sub-MIC_R_ concentrations of streptomycin. We found that the fraction of dead cells after 7h of treatment, as determined by propidium iodide staining, increased from an average of 3-4% in streptomycin-free conditions to >20% at 1/8 × MIC_R_ streptomycin (**Fig. 4a** and **SI Appendix, Fig. S6** and **Table S3**). Note that this is a conservative measure of cell death, because this assay only detects cells that have compromised membrane permeability, and not, for example, cells that have already lysed. Furthermore, this assay provides only a snapshot in time.

**Figure 4:**
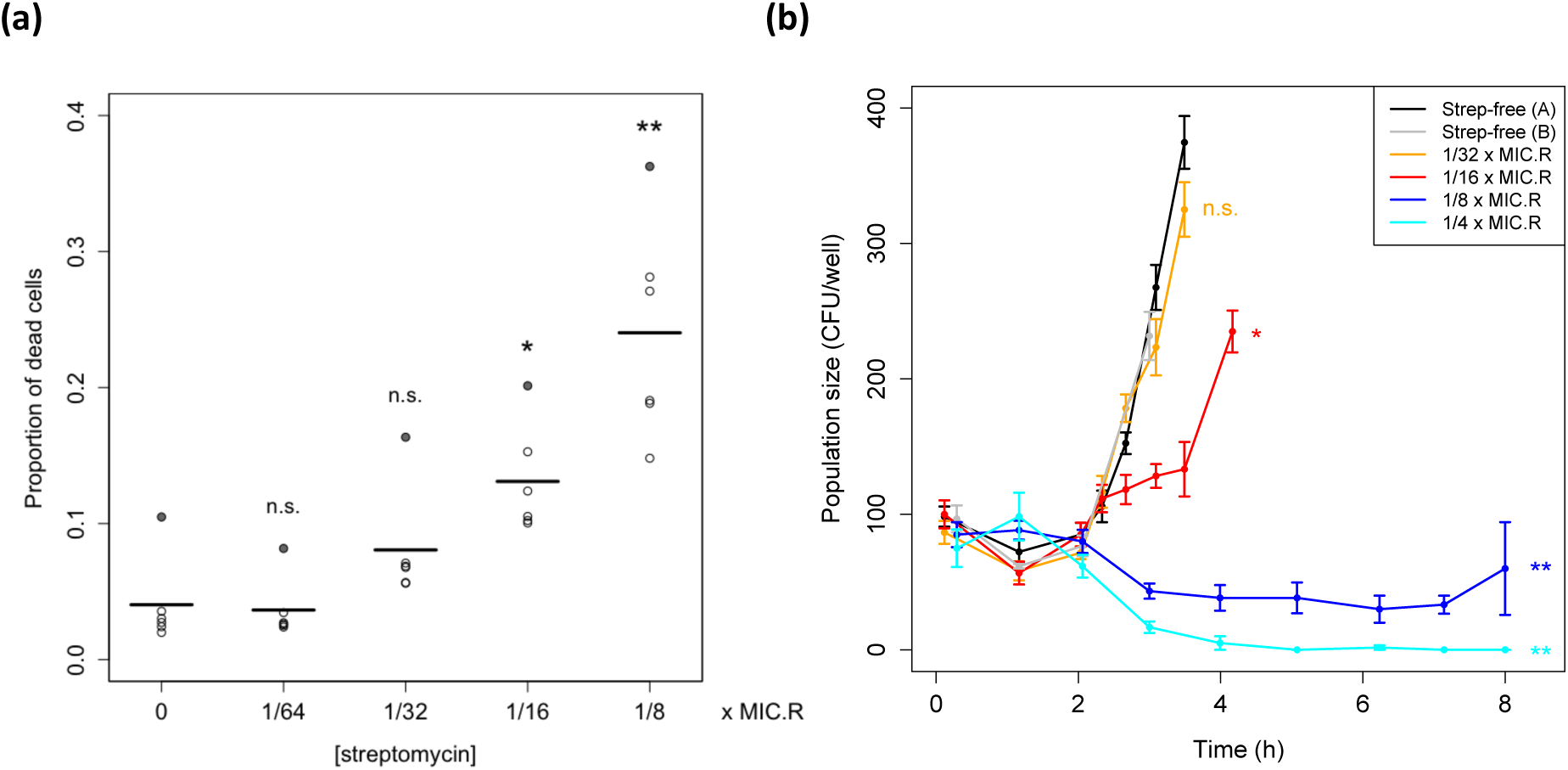
Effects of sub-MIC_R_ streptomycin treatment on PA01:Rms149 resistant cell dynamics. (a) Proportion of dead cells after 7h in sub-MIC_R_ streptomycin. The proportion of dead cells in streptomycin-treated cultures was estimated using live-dead staining and flow cytometry. Points represent six independent treatment replicates at each concentration and line segments indicate their mean. Differences from the streptomycin-free control cultures were assessed using a one-way ANOVA followed by a post-hoc Dunnett’s test (n.s.: not significant, *p* > 0.05; *: *p* = 9e-3, **: *p* < 1e-4). Effects identified as significant do not change if we exclude an outlier replicate (shaded-in points) showing consistently elevated dead cell fractions (**Table S3**). **(b) Viable cell population dynamics in sub-MIC**_**R**_ **streptomycin**. Points with connecting lines indicate the mean number of viable cells across six replicate cultures per streptomycin concentration, per sampling time point (or twelve replicates for streptomycin-free controls); the error bars indicate standard error. **Fig. S7** shows all individual replicates. Viable cell numbers were estimated by plating undiluted culture samples; plots are truncated when colonies became too dense to count. Significance of each streptomycin concentration compared to the streptomycin-free control was assessed by a post-hoc Dunnett’s test (n.s.: not significant, *p*=0.87; * *p*=4e-4; ** *p*<1e-4).

To gain further insight into how sub-MIC_R_ streptomycin impacts the population dynamics of the resistant strain, we quantified viable cell density over the first few hours after inoculation into streptomycin-containing media. Cultures were inoculated with approximately 100 cells in this experiment, to ensure that cell numbers were low enough for demographic stochasticity to be relevant, yet large enough to be detectable using conventional plating methods.

We found that streptomycin treatment has a significant effect on the growth of resistant cultures (ANOVA, main effect: *p* < 2e-16), and this effect varies over time (ANOVA, interaction term: *p* < 2e-16; **Fig. 4b** and **SI Appendix, Fig. S7**). Following inoculation, cultures exhibited a lag phase of approximately 2 hours. Control cultures in streptomycin-free media then began to grow exponentially. The lowest tested concentration of streptomycin (1/32 × MIC_R_) had no significant effect on these dynamics (Dunnett’s test: *p*=0.87); however, 1/16 × MIC_R_ was already sufficient to slow growth (*p*=4e-4). Nonetheless, all replicate cultures (*n*=48 per concentration) eventually grew, as detected by OD. Meanwhile, higher doses of streptomycin (1/8 × or 1/4 × MIC_R_) had dramatic effects on growth dynamics (*p* < 1e-4), with cultures exhibiting an extended lag phase of at least 7-8 hours, in which viable cell density initially declined. After further incubation (up to 3 days), 25% of cultures (15/60) exposed to 1/8 × MIC_R_ eventually showed growth, while the remaining 75% (45/60) failed to reach detectable OD. At 1/4 × MIC_R_, no viable cells were detected in most cultures from 4h on, and only 1/60 cultures reached detectable OD within 3 days.

In summary, sub-MIC_R_ streptomycin treatment has the effect of extending the lag phase, before cultures eventually either grow to saturation or die out. Failure to grow can be explained by significantly elevated cell death rates beginning at 1/16 × MIC_R_, which can lead to stochastic loss of initially small populations.

### Stochastic establishment is recapitulated for a clinically relevant antibiotic and resistance plasmid

If the frequent failure of resistant cells to establish surviving populations at antibiotic doses well below their MIC is a general phenomenon, it would have important implications for understanding the emergence of resistance during antibiotic treatment. To confirm that our result was not driven by the specific choice of antibiotic or resistance mechanism, we repeated the key seeding experiment using a *P. aeruginosa* PA01 strain carrying a recently isolated multi-drug resistance plasmid, PAMBL2 [27, 28], that confers resistance to meropenem through the *bla*_vim-1_ carbapenemase. Carbapenems are an important treatment option for serious infections caused by gram-negative bacterial pathogens, and resistance is of current clinical concern [29, 30]; carbapenem-resistant *P. aeruginosa* has been identified as a “critical priority” for new antibiotic development by the WHO [31]. In agreement with our previous findings, the establishment probability of PA01:PAMBL2 cells declined at concentrations of meropenem well below this strain’s MIC_R_ (**SI Appendix, Table S1**), reaching ∼5% at 1/8 × MIC_R_, while no establishment was observed at 1/4 × MIC_R_ (**Fig. 5** and **SI Appendix, Table S4**). This result highlights that the stochastic loss of resistant cells is not unique to our primary model system of PA01:Rms149 in streptomycin.

**Figure 5:**
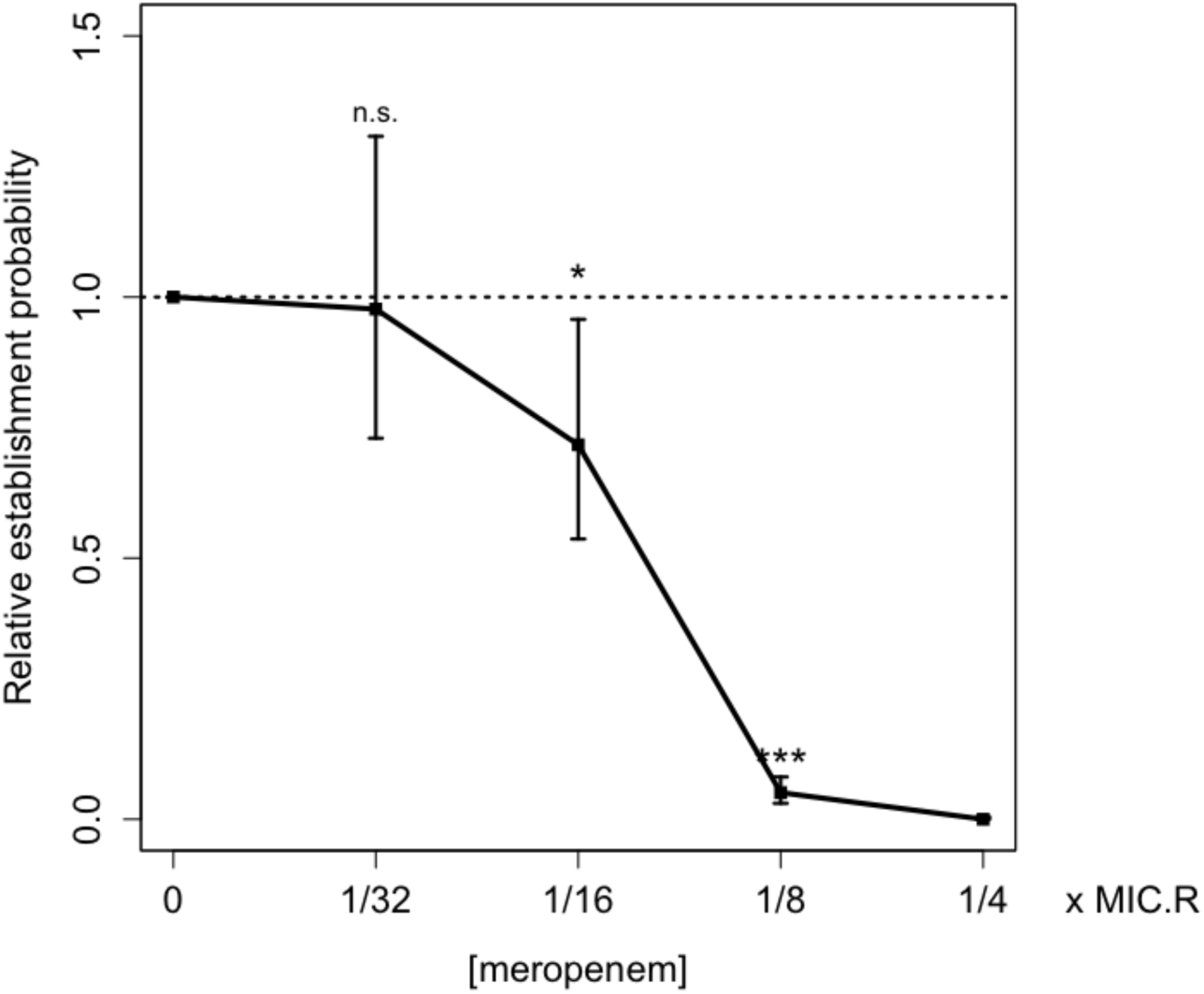
Estimated relative per-cell establishment probability of the PA01:PAMBL2 meropenem-resistant strain as a function of meropenem concentration. Concentration is scaled by the standard MIC of this strain in meropenem (MIC_R_ = 512 μg/ml; **Table S1**). Plotted points indicate the maximum likelihood estimate of 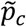 and error bars indicate the 95% confidence interval, using the fitted model selected by the likelihood ratio test (Model C’, the null model [**Eqn. 1**], which pools data across two tested inoculation densities). Significance of the meropenem effect is determined by fitting a generalized linear model (GLM) to population growth data (n.s.: not significant, *p* > 0.05; * *p* = 0.02; *** *p* < 2e-16; see **Suppl. Text**, section 14.2, for full results). 1/4 × MIC_R_ meropenem was excluded from the GLM because zero replicates established.

### The sensitive population modulates probability of establishment of resistant cells

So far, we focused on the direct effects of antibiotics on resistant cells by conducting experiments with monocultures of resistant strains. However, *de novo* resistance will actually arise within a sensitive population, by mutation or transfer of a mobile genetic element into a sensitive cell. Moreover, antibiotic treatment will only begin in clinical settings once the total pathogen population is large enough to cause symptoms. We therefore asked whether the presence of a large sensitive population affects the establishment of initially rare resistant cells during antibiotic treatment, returning to the interaction between PA01 (sensitive) and PA01:Rms149 (streptomycin-resistant) as a model system.

We expect the sensitive population and the antibiotic to have interacting effects on establishment of resistance. In particular, at sufficiently low antibiotic concentrations, a sensitive strain is generally expected to outcompete a resistant strain due to the fitness cost associated with resistance [13, 7, 8]. We confirmed this expectation in our experimental system using a standard competition assay, where both strains start from reasonably large inoculum sizes (**SI Appendix, Fig. S8-S10** and **Table S5**). We found that the sensitive strain is favoured up to a minimum selective concentration (MSC) between 1-2 μg/ml streptomycin (equivalent to 1/32 – 1/16 × MIC_S_, or 1/2048 – 1/1024 × MIC_R_), in agreement with previous results for these strains [32]. We hypothesized that competition from the sensitive strain would prevent establishment of resistance at streptomycin concentrations below the MSC.

As a simple test of this idea, we modified the seeding experiment to inoculate very few resistant cells into a large sensitive population. Since bacterial densities in clinical infections can vary widely [9, 33], we inoculated the sensitive strain at two different densities: approximately 5 × 10^5^ CFU/ml (as in a standard MIC assay; labelled “low”) and 5 × 10^7^ CFU/ml (labelled “high”). The resistant strain was seeded, with mean inoculum size on the order of one cell per culture, immediately thereafter.

As hypothesized, we found that the presence of the sensitive population (at either density) abolished establishment of resistant cells in the absence of streptomycin (**Fig. 6** and **SI Appendix, Fig. S11** and **Table S6**). Meanwhile, at streptomycin concentrations above the MSC (1/256 to 1/8 × MIC_R_, or 1/4 to 8 × MIC_S_), adding the sensitive population at low density had a negligible effect on the probability of establishment of resistant cells. At high density, the sensitive population also had negligible effects on establishment of resistance at streptomycin concentrations up to 1/16 × MIC_R_ (4 × MIC_S_). However, at 1/8 × MIC_R_ (8 × MIC_S_), the presence of a high-density sensitive population increased the establishment probability from near zero to 65%. To confirm and further probe the extent of this apparent protective effect, we repeated the experiment over a higher range of streptomycin concentrations. The boost in establishment probability was repeatable and highly significant at 1/8 × MIC_R_ (Wilcoxon rank-sum test, high-vs. zero or low-density sensitive: *p* < 5e-8 in both experiments). However, at 1/4 × MIC_R_ (16 × MIC_S_), an apparent slight boost in establishment probability was non-significant, and by 1/2 × MIC_R_ (32 × MIC_S_) the effect was abolished. Thus, a sufficiently dense sensitive population can extend the range of streptomycin concentrations at which the resistant strain is likely to emerge, but does not change the qualitative pattern of stochastic establishment.

**Figure 6:**
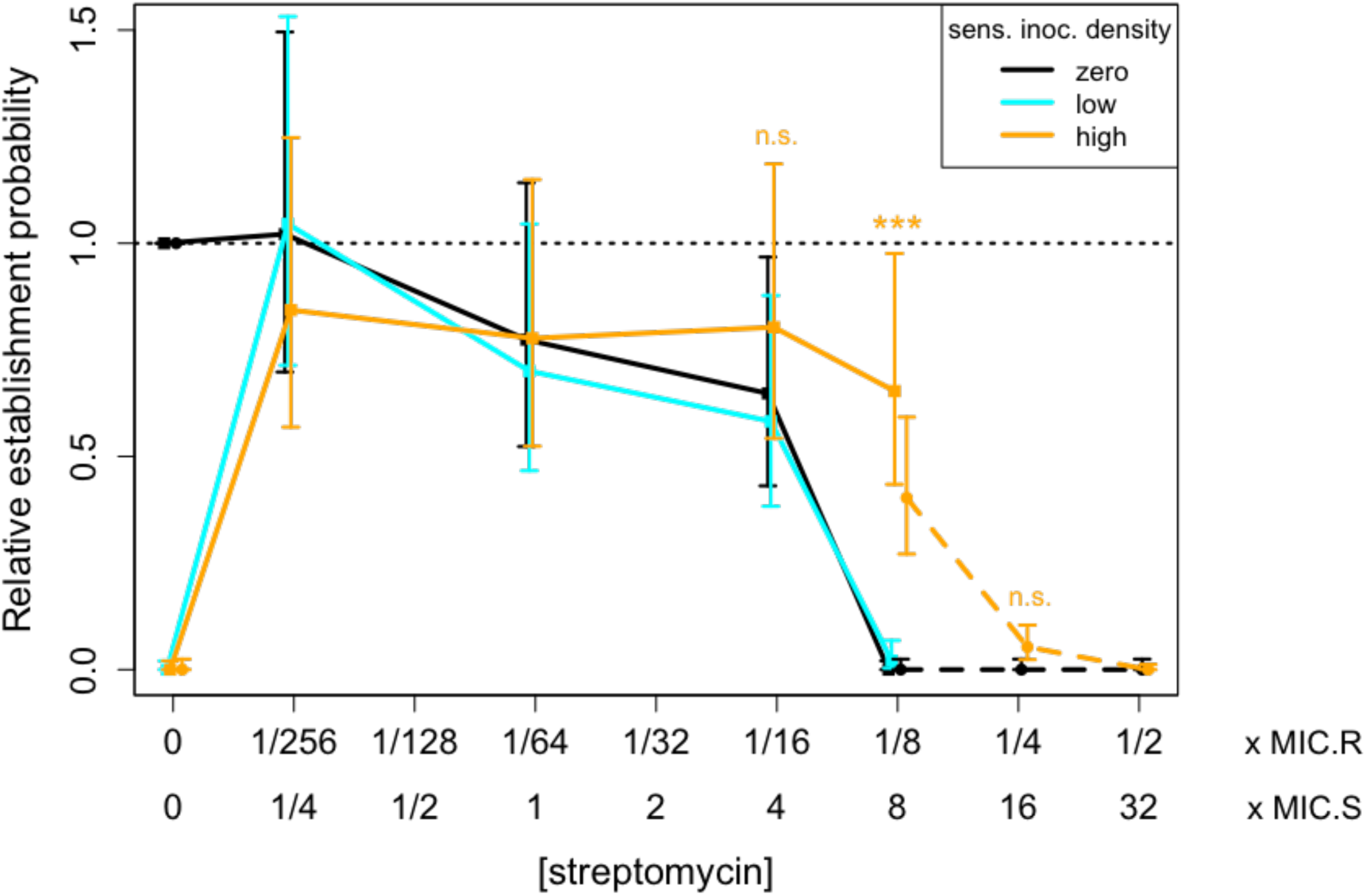
Impact of a large sensitive population on the establishment probability of a resistant cell. The PA01:Rms149 resistant strain was seeded either alone (black) or into a low-density (cyan) or high-density (orange) sensitive PA01 population, across a range of streptomycin concentrations. Results are shown from two separate experiments, testing different subsets of conditions (experiment 1 – data points in squares with solid line; experiment 2 – data points in circles with dashed line). Within each experiment, the estimated relative establishment probability per resistant cell 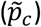 in each condition is normalized by the result for the resistant strain alone in streptomycin-free media. Points indicate the maximum likelihood estimate of 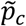 and error bars indicate the 95% confidence interval, using the fitted model selected by the likelihood ratio test (Model C’, the null model [**Eqn. 1**] for both experiments). At streptomycin concentrations of particular interest, the number of replicates in which the resistant strain established in the presence of no or low-density sensitive (pooled where applicable) vs. high-density sensitive was compared using a two-sided Wilcoxon rank-sum test, with significance annotated on the plot (1/16x MIC_R_: experiment 1, *p*=0.17; 1/8x MIC_R_: experiment 1, *p* < 2.2e-16 and experiment 2, *p*=4.8e-8; 1/4x MIC_R_: experiment 2, *p*=0.042, not significant after Bonferroni correction); see **Suppl. Text**, section 16, for further details.

## Discussion

In order for resistance to emerge *de novo*, not only must a resistance gene arise in a bacterial population by mutation or horizontal gene transfer; this first resistant cell must also successfully expand to form a large population. Since any individual cell may fail to replicate, particularly in challenging environmental conditions, the expansion of newly arisen resistant strains is not guaranteed. Our key finding is that demographic stochasticity imposes a significant barrier to the emergence of resistance in the presence of antibiotics at concentrations within the mutant selection window.

We empirically demonstrated the importance of stochasticity with a simple “seeding experiment” mimicking the growth of clonal resistant lineages founded by a single cell. First, to assess the direct impact of antibiotics, we inoculated fresh antibiotic-containing media with approximately one resistant cell per replicate culture and quantified the per-cell probability of establishing a detectable population. Strikingly, this establishment probability dropped off at concentrations well below the MIC of the corresponding resistant strain (MIC_R_). For example, the establishment probability of PA01:Rms149 was significantly reduced by streptomycin concentrations as low as 1/16 × MIC_R_, and dropped to <5% at 1/8 × MIC_R_ (**Fig. 2**). Resistant cells failed to establish viable populations because of the toxic effects of exposure to sub-MIC_R_ concentrations of antibiotics (**Fig. 4**) coupled with the inherently stochastic nature of individual cell death and division. Importantly, we were able to replicate our key finding of frequent stochastic loss using a different, meropenem-resistant strain (PA01:PAMBL2; **Fig. 5**). This demonstrated that our results are not limited to a particular model system, but are also relevant to bacterial pathogens of clinical concern: carbapenem-resistant *P. aeruginosa* is considered by the WHO to be of “critical priority” for antibiotic development [31].

In clinical settings, antibiotic treatment will typically begin only when the total bacterial population is large enough to cause symptoms. Assuming resistance has not been transmitted, this population will be predominantly antibiotic-sensitive, and *de novo* mutation or acquisition of a mobile element conferring resistance will occur within a sensitive cell. We therefore next asked how the presence of a large sensitive population would combine with the above effects of antibiotics to shape the emergence of resistance from this first cell, again in the streptomycin model system. As predicted from standard competition assays, emergence of resistance was abolished in the absence of antibiotics (**Fig. 6**), presumably due to competitive suppression by the sensitive strain [8]. More interestingly, a sufficiently dense sensitive population (inoculated at ∼5 × 10^7^ CFU/ml here) was able to shift the range of concentrations at which resistance established upwards by approximately two-fold. We speculate that this apparent protection is due to sensitive cells absorbing antibiotics, thus lowering their concentration in the media [23, 34], despite these concentrations being high enough to cause decline of the sensitive population (i.e. > MIC_S_). *A priori*, one may not expect resistant cells to “need” protection at sub-MIC_R_ antibiotic concentrations. However, in the stochastic regime of establishment, any increase in the probability of individual cells surviving and dividing can make a qualitative difference to the fate of a rare resistant lineage. This protective effect depends on bacterial density at the time of treatment, within a realistic range for some bacterial infections [9, 33]: when the sensitive population was inoculated at 100-fold lower density (∼5 × 10^5^ CFU/ml), protection was not apparent. We emphasize that although these experiments provide an initial proof of concept, a complete investigation of the interacting effects of sensitive population density, antibiotic dose and timing remains an important direction for future work. Importantly, however, our main message continues to hold even in the more realistic context of resistance arising within a sensitive population: stochastic loss of resistant cells is frequent at antibiotic concentrations within the MSW.

The failure of resistant cells to establish successful lineages at concentrations well below the MIC_R_ shows a clear disconnect between antibiotic susceptibility of individual cells and populations. To explain this effect rigorously, we quantified the probability of outgrowth of a detectable population at a fixed streptomycin concentration, starting from inoculum sizes spanning three orders of magnitude. We fit these data to a mathematical model relating inoculum size to probability of population growth, under the hypothesis that each cell in the inoculum behaves independently (**Eqn. 1**). This simple stochastic model, with a constant per-cell probability of establishment 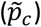, provides a good explanation for inoculum size-dependent population growth in PA01:Rms149 (**Fig. 3b**). In this model, individual cells are not “more susceptible” to antibiotic in smaller populations. Instead, the cumulative effect of many cells, each with a small chance of establishment (e.g. <5% at 1/8 × MIC_R_ in this system), virtually guarantees population growth from a sufficiently large inoculum size, reconciling our results with the standard definition of the MIC. We thus emphasize that MIC is not an innate property of a cell or strain, but rather an emergent property of a population of cells. We also note that the inoculum size effect on MIC that we found here – a purely stochastic phenomenon arising at low <u>absolute numbers</u> (CFU) – is distinct from the inoculum size effect already widely recognized in the literature, which is seen at high cell <u>density</u> (CFU/ml) and attributed to various density-dependent mechanisms, such as titration or enzymatic inactivation of antibiotics [35, 36, 24, 37, 32, 23, 38]. Although there are hints of the former absolute-number effect in earlier studies [25, 26], to our knowledge we are the first to provide a rigorous explanation in terms of stochastic population dynamics.

The good fit of the null model, in which every cell has an equal probability of establishment, at first seems to be at odds with the growing recognition that bacteria exhibit phenotypic heterogeneity, which could affect individual cells’ susceptibility to antibiotics [39, 40]. Indeed, it is entirely possible that the resistant cells that successfully established in our experiments were those with a particular metabolic state or gene expression level. However, this variability among cells would have no effect on our experimental outcomes: under our mathematical model (**Eqn. 1**), the probability of observing growth in a given number of replicate cultures is the same for any degree of cell-to-cell variation around a fixed mean, assuming the susceptibilities of cells within an inoculum are independent from one another (**Suppl. Text**, section 10.1). Thus, the establishment probability that we infer empirically should more accurately be interpreted as a mean among cells.

Although the role of demographic stochasticity in the fate of *de novo* mutations has long been recognized in theoretical population genetics, until very recently it had never been addressed empirically [17]. Our study joins a small handful of others that have now experimentally quantified establishment probability from single cells [41, 18, 42, 19], including two [18, 19] investigating establishment of bacterial cells in the presence of antibiotics, using different methods to ours (see **Suppl. Text**, section 10.2, for a more detailed comparison). Bacterial evolution of resistance to antibiotic treatment is also a prime example of the more general phenomenon of evolutionary rescue, whereby adaptation prevents extinction of populations facing severe environmental change [43]. Experiments quantifying the probability of rescue in initially large, but declining, populations were pioneered around a decade ago in yeast exposed to high salt concentrations [44]. More recent work clarified the quantitative relationship between initial population size and probability of rescue [45], coinciding with our **Eqn. 1** (see **Suppl. Text**, section 10.2, for further discussion). More broadly, the concepts and statistical methods developed here are applicable to a variety of situations where growth depends on success of rare cells and is thus highly stochastic, for instance the establishment of productive infection in a host following pathogen transmission [46], the onset of invasive bacterial infections [47], the outgrowth of bacteria in food products from small initial contaminants [48], or the establishment of metastases from cancerous tumours [49].

In summary, our study highlights the stochastic nature of *de novo* emergence of antibiotic resistance. In a practical sense, this stochasticity implies that to accurately assess the risk of resistance emerging, we must evaluate not only mutation rates, but also the probability that resistant mutants escape extinction when rare [9], which will depend on the antibiotic dosing regimen. Our results caution against the naïve use of the mutant selection window (MSW) for this purpose. While a positive selection coefficient is a necessary condition for resistance to outcompete an initially prevalent sensitive strain, it does not guarantee emergence when rare; indeed, we showed that single resistant cells are frequently lost at antibiotic concentrations well within the MSW. Thus, our findings suggest that moderate antibiotic doses may be more effective than previously thought at preventing *de novo* emergence of resistance, especially in infections where total pathogen density is relatively low. For antibiotics that have mutagenic effects, the chance of a resistant lineage arising in the first place might also be reduced at lower doses ([50]; see however [51]). Furthermore, use of lower antibiotic doses could reduce both adverse effects on patients and release of antibiotics into the environment [52].

## Materials and Methods

Further details of experimental protocols, data processing, mathematical models and statistical methods are provided in the **SI Appendix, Suppl. Text**.

### Bacterial strains, media, and culture conditions

#### Bacterial strains

The majority of our experiments, in streptomycin, were conducted with a set of *Pseudomonas aeruginosa* PA01 strains studied previously [32]. The streptomycin-sensitive and - resistant strains are chromosomally isogenic, while resistant strains additionally carry the clinically derived, non-conjugative plasmid Rms149 [53], which is stably maintained in PA01 at approximately two copies per cell [54]. Streptomycin resistance is conferred by the *aadA5* gene on Rms149, which codes for an enzyme that adenylates streptomycin [55]. Both plasmid carriers (resistant) and non-carriers (sensitive) are available with either YFP or DsRed chromosomal fluorescent markers or with no marker [32]. The live-dead staining experiment was conducted with the unlabelled resistant strain. All other experiments reported in the main text were conducted with the YFP-labelled resistant strain and, where applicable, the DsRed-labelled sensitive strain. We chose this pairing because YFP provides a stronger signal, facilitating detection of the resistant strain in mixed cultures. Previous work with these strains suggests that the two fluorescent labels have similar fitness effects [32], and we confirmed that the label had no substantive effect on the MIC values of the sensitive strain (**Suppl. Text**, section 2.1). For the seeding experiment in meropenem, we transformed the plasmid PAMBL2 into the same PA01-YFP background (**Suppl. Text**, section 1). This plasmid, isolated in 2007 from a patient in a Spanish hospital [27], confers meropenem resistance through three copies of the *bla*_*VIM-1*_ gene, which codes for a metallo-beta-lactamase [27, 28]. It is non-conjugative [28] and stably maintained in PA01 at an average of 2-3 copies/cell [54]. MIC values of all relevant strain-antibiotic pairs are reported in **Table S1**.

#### Media and antibiotics

We cultured bacteria in LB broth containing 5g/L NaCl (Sigma-Aldrich, product no. L3022). To assess colony-forming units, we plated on LB Agar, Vegitone, containing 5g/L NaCl and 15g/L agar (Sigma-Aldrich, product no. 19344). Streptomycin was prepared from streptomycin sulfate salt (Sigma-Aldrich, product no. S6501) and meropenem was prepared from meropenem trihydrate (Santa Cruz Biotechnology, Inc., product no. SC-485799). Stocks prepared in water were stored according to supplier directions and added to media on the day of experiments. When high antibiotic concentrations were required, stocks were instead prepared directly in LB on the day of experiments to avoid excessive dilution of the media with water. Bacterial cultures were diluted in phosphate buffered saline (PBS) prepared from tablets (Sigma-Aldrich product no. P4417). Treatment cultures were set up with 90% media plus 10% inoculating culture by volume; thus, the final concentrations of LB and antibiotics in the treatments are 90% of the prepared media values denoted on plots.

#### Culture conditions

All cultures were incubated at 37°C, shaking at 225rpm. Overnight cultures were inoculated directly from freezer stocks into 2ml of LB in 14ml culture tubes and incubated for approximately 16h. Overnight cultures were then diluted in PBS and used to inoculate treatment plates. Unless otherwise noted, experimental treatments were conducted in 200μl cultures in flat-bottom 96-well microtitre plates.

#### Scoring culture growth

In all experiments, we evaluated culture growth by measuring optical density (OD_595_) using a BioTek Synergy 2 plate reader, at room temperature. Lids on microtitre plates were briefly removed in a non-sterile environment for the reading; comparison to controls mock-inoculated with PBS indicated that contamination was rare (see **Suppl. Text** for detailed quantification in each experiment). We set a threshold of OD_595_ > 0.1 to score as growth, whereas background OD in media-only controls was typically below 0.05. Final readings at 3d post-inoculation were used for data analysis unless otherwise noted. By this time, growth had typically stabilized, with OD much higher than the threshold.

### MIC assays

Standard MIC values for all applicable strain-antibiotic pairs (i.e. resistant Rms149-carrier against streptomycin; resistant PAMBL2-carrier against meropenem; sensitive non-carrier against both antibiotics) were determined under our culture conditions using the broth microdilution method. Overnight cultures were diluted 10^3^-fold and inoculated into antibiotic-containing media at 20μl/well on 96-well test plates. This dilution factor consistently yielded an inoculation density close to 5 × 10^5^ CFU/ml, in accordance with standard protocol [10]; actual density was estimated by plating. Test plates were incubated and scored for growth at approximately 20h (as per standard protocol [10]), 2d, and 3d post-inoculation. For consistency with growth scoring in seeding experiments, the standard MIC values (MIC_S_ and MIC_R_) used to scale antibiotic concentrations on plot axes are based on results at 3d. Consensus MIC values of all tested strain-antibiotic pairs, at both 20h and 3d, are reported in **Table S1**, with results of individual replicates reported in the **Suppl. Text**, section 2.1. For the YFP-labelled Rms149-carrying resistant strain, an additional MIC assay in streptomycin was conducted varying inoculum size (**Fig. 3a**). Here, inoculations were conducted with overnight culture diluted 10^3^-, 10^4^-, 10^5^-, and 10^6^-fold (see **Suppl. Text**, section 2.2 for details).

### Seeding experiments: resistant strains in isolation

#### Experimental protocol

A highly diluted overnight culture of the YFP-labelled resistant strain (Rms149- or PAMBL2-carrier) was inoculated at 20μl/well into antibiotic-containing media on 96-well test plates. For experiments with PA01:Rms149 screening across many streptomycin concentrations (**Fig. 2**), we used three dilution factors (4 × 10^7^-, 8 × 10^7^, and 1.6 × 10^8^-fold), each to inoculate 96 replicate wells at each concentration. To test the null model of the inoculum size effect (**Fig. 4b**), we screened fewer streptomycin concentrations across a larger number of dilution factors (five in streptomycin-free conditions and six to ten in each streptomycin concentration), each with 54 replicates. These dilution factors were chosen differently for each streptomycin concentration to capture the range over which the proportion of replicate cultures showing growth increased from near 0 to near 1. For the experiment with PA01:PAMBL2 in meropenem (**Fig. 5**), we used two dilution factors (5 × 10^7^- and 2 × 10^8^-fold), each with 96 replicates per concentration. In all cases, test plates were incubated and scored for growth after approximately 1, 2, and 3d; for the null model test, incubation and readings were continued up to 5d to confirm stabilization of growth. See **Suppl. Text**, sections 4-5, for further details.

#### Model fitting

The number of replicate cultures showing growth by 3d (or, additionally, by 5d for the null model test), at each inoculating dilution factor and antibiotic concentration, was used for subsequent model fitting. To estimate single-cell establishment probability and evaluate the null model of the inoculum size effect, likelihood-based methods were used to fit a stochastic model of population growth to these data (see ***Mathematical model of establishment*** below). In addition, to evaluate the effect of antibiotic concentration on establishment, generalized linear models were fit to data from the seeding experiments screening across streptomycin (**Fig. 2**) or meropenem (**Fig. 5**) concentrations. Using the built-in R function ‘glm’, growth data were treated as binomial, with inoculating dilution factor and antibiotic concentration taken as explanatory variables, applying the complementary log-log link function (**Suppl. Text**, section 12).

### Seeding experiments: resistant strain in presence of sensitive population

Overnight culture of the DsRed-labelled PA01 sensitive strain was diluted 5-fold to obtain the “high density” inoculating culture, and (in the first experiment only) further to 500-fold to obtain the “low density” inoculating culture. Overnight culture of the YFP-labelled PA01:Rms149 resistant strain was diluted up to 5 × 10^7^-fold and 2 × 10^8^-fold. These cultures were inoculated as follows into media at various streptomycin concentrations on 96-well plates. Pure sensitive control cultures (24 replicates per test condition) were inoculated with 10μl/well of the appropriate diluted culture plus 10μl/well PBS. “Blank” wells to serve as background fluorescence controls were inoculated with 20μl/well PBS. Seeding test plates were first inoculated with 10μl/well of either PBS (for pure resistant control cultures), low-density or high-density sensitive culture. The resistant strain was inoculated at 10μl/well immediately thereafter (all sensitive and resistant culture inoculations were completed within an hour). Seeding was conducted with 30-60 replicates per test condition and resistant dilution factor (see **Suppl. Text**, section 6, for details). All test plates were then incubated as before, with optical density (OD_595_) and fluorescence (excitation: 500+/-27 nm; emission: 540+/-25 nm) measured at approximately 1, 2, and 3d post-inoculation. Among wells showing growth (OD>0.1), we considered the YFP-labelled resistant strain to have established if fluorescence exceeded 5 × 10^5^ units, chosen by comparison to pure cultures. In each test condition, the number of replicates in which resistance established was taken as data for model fitting, as in the previous seeding experiments.

### Fraction of dead cells by live-dead staining

This experiment used the PA01:Rms149 resistant strain with no fluorescent label, to avoid interfering with the signal from the stains. We inoculated streptomycin treatment cultures (six replicates per concentration) with 10^3^-fold diluted overnight culture, as in the standard MIC assay. After 7h of treatment, we diluted test cultures 100-fold and stained with thiazole orange and propidium iodide (BD Cell Viability Kit, product no. 349483). In parallel, we diluted and stained media and heat-killed cultures as controls. We sampled 50μl per diluted culture using flow cytometry (BD Accuri C6 Flow Cytometer with fast fluidics, discarding events with forward scatter FSC-H < 10000 or side scatter SSC-H < 8000). The staining and flow cytometry steps were carried out in groups containing one replicate per concentration plus controls, to avoid potentially toxic effects of stain exposure over prolonged times (**Suppl. Text**, section 7). To better discriminate cells from background in the flow cytometry data, we first gated on events according to forward and side scatter before defining clusters of dead (membrane-compromised) and intact cells based on fluorescence; see **Suppl. Text**, section 7, and **Suppl. Fig. 6** for details.

### Viable cell density dynamics

Using the YFP-labelled PA01:Rms149 strain, we tracked the number of viable cells over time in streptomycin-free media (twelve replicates per time point) and at 1/32, 1/16, 1/8, and 1/4 × MIC_R_ streptomycin (six replicates per time point). An independent test plate was used for sampling at each time point. Lower (Set A) and higher (Set B) streptomycin concentrations were split across separate plates and sampled at different times. Cultures were inoculated with 20μl of 5 × 10^5^-fold diluted overnight culture. At each sampling time, we plated 5 × 4μl spots of undiluted cultures (10% sampling by volume). The number of viable cells was estimated from total colony count following incubation. Comparison of streptomycin-free controls from both sets (A and B) indicated that the plate set effect was non-significant (ANOVA: *p* = 0.10); thus, controls were pooled for further analysis of the streptomycin effect (see **Suppl. Text**, section 8, for further details).

### Mathematical model of establishment

#### Model

We denote by *p*_*w*_ the probability that a small number of inoculated cells grows into a large population, i.e. that the culture reaches detectable OD as described above. Among a set of *n* independent replicates, the number of cultures showing growth is thus described by a Binomial(*n,p*_*w*_) distribution.

In the “null” model, similar to previous work [46, 45], a simple expression for *p*_*w*_ is derived under the assumptions that: (i) the number of cells in the inoculum is Poisson-distributed with mean 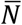; (ii) each cell, *independently*, establishes a surviving lineage with probability *p*_*c*_, which depends only on antibiotic concentration *x*; and (iii) culture growth is observed provided at least one cell establishes a surviving lineage. Then the probability of observing culture growth, as a function of mean inoculum size and antibiotic concentration, can be written as follows (**Suppl. Text**, section 10):

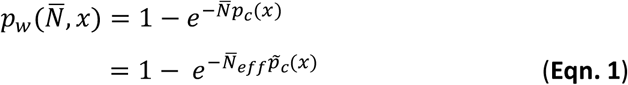

In the second line, we have rewritten the expression in terms of the “effective mean inoculum size”, 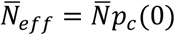, which is the mean number of established lineages in the absence of antibiotics; and the “relative establishment probability” 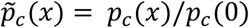. Although we expect that *p*_*c*_(0) is close to 1, 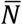 and *p*_*c*_(0) play indistinguishable roles in this model, so that in practice we can only estimate their product. This definition of effective inoculum size based on cells that grow in benign conditions is similar to the usual quantification of “viable” cells according to successful formation of a colony; we simply assess growth in liquid rather than on solid medium. Scaling up 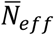 by the dilution factor applied to the inoculating culture, we have an estimate of bacterial density in this culture, equivalent to the historical “most probable number” method [56, 57]. If cells are phenotypically heterogeneous (i.e. vary in their propensity to establish), or if the individual units in the inoculum are actually clumps of cells, then *p*_*c*_ should be interpreted as the mean establishment probability among individuals (**Suppl. Text**, section 10.1).

More generally, we need not assume that cells establish independently. If we suppose simply that the number of established lineages is Poisson-distributed with some mean α (which is supported empirically by the distribution of colony-forming units counted in highly diluted cultures; **Suppl. Fig. 1**), we have the relationship

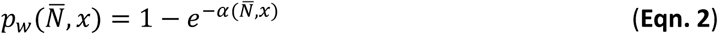

where α, and hence *p*_*w*_, have an arbitrary dependence on mean inoculum size and antibiotic concentration. In the statistical “full model”, we estimate a distinct *p*_*w*_ (or equivalently α, by the one-to-one mapping in **Eqn. 2**) in each test condition. Relative establishment probability is then generally defined by 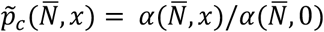. Nested models, including the null model above, make additional assumptions about the form of α (see **Suppl. Text**, section 10, for details).

#### Likelihood-based model fitting and comparisons

These stochastic models are fit to experimental population growth data using likelihood-based methods (**Suppl. Text**, section 11). Specifically, under each model we obtain a maximum likelihood estimate and a 95% confidence interval (determined by the range of parameter values that would not be rejected by a likelihood ratio test at 5% significance level) on the parameter *p*_*w*_, which can be transformed to an estimate for α. In the case of relative establishment probability, 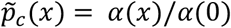, we use a profile likelihood confidence interval accounting for the uncertainty in both numerator (i.e. results at antibiotic concentration *x*) and denominator (i.e. results in antibiotic-free conditions). The fit of nested models is compared using the likelihood ratio test (LRT) at 5% significance level, i.e. a χ^2^ test on model deviance with degrees of freedom equal to the difference in number of fitted parameters between the two models.

To test the null model of the inoculum size effect, we neglect any experimental error in preparing overnight culture dilutions, and assume that mean inoculum size 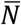 is inversely proportional to the applicable dilution factor. Effective mean inoculum size, 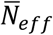, is estimated by fitting **Eqn. 1** to population growth data in antibiotic-free media. Per-cell relative establishment probability 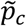 then remains as the single free parameter to fit at each tested antibiotic concentration. The goodness of fit of the null model (**Eqn. 1**) is assessed for each test concentration separately, using the LRT to compare it to the fit of the full model (**Eqn. 2**).

All model fitting was implemented in R, version 3.3.1 (The R Foundation for Statistical Computing, 2016).

## Acknowledgements

H.K.A. was supported by an Early Postdoc.Mobility Fellowship (P2EZP3_165188) and an Advanced Postdoc.Mobility fellowship (P300PA_177789) from the Swiss National Science Foundation. R.C.M. was supported by Wellcome Trust Grant 106918/Z/15/Z. We thank Isabel Frost, Natalia Kapel, Lois Ogunlana, Andrei Papkou, and Célia Souque for advice and assistance on experimental protocols.

## Author contributions

H.K.A. and R.C.M. conceived of the study and designed experiments. H.K.A. carried out experiments and data analysis with advice from R.C.M. H.K.A. developed the mathematical model, wrote the code and carried out model fitting. H.K.A. and R.C.M. wrote the manuscript.

## Data and code availability

Data generated in this study, as well as custom R scripts for likelihood-based model fitting and comparisons, will be made available upon manuscript acceptance.

## Notes

#### Summary of Updates

Two major new experiments (new Fig. 5 and Fig. 6), re-organization of materials between main and supplementary, and revised text (especially Discussion).

