## Supplementary material for "Stochastic bacterial population dynamics prevent the emergence of antibiotic resistance from single cells": SI Appendix

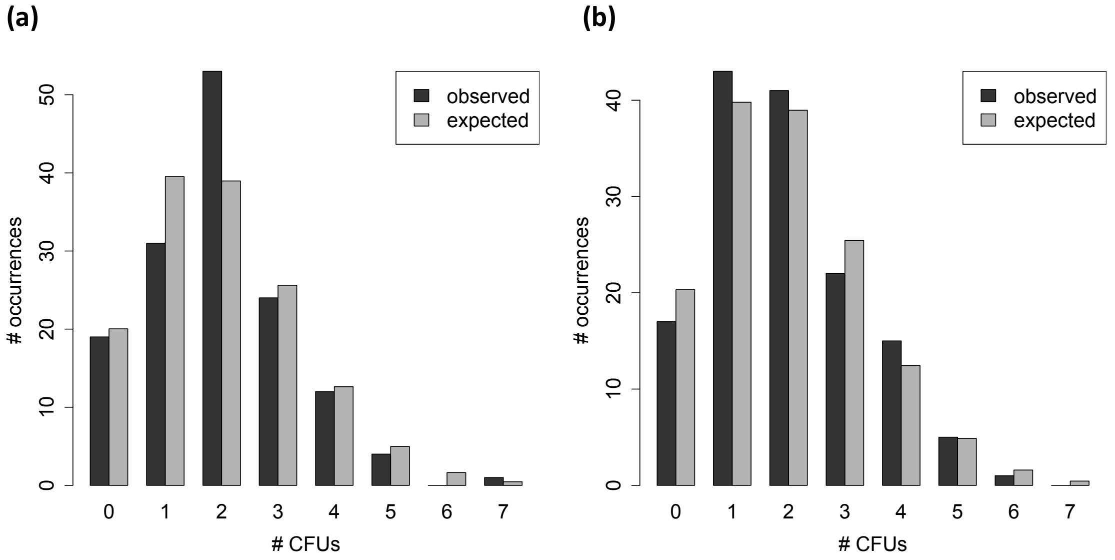


**Fig. S1. Number of colony-forming units in highly diluted culture is well described by a Poisson distribution.** In each of two separate experiments, we diluted a single overnight culture of PA01:Rms149 by $10^7$-fold and plated 144 replicate spots (4μl/spot) on LB-agar. The plots compare the observed frequency (black) of colony-forming units (CFUs) per plated spot, and the expected frequency (grey) from the best-fitting Poisson distribution. In both experiments, the Poisson distribution was not rejected under a goodness-of-fit test (categories of 0, 1, 2, 3, 4, or 5+ colonies per spot; chi-squared test with 4 degrees of freedom: $p = 0.10$ in experiment 1, panel a; $p = 0.73$ in experiment 2, panel b; see also **Suppl. Text** sections 3, 13).

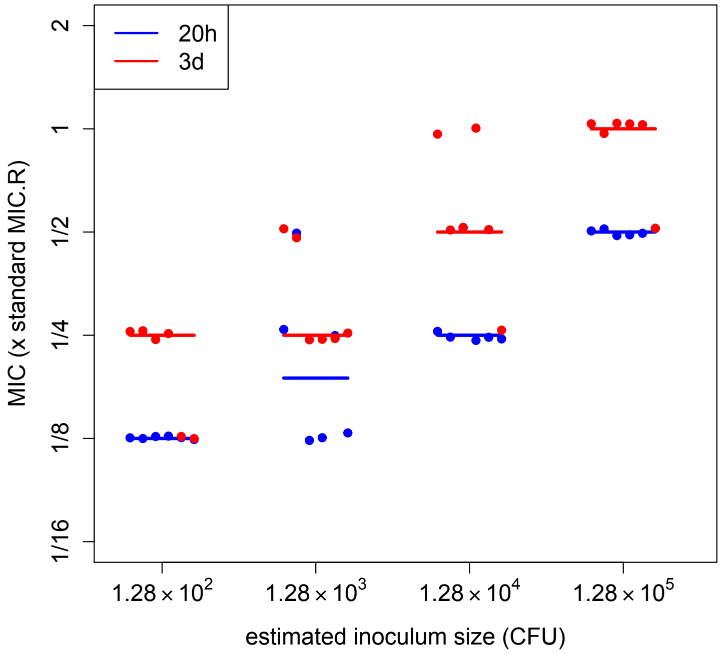


**Fig. S2. MIC of PA01:Rms149 in streptomycin, evaluated at 20h vs. 3d post-inoculation.** Test cultures at two-fold concentration steps of streptomycin were inoculated with the PA01:Rms149 strain at four different inoculum sizes. MIC was evaluated as the minimal tested concentration that prevented detectable growth up to 20h (blue) or 3d (red) post-inoculation. The data at 3d are the same as in the main Fig. 3a. The y-axis is scaled by the MIC of this strain at standard inoculation density ($MIC_R$; see **Table S1**). The points (plotted with slight offsets in the y-direction for visual clarity) represent six biologically independent replicates at each inoculum size, with the line segments indicating their median at each time point.

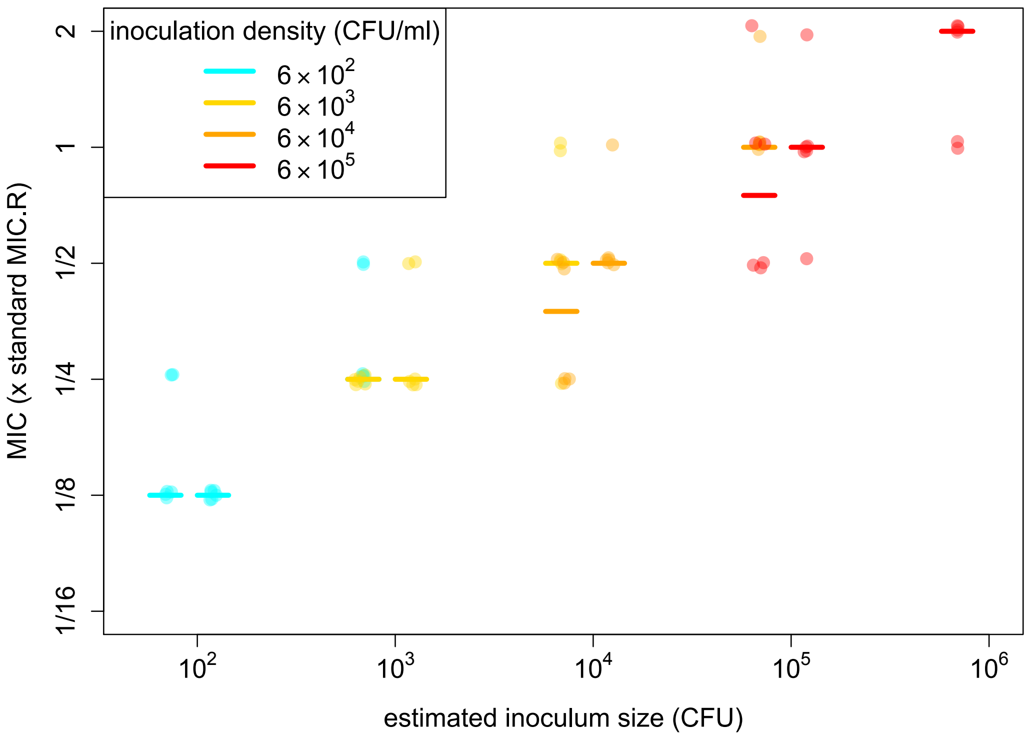


**Fig. S3. MIC of PA01:Rms149 in streptomycin as a function of inoculum size (CFU) and density (CFU/ml).** MIC, scored based on detectable growth by OD 3d post-inoculation, is scaled by the MIC of this strain at standard inoculation density ($MIC_R$; see **Table S1**). Growth was tested in two-fold concentration steps of streptomycin, up to a maximum of 1 x $MIC_R$; if growth occurred at this concentration, the MIC is plotted here as 2 x $MIC_R$ but could be higher. The absolute inoculum size in CFU, in log scale on the x-axis, was estimated by plating. The plotted points represent six biologically independent replicates at each condition and the line segments indicate their median. Points are slightly offset in both x- and y-directions for visual clarity. Inoculation density in CFU/ml is indicated by colour as per the legend. At three of the nine tested inoculum sizes (6.9 x $10^2$, 6.9 x $10^3$, and 6.9 x $10^4$ CFU), two different densities were tested in each case; note that the medians (in cyan and yellow) coincide for the two densities tested at 6.9 x $10^2$ CFU. At matched absolute inoculum sizes, in no case is there a significant effect of density on MIC: at 6.9x$10^2$ CFU, comparing 6x$10^2$ vs. 6x$10^3$ CFU/ml: $p$=0.17; at 6.9x$10^3$ CFU, comparing 6x$10^3$ vs. 6x$10^4$ CFU/ml: $p$=0.14; at 6.9x$10^4$ CFU, comparing 6x$10^4$ vs. 6x$10^5$ CFU/ml: $p$=0.21 (Wilcoxon rank-sum test with continuity correction; approximate $p$-values computed due to ties).

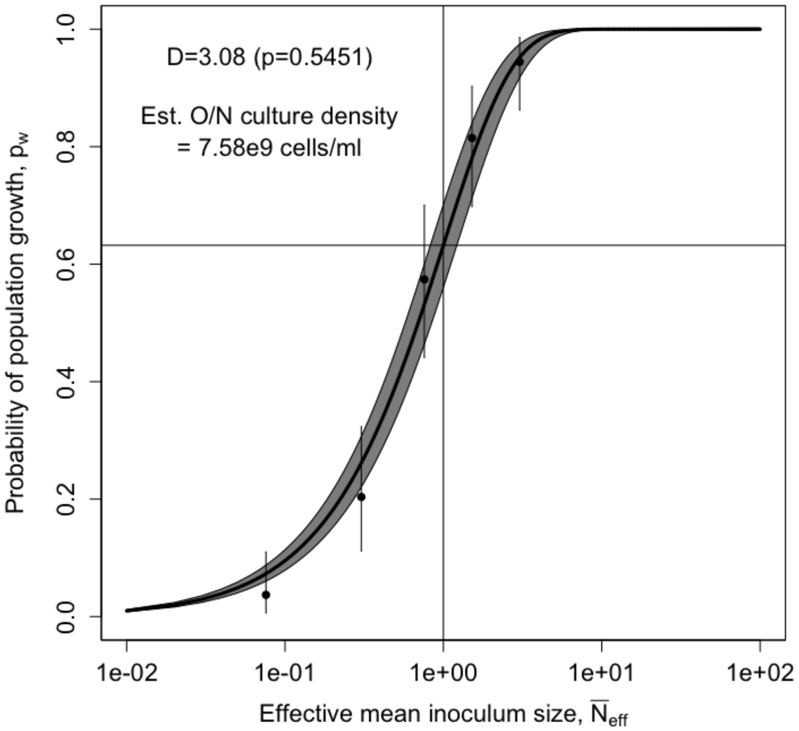


**Fig. S4. Estimating effective mean inoculum size by fitting the null model to population growth data in antibiotic-free media.** The method is illustrated here for the experimental data in the main **Fig. 3b**, testing growth of PA01:Rms149. The solid line shows the best fit of the null model (using the maximum likelihood estimate [MLE] of $\tilde{p}_c$) and the shaded area corresponds to the 95% confidence interval (CI). Points and error bars indicate the MLEs and 95% CIs in the full model, i.e. treating each inoculum size separately (here MLE simply equals the proportion of experimental replicates showing growth). Deviance (*D*) of the null model from the full model and the corresponding *p*-value from the likelihood ratio test are printed on the plot. The thin lines show the calibration of the x-axis: effective mean inoculum size ($\overline{N}_{eff}$) of 1 is defined as the point where the probability of population growth, $p_w$ = 1-exp(-1) ≈ 0.63; that is, population growth fails only in replicates receiving an effective inoculum size of zero. By scaling up this estimate by the dilution factor applied to the overnight culture for inoculations, we obtain an estimated overnight culture density of 7.58 x $10^9$ cells/ml. See **Suppl. Text,** section 11 for further details of the method and section 15 for complete results of model fitting.

**(a)** **(b)**

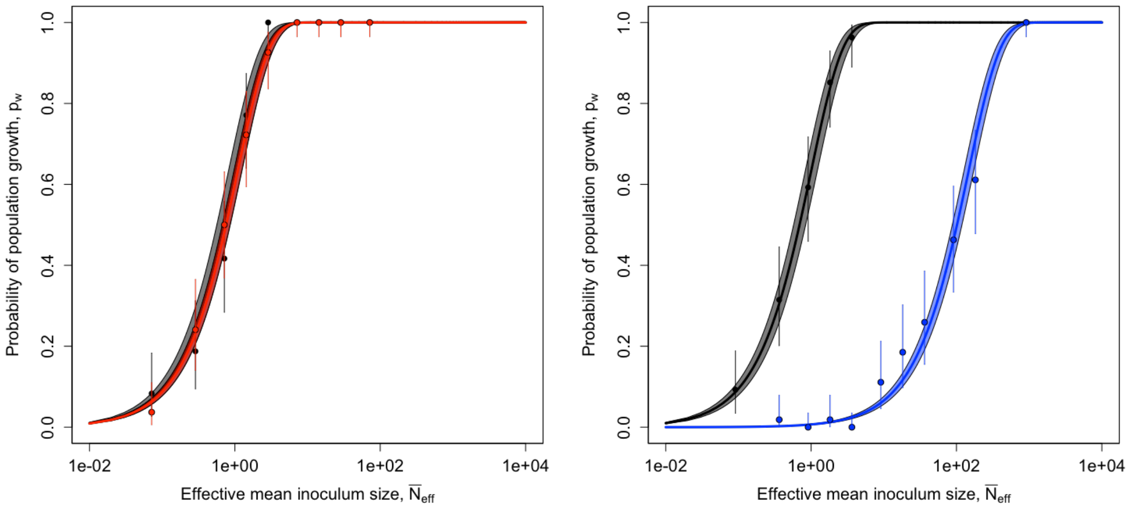


**Fig. S5: Additional tests of the null model of the inoculum size effect on population growth.** The experimental test with PA01:Rms149, similar to the main **Fig. 3b**, was repeated separately at each streptomycin concentration (panel **a**, red: 1/16 x $MIC_R$; panel **b**, blue: 1/8 x $MIC_R$) to confirm consistency of the results. The data and fits illustrated here are based on growth up to 5 days post-inoculation; see **Suppl. Text**, section 15 for comparison to results at 3 days. Effective mean inoculum size was estimated from results in streptomycin-free medium (black), tested in parallel in each experiment. The solid lines show the maximum likelihood estimate (MLE) fit of the null model, with the shaded area indicating the 95% confidence interval (CI); and the points with error bars show the MLEs with 95% CIs in the full model. In each case, the null model fits are not rejected by the likelihood ratio test (panel **a** experiment: $p$=0.073 in streptomycin-free media and $p$=1.00 at 1/16 x $MIC_R$; panel **b** experiment: $p$=0.99 in streptomycin-free media and $p$=0.16 in 1/8 x $MIC_R$). See **Suppl. Text,** section 11 for further details of the method and section 15 for complete results of model fitting.

**(a)**

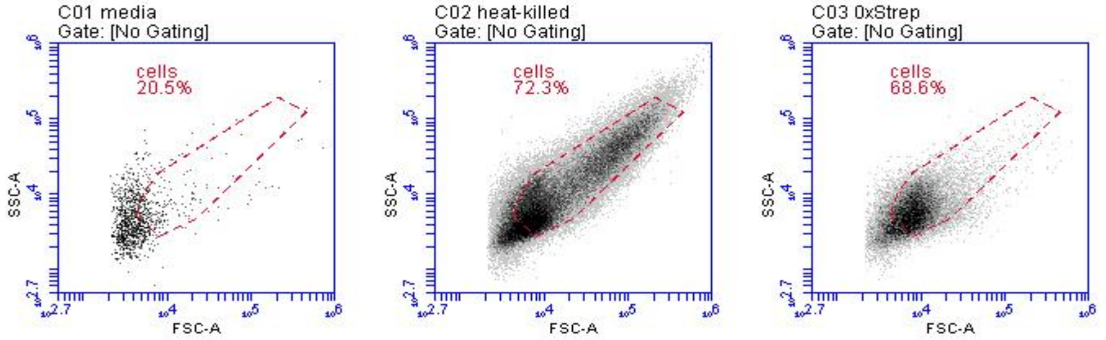


**(b)**

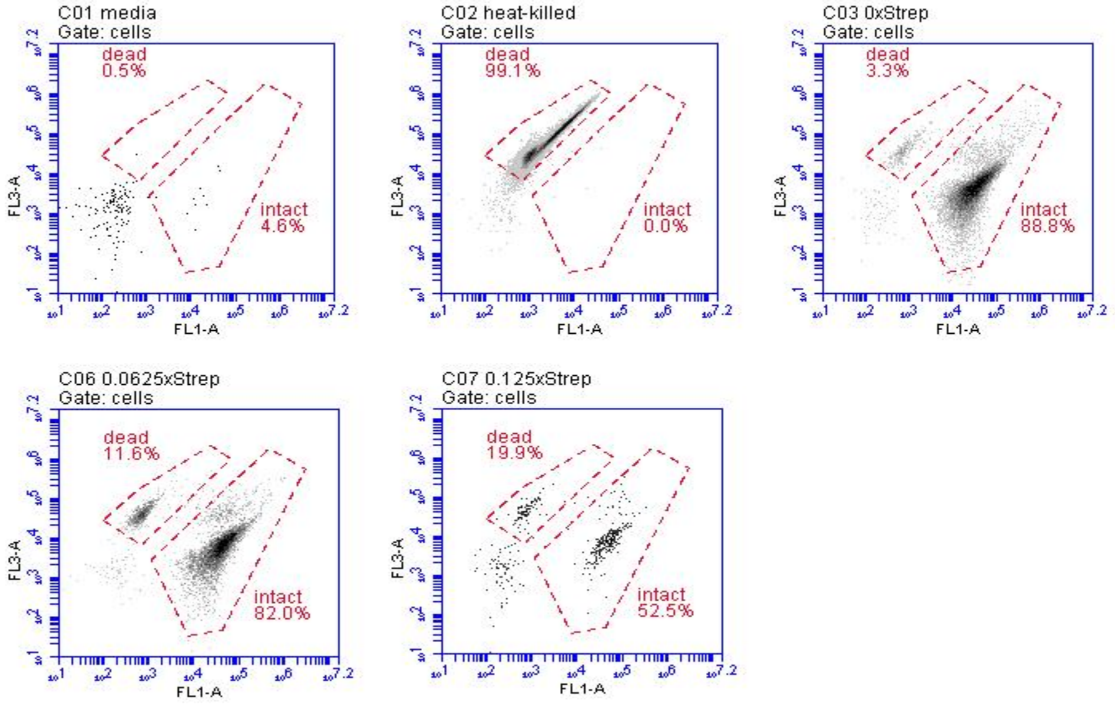


**Fig. S6: Flow cytometry with live-dead staining of PA01:Rms149 following streptomycin treatment.** PA01:Rms149 (without any fluorescent label in this experiment) was cultured in streptomycin-free media and at 1/64, 1/32, 1/16, and 1/8 x $MIC_R$ streptomycin (six replicates per concentration). After 7h, cultures were sampled, diluted and stained, along with media-only and heat-killed cell controls. **(a)** Example plots of event densities in FSC-A (forward scatter) – SSC-A (side scatter) space. In a first analysis step, the "cells" gate was drawn to better discriminate cells from background events. On average, in media-only controls, 240 events per 50µl sample fell within the "cells" gate (23% of all detected events), compared to ~40,000 events (72%) in heat-killed cell samples; ~11,000 events in streptomycin-free cultures, decreasing to ~5,300 events with 1/16 x $MIC_R$ streptomycin treatment (66-69% of all detected events); and 494 events (45%) with 1/8 x $MIC_R$ streptomycin treatment. **(b)** Example plots of event densities in FL1-A – FL3-A (fluorescence detection) space, gated on "cells" as defined above. FL1 (488nm laser with 533/30 filter) primarily detects the thiazole orange (TO) stain, while FL3 (488nm laser with 670LP filter) primarily detects the propidium iodide (PI) stain. In a second analysis step, the "dead" gate was drawn around the cluster that appeared with lower TO and strong PI staining (representing cells with compromised membranes), and the "intact" gate was drawn around the cluster that appeared with higher TO and weak PI staining (representing cells with intact membranes). This gating provided further discrimination from background events, due to the low proportion of events falling in either the dead or the intact gate in media-only controls (across six replicates, 3-17% of events within the “cells” gate, or 11-38 events per sample), compared to the high proportion in heat-killed samples (99%), cultures treated with up to 1/16 x $MIC_R$ streptomycin (86-94%), and cultures treated with 1/8 x $MIC_R$ streptomycin (67-78%). We determined the fraction of dead cells, with correction for remaining background events, as the number of events in the “dead” gate divided by the total number in both “dead” and “intact” gates (minus the numbers in each gate in the media-only control). The fraction of dead cells in the heat-killed samples was thus close to 100%, while the fraction in treated cultures varied with streptomycin concentration (**Table S3** and **Fig. 4a**).

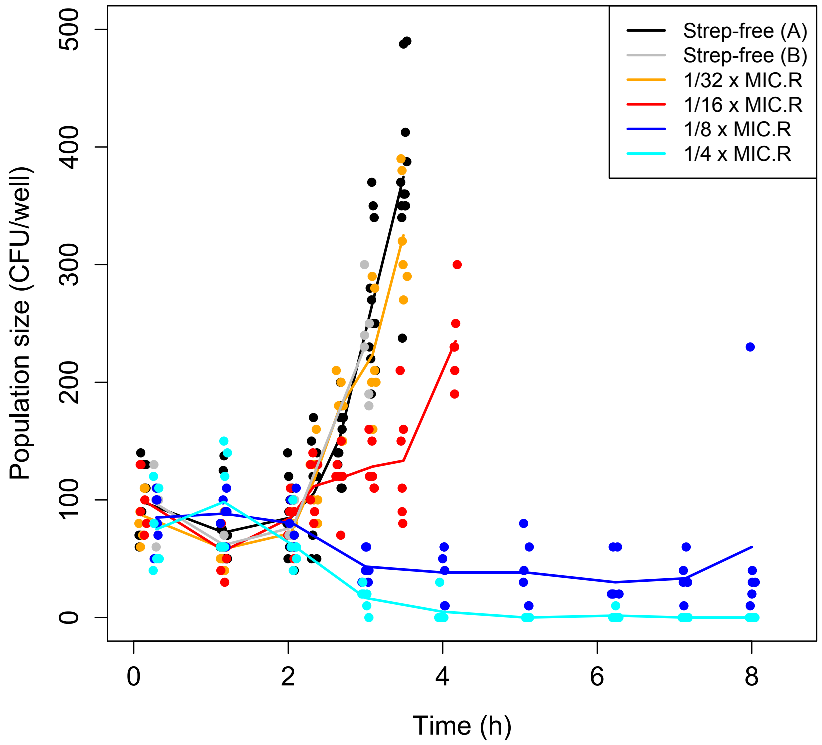


**Fig. S7: Viable cell population dynamics of PA01:Rms149 treated with sub-MIC$_R$ streptomycin.** The same data as in the main **Fig. 4b** are replotted to show individual replicates (points, plotted with slight offsets in the time axis for visual clarity), each representing an independent culture, together with their mean (connecting line) at each streptomycin concentration. There are six replicates per streptomycin concentration, per sampling time point; or twelve replicates for each streptomycin-free control.

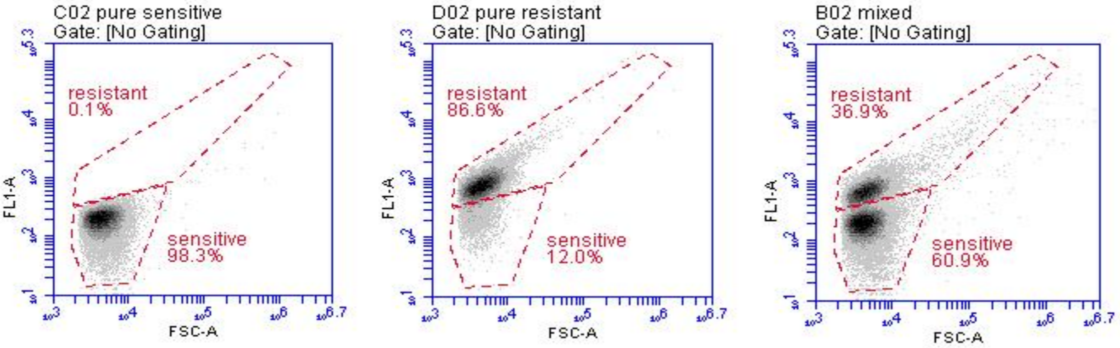


**Fig. S8: Sample plots from flow cytometry to quantify the outcome of competition assays between the PA01:Rms149 (resistant) and PA01 (sensitive) strains.** The strains are distinguished by their fluorescent labels (YFP and DsRed, respectively). These plots of event densities in FSC-A (forward scatter) – FL1-A (fluorescence detection) space illustrate the gating method. The examples shown here are samples from 500-fold diluted cultures after 24h in streptomycin-free media; from left to right: a pure sensitive strain culture, a pure resistant strain culture, and a mixed culture (inoculated with both strains in a 1:1 volumetric mixture). The FL1 detector is configured with a 488nm laser with a 533/30 interference filter, which will detect YFP fluorescence; thus, the resistant strain appears with elevated fluorescence in this channel. Note however the substantial overlap of the pure resistant culture into the “sensitive” gate; therefore, the counts falling in each gate in the mixed cultures were adjusted accordingly to infer the proportion of each strain (see **Suppl. Text**, section 9).

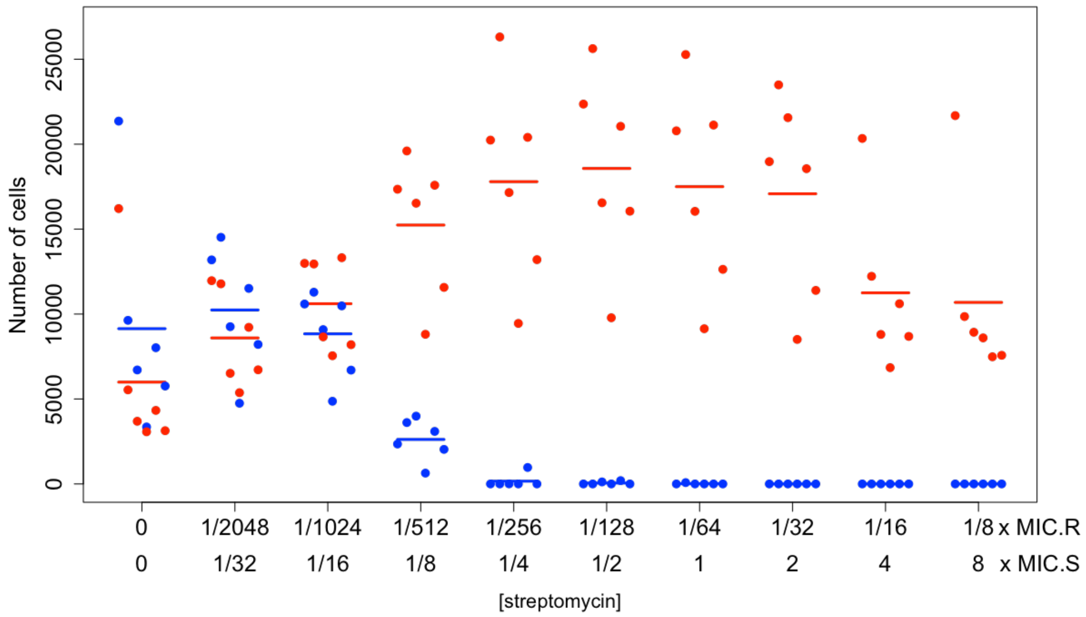


**Fig. S9: Final number of cells of each strain in competition assays between the PA01:Rms149 (resistant) and PA01 (sensitive) strains.** Following inoculation at a 1:1 volumetric mixture of both strains, cultures were incubated for 24h and then diluted 500-fold and sampled by flow cytometry. Detected events were classified as sensitive cells (blue) or resistant cells (red) according to gating by fluorescence, corrected for overlap between strains and for background events in media-only controls (**Fig. S8**). The number of cells in a 66μl sample of diluted culture is plotted here. Streptomycin concentration on the x-axis is scaled by the standard MIC values for the resistant and sensitive strains (**Table S1**). Points indicate six biologically independent replicates at each concentration, with sensitive and resistant cells in the same replicate culture plotted at the same horizontal position; line segments indicate the mean for each strain.

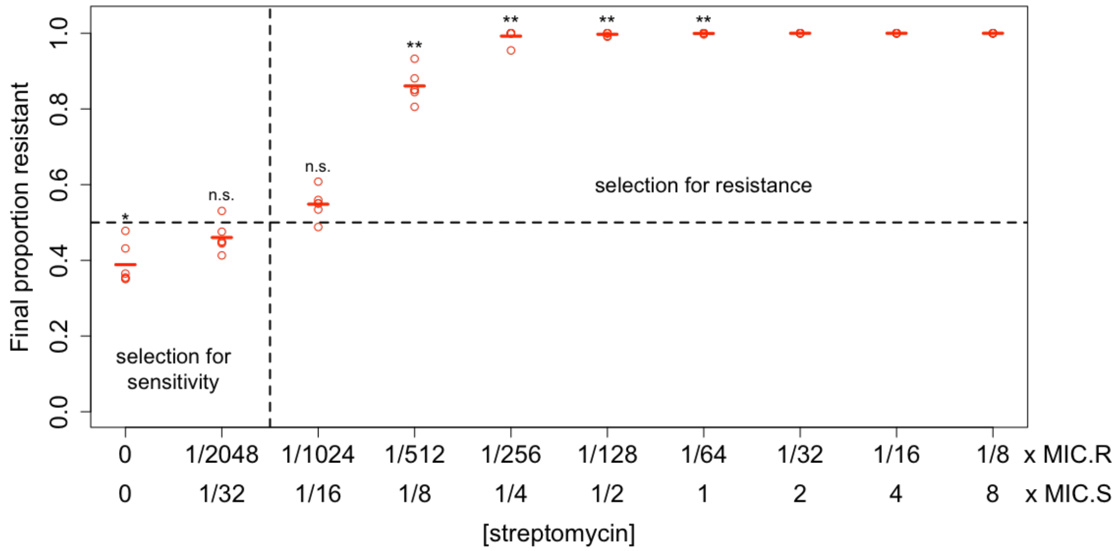


**Fig. S10: Final proportion of PA01:Rms149 (resistant) cells in competition assays with the PA01 (sensitive) strain.** Cells detected by flow cytometry after 24h of culturing were classified as resistant or sensitive, as described above (**Fig. S7-S8**), then the proportion of resistant cells was calculated within each replicate culture. The horizontal dashed line at 0.5 indicates the approximate initial proportion, which would be maintained if resistance were selectively neutral. Final proportions falling below this line indicate selection favouring the sensitive strain, while those above this line indicate selection favouring the resistant strain. The vertical dashed line indicates the approximate position of the minimum selective concentration (MSC). Streptomycin concentration on the x-axis is scaled by the standard MIC values for the resistant and sensitive strains (**Table S1**). Points represent six biologically independent replicates at each concentration, with line segments indicating their mean (see also **Table S4**). Asterisks indicate that the mean final proportion of the resistant strain significantly differs from 0.5 using a two-sided t-test at each of the lowest seven tested streptomycin concentrations, with a Bonferroni correction for multiple testing (n.s.: $p > 0.05/7$; *: $p = 4e\text{-}3$; **: $p \leq 5e\text{-}6$). At the highest three tested streptomycin concentrations, lack of variation among replicates precludes a t-test.

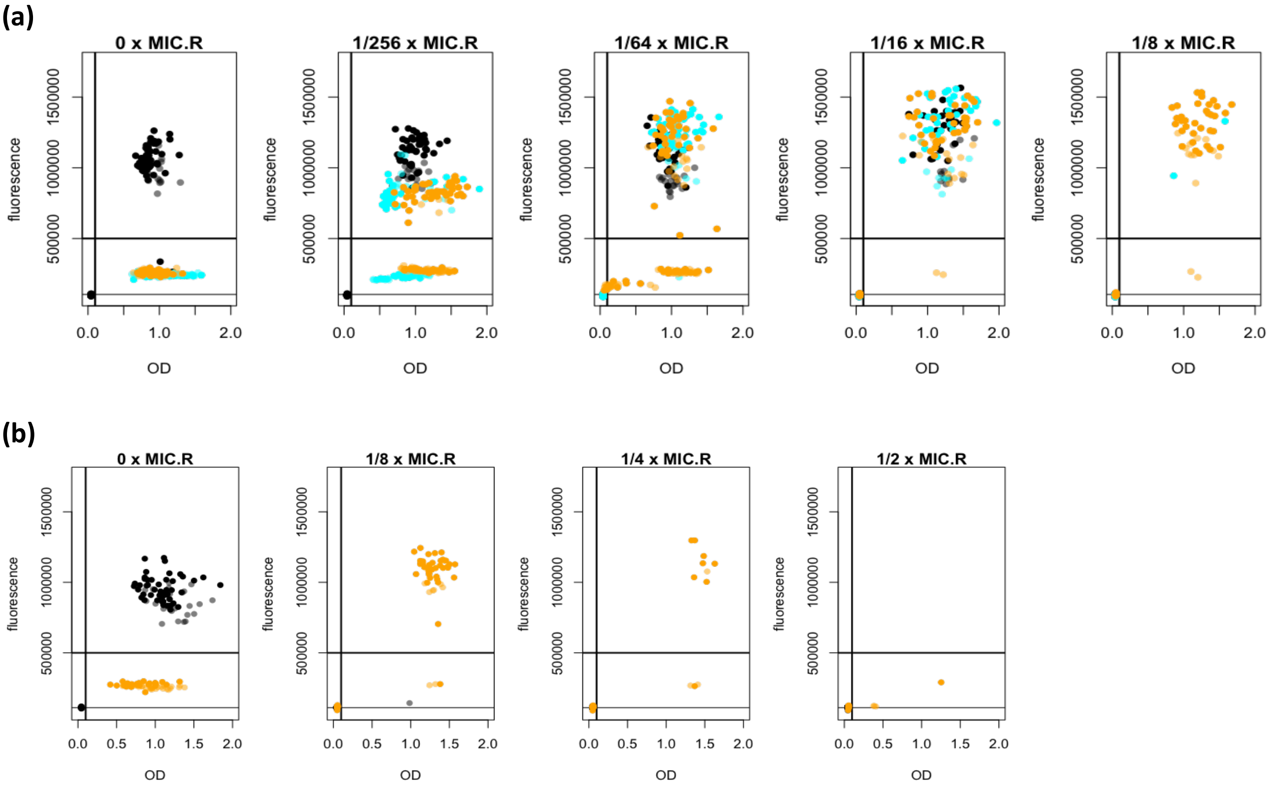


**Fig. S11: Optical density (OD) and fluorescence measured 3d post-inoculation in cultures in which PA01:Rms149 resistant cells were seeded into a PA01 sensitive cell population.** The resistant strain is labelled with YFP; elevated fluorescence (excitation: 500+/-27 nm; emission: 540+/-25 nm) was thus interpreted as establishment of the resistant strain (**Suppl. Text**, section 6). Panels **(a)** and **(b)** illustrate the results of two separate experiments that tested different sets of culture conditions, as in the main **Fig. 6**. Each point on a scatter plot represents one replicate culture. Points are colour-coded according to absence of the sensitive strain (black); presence of the sensitive strain at low density (cyan; first experiment only); or presence of the sensitive strain at high density (orange). Cultures inoculated with the resistant strain at higher mean inoculum size (5e7-fold diluted culture) are represented by points shaded darker, while those inoculated at lower mean inoculum size (2e8-fold diluted culture) are shaded lighter. Each plot corresponds to a different streptomycin concentration, as annotated above the plots. The thick lines indicate the threshold value of OD used to define growth (0.1) and the threshold value of fluorescence used to assign growth to the resistant strain (5e5). The thin lower line indicates the mean background fluorescence in media-only controls.

**Table S1. Standard MIC values of all strain–antibiotic pairs used in this study.** We used the DsRed-labelled sensitive strain and YFP-labelled resistant strains here and, unless otherwise noted, throughout the rest of our experiments. The "standard" MIC values reported here are evaluated by growth at 3d post-inoculation, from standard inoculation density of approx. 5 x $10^5$ CFU/ml. These standard values, denoted $MIC_S$ for the sensitive strain and $MIC_R$ for the relevant resistant strain, are used to scale concentrations reported throughout the manuscript. See **Suppl. Text**, section 2.1, for detailed methods.

| | **streptomycin MIC (µg/ml)** | | **meropenem MIC (µg/ml)** | |
|---|---|---|---|---|
| | **20h** | **3d** | **20h** | **3d** |
| **PA01 (sensitive)** | 16 | 32 | 0.5 | 2 |
| **PA01:Rms149 (streptomycin-resistant)** | 1024 | 2048 | N/A | |
| **PA01:PAMBL2 (meropenem-resistant)** | N/A | | 512 | 512 |

**Table S2. Estimated relative establishment probability ($\tilde{p}_c$) from likelihood-based model fits, for all seeding experiments testing PA01:Rms149 alone in streptomycin.** The fit is based on growth data at 3d or 5d post-inoculation, using the indicated model. For each experiment, the maximum likelihood estimate (MLE) of $\tilde{p}_c$, along with the lower and upper bounds of the 95% confidence intervals (CI; in parentheses), are reported. See **Suppl. Text**, section 10, for model descriptions, and sections 14.1 and 15 for complete results of model fitting.

| **[Strep] (x MIC$_R$)** | **Estimated $\tilde{p}_c$ : MLE (CI)** | | | | | |
|---|---|---|---|---|---|---|
| | **seeding expt. 1 (Model B')** | **seeding expt. 2 (Model C')** | **main inoc. size test (Model C')** | | **suppl. inoc. size tests (Model C')** | |
| | *at 3d* | *at 3d* | *at 3d* | *at 5d* | *at 3d* | *at 5d* |
| **1/64** | 0.978 (0.772,1.24) | 0.972 (0.775,1.22) | | | | |
| **1/32** | 0.925 (0.729,1.17) | 1.02 (0.811,1.28) | | | | |
| **1/16** | 0.728 (0.571,0.927) | 0.546 (0.435,0.685) | 0.909 (0.700,1.18) | 0.917 (0.706,1.19) | 0.915 (0.691,1.21) | 0.915 (0.691,1.21) |
| **1/8** | 0.0482 (0.0271,0.0795) | 0.0278 (0.0160,0.0451) | 0.0124 (0.00905, 0.0171) | 0.0165 (0.0120, 0.0226) | 0.00507 (0.00383, 0.00674) | 0.00682 (0.00516, 0.00906) |

**Table S3: Estimated fraction of dead cells in PA01:Rms149 cultures following 7h of streptomycin treatment.** The fraction of dead cells was evaluated using live-dead staining and flow cytometry. Here we report the dead fraction estimated in each individual replicate, i.e. (number of cells in dead gate)/(number of cells in dead or intact gate) after background correction (see **Fig. S6** and **Suppl. Text**, section 7, for details). At each non-zero streptomycin concentration, the difference from streptomycin-free conditions is evaluated by a one-way ANOVA with post-hoc Dunnett's test (*p*-values reported in the table; significant results in bold font). The second flow cytometry sample of the same streptomycin-free culture, taken last in each replicate, was compared to the first sample by a paired, two-sample, two-sided t-test. All analyses were repeated excluding Replicate 1, which appears as an outlier with a consistently elevated fraction of dead cells.

| | **Replicates** | | | | | | **All replicates** | | **Excluding outlier** | |
|---|---|---|---|---|---|---|---|---|---|---|
| | **1 (outlier)** | **2** | **3** | **4** | **5** | **6** | **mean** | **p-value** | **mean** | **p-value** |
| **heat-killed** | 99.99% | 100.0% | 100.0% | 100.0% | 100.0% | 100.0% | 100.0% | -- | 100.0% | -- |
| **Strep-free** | 10.48% | 3.55% | 2.42% | 1.99% | 2.73% | 3.02% | 4.03% | -- | 2.74% | -- |
| **1/64 x MIC$_R$** | 8.17% | 3.46% | 2.62% | 2.52% | 2.71% | 2.39% | 3.64% | 1.00 | 2.74% | 1.00 |
| **1/32 x MIC$_R$** | 16.35% | 7.09% | 5.61% | 5.67% | 6.77% | 6.87% | 8.06% | 0.40 | 6.40% | 0.16 |
| **1/16 x MIC$_R$** | 20.13% | 12.39% | 10.22% | 15.28% | 10.52% | 10.05% | **13.10%** | **9.3e-3** | **11.69%** | **2e-4** |
| **1/8 x MIC$_R$** | 36.25% | 28.13% | 27.09% | 19.05% | 14.80% | 18.84% | **24.03%** | **< 1e-4** | **21.58%** | **< 1e-4** |
| **Strep-free, 2$^{nd}$ sample (paired)** | 6.06% | 2.43% | 2.45% | 1.80% | 1.67% | 1.48% | 2.65% | 0.088 | 1.96% | 0.059 |

**Table S4: Estimated relative establishment probability ($\widetilde{p}_c$) from likelihood-based model fit, for the seeding experiment testing the PA01:PAMBL2 strain in meropenem.** The fit is based on growth data at 3d, using Model C' (selected by the likelihood ratio test). The maximum likelihood estimate (MLE) of $\widetilde{p}_c$, along with the lower and upper bounds of the 95% confidence intervals (CI; in parentheses), are reported. See **Suppl. Text**, section 10, for model descriptions, and section 14.2 for complete results of model fitting.

| [Mero] (x $MIC_R$) | Estimated $\widetilde{p}_c$ : MLE (CI) |
|---|---|
| **1/32** | 0.977 (0.730, 1.31) |
| **1/16** | 0.717 (0.537, 0.957) |
| **1/8** | 0.0511 (0.0305, 0.0810) |
| **1/4** | 0 (0, 0.00457) |

**Table S5: Final proportion of resistant cells in competition experiments between PA01:Rms149 (resistant) and PA01 (sensitive) strains.** The proportion of resistant cells in mixed cultures after 24h of treatment at various streptomycin concentrations (six independent replicate cultures per concentration) was determined by flow cytometry to distinguish fluorescently labelled strains (see **Fig. S7** and **Suppl. Text**, section 9, for details). The reported confidence intervals (CI) on the mean, test statistics and *p*-values are from two-sided, one-sample t-tests (d.f.=5) comparing the final proportion of the resistant strain to a mean of 0.5 (the initial proportion). Significant results after a Bonferroni correction for multiple testing (7 tests, giving a significance threshold of 0.05/7 ≈ 0.007) are in bold font. N/A: a t-test cannot be performed due to lack of variation among replicates.

| [Strep] | | Rep. 1 | Rep. 2 | Rep. 3 | Rep. 4 | Rep. 5 | Rep. 6 | mean | 95% CI on mean | test stat (*t*) | p-value (t-test) |
|---|---|---|---|---|---|---|---|---|---|---|---|
| x $MIC_R$ | x $MIC_S$ | | | | | | | | | | |
| **0** | **0** | 0.43 | 0.37 | 0.36 | 0.48 | 0.35 | 0.35 | 0.39 | (0.33, 0.44) | **-5.10** | **0.0038** |
| **1/2048** | **1/32** | 0.48 | 0.45 | 0.41 | 0.53 | 0.44 | 0.45 | 0.46 | (0.42, 0.50) | -2.44 | 0.059 |
| **1/1024** | **1/16** | 0.55 | 0.53 | 0.49 | 0.61 | 0.56 | 0.55 | 0.55 | (0.51, 0.59) | 3.06 | 0.028 |
| **1/512** | **1/8** | 0.88 | 0.84 | 0.81 | 0.93 | 0.85 | 0.85 | 0.86 | (0.82, 0.91) | **20.7** | **4.8e-6** |
| **1/256** | **1/4** | 1.00 | 1.00 | 1.00 | 1.00 | 0.96 | 1.00 | 0.99 | (0.97, 1.0) | **65.3** | **1.6e-8** |
| **1/128** | **1/2** | 1.00 | 1.00 | 0.99 | 1.00 | 0.99 | 1.00 | 1.00 | (0.99, 1.0) | **287** | **9.7e-12** |
| **1/64** | **1** | 1.00 | 1.00 | 1.00 | 1.00 | 1.00 | 1.00 | 1.00 | (1.0, 1.0) | **987** | **2.0e-14** |
| **1/32** | **2** | 1.00 | 1.00 | 1.00 | 1.00 | 1.00 | 1.00 | 1.00 | N/A | N/A | N/A |
| **1/16** | **4** | 1.00 | 1.00 | 1.00 | 1.00 | 1.00 | 1.00 | 1.00 | N/A | N/A | N/A |
| **1/8** | **8** | 1.00 | 1.00 | 1.00 | 1.00 | 1.00 | 1.00 | 1.00 | N/A | N/A | N/A |

**Table S6: Estimated relative establishment probability ($\tilde{p}_c$) from likelihood-based model fit, for the experiments seeding the resistant PA01:Rms149 strain alone or into a sensitive population, in streptomycin.** The fits are based on growth data at 3d, using Model C' (selected by the likelihood ratio test) in both experiments. The maximum likelihood estimate (MLE) of $\tilde{p}_c$ along with the lower and upper bounds of the 95% confidence intervals (CI; in parentheses) are reported. In the "baseline" condition (streptomycin-free, in the absence of the sensitive strain), $\tilde{p}_c$=1 by definition and is not estimated. See **Suppl. Text**, section 10, for model descriptions, and section 16 for complete results of model fitting.

| [Strep] | | Estimated $\tilde{p}_c$ : MLE (CI) | | | | |
|---|---|---|---|---|---|---|
| | | absence of sensitive | | sensitive at low density | sensitive at high density | |
| x $MIC_R$ | x $MIC_S$ | | | | | |
| | | Expt. 1 | Expt. 2 | Expt. 1 | Expt. 1 | Expt. 2 |
| **0** | **0** | 1, by definition | | 0 (0, 0.0206) | 0 (0, 0.0206) | 0 (0, 0.0246) |
| **1/256** | **1/4** | 1.02 (0.698, 1.50) | N/A | 1.05 (0.714, 1.53) | 0.843 (0.569, 1.25) | N/A |
| **1/64** | **1** | 0.774 (0.523, 1.14) | N/A | 0.700 (0.466, 1.05) | 0.777 (0.524, 1.15) | N/A |
| **1/16** | **4** | 0.648 (0.431, 0.968) | N/A | 0.582 (0.383, 0.878) | 0.803 (0.542, 1.19) | N/A |
| **1/8** | **8** | 0 (0, 0.0206) | 0 (0, 0.0246) | 0.0213 (0.00350, 0.0684) | 0.653 (0.435, 0.976) | 0.403 (0.271, 0.592) |
| **1/4** | **16** | N/A | 0 (0, 0.0246) | N/A | N/A | 0.0531 (0.0235, 0.104) |
| **1/2** | **32** | N/A | 0 (0, 0.0246) | N/A | N/A | 0 (0, 0.0123) |
