## Supplementary material for "Stochastic bacterial population dynamics prevent the emergence of antibiotic resistance from single cells": Suppl. Text

**Part I**

### Detailed Experimental Methods and Data Processing

#### 1 Strain construction

The PA01 strains with three fluorescent backgrounds (no marker, YFP, DsRed), with and without the Rms149 plasmid, have been used previously in our laboratory[1]. The PAMBL2 plasmid was previously obtained from clinical isolates and studied in a different PA01 genetic background[2]. For consistency, we extracted the PAMBL2 plasmid and transformed it into the same three fluorescent PA01 strains used here.

Specifically, we purified the PAMBL2 plasmid using the QIAprep Spin Miniprep Kit (QIAGEN), including steps required for large plasmids. We prepared electrocompetent cells from overnight cultures of the three fluorescently labelled PA01 recipient strains following a previously described protocol[3]. We then added 2$\mu$l of plasmid extract to 100$\mu$l of electrocompetent cells in a 2-mm gap-width cuvette and electroporated using the Bio-Rad MicroPulser (Bio-Rad Laboratories), program 'Ec-2' (single pulse, 2.5kV), then immediately added 1ml LB and incubated approximately 1 hour (37°C, 225rpm shaking), as recommended[3]. As controls, we electroporated electrocompetent cells without adding any plasmid and incubated similarly. To select transformants, we diluted cultures 10-fold and plated 100$\mu$l onto LB-agar containing streptomycin at 200$\mu$g/ml (well above the MIC of the non-plasmid-carrying PA01 strains, see SI Appendix, Table

S1; but well below the previously estimated MIC of 2048$\mu$g/ml for PAMBL2-carriers[2]). Control cultures were plated neat. After overnight incubation at 37°C, colonies appeared from all three transformed strains, but not from the controls. We picked 3 colonies per transformed strain to streak out on one streptomycin-containing plate each, incubated at 37°C overnight, then picked a single colony from each streaked plate to inoculate an overnight culture in LB containing 200$\mu$g/ml streptomycin to select for plasmid maintenance. Freezer stocks were prepared from the overnight cultures in approx. 17% glycerol and stored at -80°C.

Successful transformation of PAMBL2 into each of the three PA01 backgrounds was confirmed by PCR. Specifically, we used primers flanking the *tniA* gene on PAMBL2 (Table I.1), designed using Primer3 software[4] (courtesy of Célia Souque). The PCR reaction was carried out with GoTaq G2 Hot Start Green Master Mix (Promega), using initial denaturation (2min at 95°C); 30 cycles of denaturation (30sec at 95°C), annealing (30sec at 55°C), and extension (90sec at 72°C); and final extension (5min at 72°C). Running a gel revealed a single strong band around 500bp corresponding to *tniA* in all three strains as well as the positive control (PAMBL2 plasmid extract), but not in the negative control (water).

Table I.1: Primers used to confirm PAMBL2 transformation

| Primer | Sequence |
|---|---|
| tniA-F | GCAAGGCTCAAAAACTGCGA |
| tniA-R | CCGATGATCCGTTCCACGAT |

### 2 MIC assays

#### 2.1 Standard MIC assay (all strains/antibiotics)

We conducted standard MIC assays using the broth microdilution method. Specifically, overnight cultures were diluted $10^3$-fold, then used to inoculate antibiotic-containing media on 96-well plates at a further 1 in 10 dilution. This procedure consistently yielded final inoculation density close to $5\times10^5$ CFU/ml, as per standard guidelines[5]. Actual culture densities in each experiment were estimated by plating further-diluted cultures on LB-agar, incubating overnight at 37°C, then counting colony-forming units. Growth was tested on 96-well plates in media containing antibiotics at two-fold concentration steps, as well as in antibiotic-free media (positive controls); negative control wells were mock-inoculated with PBS. All concentrations reported here correspond to mass of the powder compound (streptomycin sulfate or meropenem trihydrate) per unit volume. Growth was assessed by optical density at 20h, 2d, and 3d post-inoculation, with

$OD_{595} > 0.1$ scored as growth. MIC in a given replicate (i.e. a row containing one culture well at each antibiotic concentration) was defined as the lowest tested antibiotic concentration preventing growth. Here, we take median MIC across replicates assessed at 3d as our standard values, to be consistent with the assessment of growth at 3d in our seeding experiments. For comparison, we also report results assessed at 20h, as per standard guidelines[5].

We first tested the sensitive strain with each fluorescent background (no marker, YFP, or DsRed) to check whether fluorescent labelling affects MIC. We used two separate overnight cultures of each strain, each tested in triplicate for growth in antibiotics (with one replicate per strain on each 96-well test plate, to control for plate effects). No contamination in the negative control wells was detected over the course of this experiment. We found no evidence that fluorescent label affects MIC (Table I.2).

Table I.2: MIC assay results: sensitive strains with each fluorescent marker

| Strain | Culture replicate[a] | Inoc. density[b] (CFU/ml) | MIC[c]: Strep. ($\mu$g/ml) | | MIC[c]: Mero. ($\mu$g/ml) | |
|---|---|---|---|---|---|---|
| | | | at 20h | at 3d | at 20h | at 3d |
| PA01 (no marker) | 1 | $4.87 \times 10^5$ | 32 | 32 | 1 | 2 |
| | 2 | $5.57 \times 10^5$ | 32 | 32 | 0.5 | 2 |
| PA01-YFP | 1 | $4.13 \times 10^5$ | 32 | 32 | 1 | 2 |
| | 2 | $4.10 \times 10^5$ | 32 | 32 | 1 | 2 |
| PA01-DsRed | 1 | $4.27 \times 10^5$ | 32 | 32 | 0.5 | $> 2$ |
| | 2 | $5.13 \times 10^5$ | 32 | 32 | 0.5 | 2 |

[a] Independent overnight cultures

[b] Inoculation density, estimated by plating

[c] Median of three growth test replicates

For consistency with strains used throughout the study, subsequent assays were conducted with the DsRed-labelled sensitive strain and the YFP-labelled resistant strains. We tested the MIC of the resistant Rms149-carrier against streptomycin; the resistant PAMBL2-carrier against meropenem; and the sensitive non-carrier against both antibiotics. For this assay we again used two overnight cultures per strain, now with six growth test replicates per culture (distributed across test plates to control for plate effects). Complete results are reported in Table I.3, with "consensus" values across the two culture replicates summarized in SI Appendix, Table S1. In the single case where the two overnight cultures yielded different MIC values (for YFP:Rms149 in streptomycin at 3d), we broke the tie by looking at individual growth replicates; this yielded an overall median of 2048$\mu$g/ml, which was consistent with the median MIC typically obtained at standard inoculation density in further experiments (section 2.2; main manuscript, Fig. 3a; and SI Appendix, Fig. S3). A single edge well (negative control) on one meropenem test plate showed apparent contamination, i.e. OD $> 0.1$, first appearing at the 3d measurement; we considered this contamination rate of $\sim 0.3\%$ across test plates to be negligible.

Table I.3: MIC assay results: all strain/antibiotic pairs used in the study

| Strain | Culture replicate[a] | Inoc. density[b] (CFU/ml) | MIC[c]: Strep. ($\mu$g/ml) at 20h | at 3d | MIC[c]: Mero. ($\mu$g/ml) at 20h | at 3d |
|---|---|---|---|---|---|---|
| DsRed | 1 | $7.45 \times 10^5$ | 16 | 32 | 0.5 | 2 |
| (sensitive) | 2 | $3.82 \times 10^5$ | 16 | 32 | 0.5 | 2 |
| YFP:Rms149 | 1 | $7.02 \times 10^5$ | 1024 | 1024 | N/A | |
| (resistant) | 2 | $4.08 \times 10^5$ | 1024 | 2048 | N/A | |
| YFP:PAMBL2 | 1 | $5.88 \times 10^5$ | N/A | | 512 | 512 |
| (resistant) | 2 | $4.02 \times 10^5$ | N/A | | 512 | 512 |

[a] Independent overnight cultures

[b] Inoculation density, estimated by plating

[c] Median of six growth test replicates

#### 2.2 MIC assay with varying inoculum size (YFP:Rms149 in streptomycin)

We extended the standard MIC assay protocol to evaluate the MIC in streptomycin of the YFP-labelled, Rms149-carrier strain at variable inoculation density. For this purpose, we used an overnight culture diluted $10^3$-, $10^4$-, $10^5$-, and $10^6$-fold to inoculate test cultures at a further 1 in 10 dilution. Thus, the highest inoculation density corresponds to the standard assay. Actual inoculum sizes were again estimated by plating. Specifically, starting from the $10^6$-fold-diluted overnight culture used for the smallest inoculum size, we took a further 5-fold dilution step, replicated three times. Each of these three $5 \times 10^6$-fold diluted cultures was plated on 5-6 LB-agar plates ($20\mu l$/plate). The number of colony-forming units was then averaged across the three replicate dilutions to obtain the reported estimate.

In the main, "standard volume" experiment (main manuscript, Fig. 3a; and SI Appendix, Fig. S2), we assessed growth in $200\mu l$ cultures on 96-well test plates, as usual. In the supplementary, "varying volumes" experiment, in order to co-vary absolute inoculum size and density (SI Appendix, Fig. S3), we additionally tested $116\mu l$ cultures on 96-well test plates and $1160\mu l$ cultures on 24-well test plates. The $1160\mu l$ volume was chosen to match surface area to volume ratio on the standard test plates. The $116\mu l$ volume was then chosen to obtain 10-fold lower volume and hence 10-fold higher density at matched absolute inoculum sizes. In both experiments, we tested streptomycin-free positive controls along with streptomycin-containing media in 2-fold concentration steps from $1/16\times\mathrm{MIC}_R$ up to $2\times\mathrm{MIC}_R$ in the standard volume experiment, or up to $1\times\mathrm{MIC}_R$ in the varying volumes experiment (in which the capacity was limited by the 24-well plates). We scored growth in six replicates per test condition.

In the standard volume experiment, the four inoculation densities were distributed across four test plates to control for any plate effects. No contamination (growth in outer, negative control wells) was detected over the course of the experiment. In the varying volume experiment, on 96-well test plates, each plate included two replicates at each inoculation density along with negative controls in outer wells; again, no contamination was detected over the course of the experiment. On 24-well test plates, each test plate included one replicate per inoculation density, while an additional plate processed in parallel served as a negative control and showed no contamination.

Occasionally, a replicate showed no growth at a given streptomycin concentration, but did grow at the next highest concentration step, before growth was abolished. (This occurred in the standard-volume experiment for one replicate at estimated inoculum size $1.28\times10^3$ CFU evaluated at 20h, and one replicate at $1.28\times10^4$ CFU evaluated at 3d.) This result can arise through stochastic effects leading to growth in some cultures but not others at any given streptomycin concentration. Note that cultures at successive streptomycin concentrations grew independently of one another, and were grouped simply by plate row as one replicate for MIC evaluation. In these ambiguous cases, we scored MIC for the replicate as the higher concentration, at and beyond which no growth occurred.

### 3 Probability distribution of the number of colony-forming units

The goal of this experiment was to test whether the number of viable cells, quantified by colony-forming units (CFUs), in a small volume of highly diluted culture can be described by a Poisson distribution. An overnight culture of the YFP:Rms149 strain was diluted $10^7$-fold in PBS and plated in a 6×8 array of $4\mu l$ spots on each of three square (12cm × 12cm) LB-agar plates, yielding a total of 144 replicate spots. These LB-agar plates were incubated for the remainder of the day of plating at 37°C, moved to room temperature overnight, then returned to 37°C in the morning until colonies were visible but still separated for counting.

#### 4 Seeding experiments to estimate establishment probability across antibiotic concentrations: resistant strain alone

This experiment was used to estimate the establishment probability of a resistant strain, either the YFP:Rms149 strain in streptomycin (main manuscript, Fig. 2) or the YFP:PAMBL2 strain in meropenem (main manuscript, Fig. 5), in monoculture. We note that a similar seeding experiment to assess population growth from an inoculum size of approximately one cell has also been used previously, on a much smaller scale, by Levin-Reisman et al. [6] (see their Supplementary Material). However, these authors did not infer a single-cell establishment probability from the proportion of populations showing growth.

**Experimental protocol:** Media was prepared at various antibiotic concentrations and dispensed column-wise onto 96-well plates at $180\mu l$/well using an automated liquid handler (BioTek Precision XS). An overnight culture of the appropriate resistant strain was diluted in PBS in a single series up until the penultimate step, then the final step was taken independently to each dilution used for inoculations. For the Rms149 strain, these dilutions were $4 \times 10^7$, $8 \times 10^7$, and $1.6 \times 10^8$-fold. For the PAMBL2 strain, these dilutions were $5 \times 10^7$ and $2 \times 10^8$-fold. These diluted cultures were used to inoculate the treatment plates at $20\mu l$/well. In parallel, negative controls were mock-inoculated with $20\mu l$/well of PBS to check for contamination, since the automated liquid handler is not maintained in a sterile environment. All plates were incubated (37°C, 225rpm) for 3d, and removed daily to measure optical density ($OD_{595}$) on a BioTek Synergy 2 plate reader with Gen5 v. 2.00 software (BioTek Instruments, Inc.). Lids were removed briefly for this reading; again, the negative controls served as checks for contamination.

In the two experiments with Rms149, each 96-well treatment plate corresponded to 96 replicates at the same streptomycin concentration and dilution factor (with negative controls on separate plates, containing antibiotic-free media, processed in parallel). Thus, it is possible that plate effects were confounded with the treatment conditions. This potential source of error was mitigated by pooling data across three dilution factors (hence three separate plates) at each streptomycin concentration in the likelihood-based model fitting (see Sections 11 and 14). Furthermore, since we obtained similar results in independent experiments (SI Appendix, Table S2), we can be confident in the effect of streptomycin, as opposed to random plate effects dominating the results. In the experiment with PAMBL2, all meropenem concentrations were tested across columns on each plate (with negative controls in outer wells), for a total of 96 replicates across plates, thus mitigating the issue of plate effects.

**Data processing:** As described in the main text Methods, wells were scored as showing growth if their optical density exceeded 0.1. The number of wells in each condition eventually (by Day 3 post-inoculation) showing growth was taken as data for model fitting (see Sections 11 and 14).

- **Rms149 experiments:** At streptomycin concentrations up to $1/16{\times}\mathrm{MIC}_R$, there were some new appearances of growth between Days 1 and 2, but no further new appearances by Day 3. Thus, at these concentrations, we are fairly confident that the growth time was sufficient to allow detection of all established populations. At $1/8{\times}\mathrm{MIC}_R$, there were still some new appearances on Day 3; thus, it is possible that we missed rare slow-growing wells that would have crossed the OD threshold even later. However, our estimates of establishment probability at $1/16{\times}\mathrm{MIC}_R$ in these experiments were similar to, and even slightly higher than, our estimates from the later inoculum size effect tests where we read OD up to 5 days post-inoculation (see Section 5 and SI Appendix, Table S2). Thus, we do not expect that undetected growth substantially affected our results.

  In experiment 2, a partially clogged dispenser needle on the automated liquid handler resulted in a reduced volume of media in row H of each plate (estimated $\sim 150\mu l$ instead of $180\mu l$) and a few of these wells dried up by Day 3. In one case, this resulted in OD that was above the threshold on Day 2 but dropped below the threshold on Day 3. This well was checked visually to confirm bacterial growth and counted as showing growth for data analysis.

  In each of the two experiments, a single well out of two 96-well negative control plates showed growth, first appearing on Day 2 or 3. This indicates a negligible contamination rate of $\sim$0.5% in antibiotic-free media.

- **PAMBL2 experiment:** Here, growth stabilized faster, with new appearances on Day 2 only at $1/8{\times}\mathrm{MIC}_R$ meropenem, and no further new appearances on Day 3. A single negative control edge well out of 32 test plates showed growth (first appearing on Day 2), indicating a negligible contamination rate of $\sim$0.2% in antibiotic-free media or $\sim$0.09% overall (across various meropenem concentrations).

### 5 Seeding experiments to test the null model of the inoculum size effect

In this experiment, we evaluated the probability of outgrowth of a detectable population of the YFP:Rms149 strain in monoculture, at a given streptomycin concentration, as a function of inoculum size (main manuscript, Fig. 3b; and SI Appendix, Fig. S4-S5). In each experiment, we also tested growth in streptomycin-free media in parallel, in order to estimate effective inoculum size in liquid culture conditions (see Sections 10-11).

**Experimental protocol:** The experiment proceeded similarly to the seeding experiments described above with the resistant strain in monoculture (Section 4). We made a single dilution series of an overnight culture of the YFP:Rms149 strain to inoculate test plates, but selected a different subset of these diluted cultures to use at different streptomycin concentrations (Table I.4). Thus, any inaccuracy in individual steps of the dilution series is a possible source of discrepancy between streptomycin-free and streptomycin-containing cultures. We did not account for this possible error in our model (which assumes perfect dilution steps), but we minimized the possibility of compounding errors in the experimental protocol by taking dilutions in parallel rather than in series wherever possible (e.g. 1e6- through 2e7-fold dilutions would each be prepared in a single independent step from a common 2e5-fold dilution). To control for possible plate effects, different inoculum sizes at a given streptomycin concentration were distributed across test plates. All test plates included edge wells as negative controls to check for contamination.

Table I.4: Testing the null model of inoculum size effect: experimental set-up

| Experiment | [Strep] ($\times$MIC$_R$) | Dilution factors inoculated[a] | replicates per dil. fac. |
|---|---|---|---|
| main | 0 | 2e9, 5e8, 2e8, 1e8, 5e7 | 54 |
| | 1/16 | 5e9, 2e9, 1e9, 5e8, 2e8, 1e8, 5e7, 2e7, 1e7 | 54 |
| | 1/8 | 5e8, 1e8, 2e7, 5e6, 1e6, 2e5 | 54 |
| suppl. 1 | 0 | 2e9, 5e8, 2e8, 1e8, 5e7 | 48[b] |
| | 1/16 | 2e9, 5e8, 2e8, 1e8, 5e7, 2e7, 1e7, 5e6, 2e6 | 54 |
| suppl. 2 | 0 | 2e9, 5e8, 2e8, 1e8, 5e7 | 54 |
| | 1/8 | 5e8, 2e8, 1e8, 5e7, 2e7, 1e7, 5e6, 2e6, 1e6, 2e5 | 54 |

[a] Dilution factors applied to the overnight culture, then inoculated at $20\mu l$ per well.

[b] Number of replicates reduced because one test plate was dropped early in the experiment.

**Data processing:** Growth of streptomycin-free cultures was scored (by $OD_{595} > 0.1$) after one day, which was determined to be sufficient for stabilization of detectable OD. (In the main and first supplementary experiments, no new appearances of growth occurred after the first day. In the second supplementary experiment, there were two new appearances of growth, but also several contaminated outer negative control wells on the second day; thus, we did not count these new appearances.)

Growth of streptomycin-treated cultures was scored daily up to five days post-inoculation. This extended protocol was chosen because in the previous seeding experiments (Section 4) we observed some new appearances of growth on the third day at the highest streptomycin concentration ($1/8\times\mathrm{MIC}_R$). In the present experiment, we observed a few cases of new growth beyond Day 3 at $1/8\times\mathrm{MIC}_R$, but only one replicate in one experiment at $1/16\times\mathrm{MIC}_R$. For comparison, we report results based on growth up to Day 3 and up to Day 5 in Table III.5 and SI Appendix, Table S2; the plots in the main manuscript, Fig. 3b, and SI Appendix, Fig. S5, are based on growth up to Day 5 as this was judged the more accurate reflection of final establishment.

Occasionally a well dried up due to evaporation by Day 5; in these cases, we counted culture growth if it appeared earlier. In supplementary experiment 2, a cluster of four wells on one plate hovered just below the threshold OD of 0.1, with one of these wells just crossing the threshold on two of the five measurements. However, there did not appear to be bacterial growth in these wells and thus they were not counted.

Contamination was generally rare, and when it did appear, it was usually late in the experiment (likely due to contamination during OD measurements with lids removed) and judged unlikely to have affected our results. There was one possible exception in supplementary experiment 2, in which there were four new appearances of growth in test cultures on Day 5, but also two cases of contamination appearing in adjacent wells on one plate. In this case, we repeated the model fitting on growth data assessed at Day 4 instead of Day 5, and still did not reject the null model ($D = 11.5$, $p = 0.24$, cf. Table III.5).

#### 6 Seeding experiments to estimate establishment probability across antibiotic concentrations: resistant strain in presence of sensitive strain

These experiments (main manuscript, Fig. 6) assessed the establishment probability of a resistant strain in the presence of a much larger initial population of the sensitive strain. We used the DsRed sensitive strain and the YFP:Rms149 resistant strain. Experiment 1 tested a lower range of streptomycin concentrations with the sensitive strain inoculated at two different densities. Experiment 2 tested a higher range of streptomycin concentrations with the sensitive strain only at higher inoculation density.

**Experimental protocol:** Overnight cultures of each strain were serially diluted in PBS. In experiment 1, where a larger volume of sensitive culture was required, two overnight cultures of the sensitive strain were first pooled and then diluted. The 5-fold diluted culture was used for "high density" inoculation in both experiments, while the 500-fold dilution was used for "low density" inoculation in experiment 1 only. Further dilution steps were plated out to estimate actual culture density (CFU/ml). The resistant strain was diluted as in the previous seeding experiments to obtain $5 \times 10^7$ and $2 \times 10^8$-fold diluted cultures for inoculation. Media was prepared at various streptomycin concentrations and dispensed column-wise onto 96-well test plates as before. Experiment 1 tested streptomycin at concentrations of 0, 1/64, 1/16, and $1/8 \times \text{MIC}_R$, with 60 replicates (distributed across plates) at each streptomycin concentration, sensitive strain density, and resistant strain dilution factor. Experiment 2 tested 0, 1/8, 1/4, and $1/2 \times \text{MIC}_R$, with 30 replicates per condition in cases where we expected little variation (streptomycin-free with high-density sensitive, and all other streptomycin concentrations in the absence of the sensitive), and 60 replicates per condition otherwise.

Seeding test plates were first inoculated with either $10\mu$l/well of PBS (absence of sensitive strain) or $10\mu$l/well of sensitive strain culture at the appropriate dilution factor, yielding initial densities of approximately $5 \times 10^5$ CFU/ml (low density) or $5 \times 10^7$ CFU/ml (high density). Then, the resistant strain culture at the appropriate dilution factor was inoculated at $10\mu$l/well. Separate plates to test growth of the sensitive strain alone (24 replicates per condition in each experiment) were inoculated with $10\mu$l/well PBS plus $10\mu$l/well sensitive strain culture. On all plates, edge wells were mock-inoculated with $20\mu$l/well PBS to serve as negative controls. In experiment 1, two additional media-only plates were inoculated with $20\mu$l/well PBS to serve as controls on background fluorescence during later plate readings. (Since fluorescence level depends on well volume and differential evaporation occurs in edge vs. interior wells, edge wells were not sufficient for this purpose.) In experiment 2, which tested fewer streptomycin concentrations, two spare interior columns on each plate served this purpose.

Test plates were incubated and read 1, 2, and 3 days post-inoculation, as before. In addition to OD, we now measured fluorescence near the peak of YFP (excitation $500\pm27$nm; emission $540\pm25$nm). For consistency across daily readings, the fluorescence 'gain' setting was fixed to 100, chosen based on pilot readings using the plate reader's 'autogain' function.

**Data processing:** As before, we set a threshold OD of 0.1 to score as culture growth. In addition, we set a threshold fluorescence of $5 \times 10^5$ units to score growth by the YFP-labelled resistant strain. This threshold was chosen such that all cultures showing growth by Day 3 that were inoculated with the resistant strain alone fell above the threshold (except for two suspected cases of cross-contamination in experiment 1; see below), while all those inoculated with the sensitive strain alone fell below the threshold. In nearly all cases, this separation was clear-cut (SI Appendix, Fig. S11). In wells scored as "growth by resistant strain", we cannot rule out that the sensitive strain is also still present; however, based on the high fluorescence level, we can be confident that a sizeable population of the resistant strain is present, and can therefore be considered as established. The number of replicate cultures scored as growth by the resistant strain was then used to estimate its establishment probability in each condition (Sections 11 and 16).

In experiment 1, in streptomycin-free media, two cultures seeded with the resistant strain alone (out of a total of 120 cultures across the two resistant dilution factors) showed growth (OD>0.1), but low fluorescence ($< 5\times10^5$). Similarly, in experiment 2, one culture in $1/8\times\text{MIC}_R$ streptomycin showed this pattern. These cases are suspected to represent cross-contamination by the sensitive strain or by other (possibly also streptomycin-resistant) bacteria in the lab. Meanwhile, growth in negative control wells also occurred at a low rate (1-2% of wells in streptomycin-free media and $1/128\times\text{MIC}_R$ streptomycin in experiment 1; 1.25% of wells only in streptomycin-free media in experiment 2), with low fluorescence readings consistent with cross-contamination by the sensitive strain or by unrelated bacteria. These contamination rates were considered negligible for the analysis.

### 7 Fraction of dead cells (live-dead staining and flow cytometry)

The goal of this experiment was to assess the proportion of dead cells induced by sub-$\text{MIC}_R$ streptomycin treatment of the Rms149 resistant strain.

**Experimental protocol:** We used the Rms149 resistant strain without any fluorescent label, so as not to interfere with the fluorescent signal from the live-dead stain. Treatment cultures on 96-well plates were inoculated with a $10^3$-fold diluted overnight culture ($20\mu l$ per $200\mu l$ total culture volume), as in the MIC tests at standard density. Media-only controls were mock-inoculated with the same volume of PBS. We tested streptomycin-free and 1/64, 1/32, 1/16, and $1/8\times\text{MIC}_R$ streptomycin with six replicate cultures per concentration. The treatment plate was incubated (37°C, 225rpm) for 7h, then cultures were immediately diluted 10-fold into sterile filtered PBS. One additional streptomycin-free culture was sampled and heat-killed (10min at 70°C) before likewise diluting 10-fold.

Live-dead staining and flow cytometry were carried out with one set of replicates at a time in order to avoid having samples exposed to the stain for too long. Each of the six replicate sets included a media-only control, a heat-killed control, and a culture treated at each tested streptomycin concentration; the streptomycin-free culture was also repeated as the last sample in order to check for an effect of time exposed to the stain before sampling. For each replicate set, the 10-fold diluted samples were diluted a further 10-fold into pre-warmed, sterile filtered 1mM EDTA in PBS and incubated for 10min at 37°C. Then $2\mu l$ each of thiazole orange [TO] and propidium iodide [PI] (BD Cell Viability Kit, product no. 349483) were added per $200\mu l$ sample and incubated 5min further at room temperature. We then analyzed $50\mu l$ per sample using flow cytometry (BD Accuri C6 Flow Cytometer with software version 1.0.264.21 – Accuri Cytometers, Inc.) with fast fluidics ($66\mu l$/min), discarding events with forward scatter FSC-H $< 10{,}000$ or side scatter SSC-H $< 8000$.

**Data processing:** To analyze the flow cytometry data, we proceeded as follows.

1. Cell densities in diluted treated cultures were sometimes low, especially at higher streptomycin concentrations. In order to better discriminate cells from background, we first defined a gate (labelled "cells") in the FSC-A/SSC-A (forward/side scatter) plot that incorporated the majority of events in the sampled cultures, but excluded the majority of events in the media-only controls (SI Appendix, Fig. S6a). Further analysis was limited to events within this gate.

2. We then defined gates around events in the FL1/FL3 plot that clustered according to their fluorescence (SI Appendix, Fig. S6b). TO (live stain) is primarily detected in the FL1 channel (488nm laser with 533/30 filter), while PI (dead stain) is primarily detected in FL3 (488nm laser with 670LP filter). Thus a cluster appearing with higher FL1 and lower FL3 was labelled "intact" and a cluster appearing with lower FL1 and higher FL3 was labelled "dead". Nearly all events in the heat-killed controls fell within the "dead" gate. Together, these two gates incorporated the majority of events in the sampled cultures, but only a minority of events in the media-only controls, providing further discrimination from background events.

3. Finally, to correct for any remaining background, within each replicate set we subtracted the number of events in the media-only control from the number of events in each sampled culture, within each of the two gates ("dead" and "intact").

4. The proportion of dead cells in sampled cultures was defined as the number of events falling within the "dead" gate divided by the total number falling in either the "dead" or the "intact" gate (after background correction as described above).

One replicate set appears as an outlier with an elevated fraction of dead cells (see SI Appendix, Table S3). Since this elevation occurs at all streptomycin concentrations, it can likely be attributed to the staining or flow cytometry steps, in which all samples were processed together, rather than the culture growth step, in which cultures at each concentration grew independently. If we exclude this replicate from the analysis, the mean fractions of dead cells correspondingly drop slightly, but the significance of streptomycin effects does not change (Table S3).

TO has toxic effects on cells, and thus the order of sampling the diluted cultures by flow cytometry within each replicate set (hence time exposed to the stain before sampling) could potentially have been confounded with the effect of streptomycin treatment. However, by comparing the streptomycin-free culture sampled earlier vs. later within each replicate set, we found that the proportion of dead cells on average actually decreased (mean of six replicates: 0.0403 in first sample vs. 0.0265 in second; SI Appendix, Table S3), though this difference is not significant (two-sided, paired t-test with 5 d.f.: $p$=0.088). This conclusion is unchanged by excluding one replicate set appearing as an outlier with consistently elevated dead fraction (two-sided, paired t-test with 4 d.f.: $p$=0.059). Thus, increases in the proportion of dead cells with streptomycin concentration can be attributed to streptomycin treatment rather than duration of staining.

### 8 Viable cell population dynamics

In this experiment, we tracked the dynamics of viable cells (CFUs) of the YFP:Rms149 strain over time, under treatment with various concentrations of streptomycin.

**Experimental protocol:** Treatment plates were split into Set A (containing 1/32 × and 1/16 × $MIC_R$ streptomycin treatments, as well as streptomycin-free controls) and Set B (1/8 × and 1/4 × $MIC_R$, and streptomycin-free controls). Importantly, we inoculated multiple treatment plates in order to sample a separate plate at each time point; thus, all replicates, at both the same and different time points, are independent of one another.

An overnight culture of the YFP:Rms149 strain was diluted $5 \times 10^5$-fold and used to inoculate treatment plates (20$\mu l$ per 200$\mu l$ total culture volume), with six independent replicate cultures at each streptomycin concentration (or twelve in streptomycin-free media) on each plate. Treatment plates were incubated at 37°C, 225rpm. In each treatment set (A and B), one plate was taken for sampling immediately after inoculation to give a time point close to 0h, in order to assess initial population size using the same method as at all later time points. Subsequent target sampling times were chosen differently for Set A (1h, 2h, 2h 20min, 2h 40min, 3h, 3h 30min, 4h) and Set B (hourly from 1h to 8h, and at approx. 24h) to account for slower growth at higher streptomycin concentrations. In total, we thus had 8 treatment plates in Set A and 10 in Set B. Actual sampling times (time elapsed between inoculation and plating) were recorded in the course of the experiment.

Upon sampling, the treated cultures were plated undiluted in $4\mu l$ spots on each of five square (12cm $\times$ 12cm) LB-agar plates, for a total sampling volume of $20\mu l$ out of each $200\mu l$ culture. After sampling, treatment plates were returned to the incubator for later OD reading (after approx. 1, 2, and 3 days) to assess eventual growth in all the sampled plates, as in our previous experiments. Contamination was rare (one negative control edge well on each of two plates showed contamination first appearing on Day 2, corresponding to an overall contamination rate of 0.3% across all plates).

LB-agar plates were immediately moved to 37°C for the rest of the day, then removed to the bench (room temperature) overnight to prevent overgrowth of colonies, then returned to 37°C the following day for several hours until colonies were visible, but still separated, for optimal counting by eye. Total colony counts from the five plates were used to estimate viable population size in the treated cultures at time of sampling (scaling up by a factor 10 from sampled to total volume). Later plated time points were excluded if colonies became too dense to count at a given streptomycin concentration.

**Data processing:** For the purpose of statistically testing the effects of streptomycin, time, and their interaction on population size (ANOVA and post-hoc Dunnett's test), we counted sampling times from Sets A and B that were within approximately 10min of each other as the same categorical sampling time. However, precise sampling times were used for plotting in the main manuscript, Fig. 4b, and SI Appendix, Fig. S7.

#### 9 Competition assay (flow cytometry)

The goal of this experiment was to determine the direction of selection for resistance vs. sensitivity across a range of streptomycin concentrations, by competing resistant (YFP:Rms149) and sensitive (DsRed) strains at reasonably high starting densities, such that demographic stochasticity is negligible.

**Experimental protocol:** We used the YFP-labelled Rms149 resistant strain and the DsRed-labelled sensitive strain. Although DsRed does not provide a strong enough fluorescent signal to aid in discrimination, it controls for the fitness effect of carrying a fluorescent marker.

Overnight cultures of each strain were mixed at an initial 20-fold dilution each, then this mixture was diluted 100-fold further. This yielded a 1:1 volumetric mixture of strains each at 2000-fold dilution, which was inoculated at $20\mu l$ per $200\mu l$ total culture volume. The total bacterial density at the start of treatment was thus expected to be around $5 \times 10^5$ CFU/ml, as we also used for MIC tests at standard inoculation density. For the pure cultures, each strain was diluted 2000-fold alone and then inoculated similarly. That is, we chose to match the density of a given strain between pure and mixed cultures, rather than the total bacterial density.

At each streptomycin concentration, we inoculated a total of 6 replicate mixed cultures and 2 replicates of each pure culture, split evenly across two treatment plates. Test concentrations ranged from 1/2048 to 1/8 $\times$ $\text{MIC}_R$ (1/32 to 8 $\times$ $\text{MIC}_S$) streptomycin in 2-fold steps, as well as streptomycin-free. Outer wells were also filled with streptomycin-free media and mock-inoculated with 20$\mu$l of PBS to serve as media-only controls. Treatment plates were incubated (37°C, 225rpm), sampled (20$\mu$l per well) at 6.5h, then immediately returned to the incubator and sampled again at approx. 24h. The latter time point provided better resolution at higher streptomycin concentrations and is thus used for data analysis. The 24h treatment culture samples (along with media-only controls) were diluted a total of 500-fold in sterile filtered PBS for flow cytometry (BD Accuri C6 Flow Cytometer). From each diluted sample, 66$\mu$l were sampled using fast fluidics, i.e. 66$\mu$l/min, discarding events with forward scatter FSC-H $< 10,000$ or side scatter SSC-H $< 8000$.

**Data processing:** To analyze the flow cytometry data, we proceeded as follows.

1. We defined non-overlapping gates 'S' and 'R' in the FL1 (fluorescence detection) – FSC-A (forward scatter) plots (see SI Appendix, Fig. S8). FL1 is configured with a blue (488nm) laser and 533/30 interference filter, which primarily detects the YFP signal. The S and R gates roughly correspond to DsRed-labelled sensitive cells (lower fluorescence) and YFP-labelled resistant cells (higher fluorescence), respectively. However, the pure cultures revealed overlap into the opposite gates, particularly of resistant cells with low fluorescence into the 'S' gate, which is accounted for in the following steps. The gates were drawn separately, but similarly, for each of the two treatment plates. In total, these gates comprised an average of 98% of all detected events on one treatment plate (range across wells: 95-99%) and an average of 96% (range: 89-98%) on the other treatment plate. Events falling outside both gates were excluded from analysis.

2. We corrected for background events in each well by subtracting the number of events in the corresponding media-only control from the number of events in the sample of interest, in each gate. If negative, we set this value to zero.

3. From the (background-adjusted) number of events in each gate in pure cultures, where we know only a single strain is present, we calculated the parameters $p_{i,j}$: the proportion of cells of strain $i$ that fall into gate $j$. For example, $p_{S,R}$ is the proportion of sensitive cells

that fall into the 'R' gate, calculated as:

$$p_{S,R} = \frac{G_R^{\text{pure } S}}{G_S^{\text{pure } S} + G_R^{\text{pure } S}}$$

where $G_j^{\text{pure } i}$ is the number of events falling into gate $j$ in the pure culture of strain $i$. (Thus, $p_{S,S} + p_{S,R} = 1$ from the pure sensitive culture and $p_{R,S} + p_{S,S} = 1$ from the pure resistant culture.) These parameters are calculated separately at each streptomycin concentration, but crucially, we assume below that they are fixed for a given strain whether it is in a mixed culture or a pure culture.

4. In mixed cultures, we want to know the "true" number of cells of each strain ($N_{\text{S,tot}}^{\text{mix}}$ and $N_{\text{R,tot}}^{\text{mix}}$), adding up cells that fall into either gate: that is, for each strain $i$, $N_{\text{i,tot}}^{\text{mix}} = N_{\text{i,S}}^{\text{mix}} + N_{\text{i,R}}^{\text{mix}}$. On the other hand, what we observe in a mixed culture is the total number of cells of either strain that fall into each gate $j$, $G_j^{\text{mix}}$. We can express the relationship between these quantities as:

$$\begin{aligned} G_S^{\text{mix}} &= p_{S,S} N_{\text{S,tot}}^{\text{mix}} + p_{R,S} N_{\text{R,tot}}^{\text{mix}} \\ G_R^{\text{mix}} &= p_{S,R} N_{\text{S,tot}}^{\text{mix}} + p_{R,R} N_{\text{R,tot}}^{\text{mix}} \end{aligned} \tag{S1}$$

where the parameters $p_{i,j}$ were calculated above from the pure cultures. With Eqn. S1 we have two linear equations in two unknowns, which can be readily solved to obtain:

$$\begin{aligned} N_{\text{S,tot}}^{(\text{mix})} &= \frac{p_{R,R} G_S^{\text{mix}} - p_{R,S} G_R^{\text{mix}}}{p_{S,S} p_{R,R} - p_{S,R} p_{R,S}} \\ N_{\text{R,tot}}^{(\text{mix})} &= G_S^{\text{mix}} + G_R^{\text{mix}} - N_{\text{S,tot}}^{(\text{mix})} \end{aligned} \tag{S2}$$

5. We typically applied Equation S2 to infer the number of cells of each strain in each mixed culture after treatment, with the proportion resistant then calculated as $N_{\text{R,tot}}^{(\text{mix})} / \left( N_{\text{S,tot}}^{(\text{mix})} + N_{\text{R,tot}}^{(\text{mix})} \right)$. A few special cases required adjustments. At the highest streptomycin concentrations, we observed no events above background counts in the pure sensitive-strain cultures; thus the parameters $p_{S,j}$ were undefined. Here we assumed that all events detected in the mixed cultures at these concentrations were resistant cells. At intermediate streptomycin concentrations, where sensitive cells were at low density but not eradicated, the general formula sometimes returned a small negative value for the number of sensitive cells. This error reflects the imperfect assumption that the proportion of cells falling in each gate is the same across cultures, whereas it will in reality show some variation. In these cases, we manually set the number of sensitive cells to zero; this adjustment had a very small effect relative to the total number of cells.

**Part II**

### Mathematical modelling and model fitting

Here we describe the models that we fit to population growth data in the seeding experiments and the tests of the inoculum size effect. We note that our approach is not entirely novel, but is presented here in full for clarity. Connections to previous work are discussed in Section 10.2, and the correspondence between different methods of fitting the data is explained in Section 12.1.

#### 10 Theoretical model of population growth

We treat the number of cultures showing growth, across independent biological replicates in a given test condition, as binomially distributed with number of trials equal to the number of replicate cultures and "success" probability $p_w$, the probability of detectable bacterial population growth. The parameter $p_w$ will depend on the inoculum size, whose expected value $\bar{N}$ is controlled by varying dilution factors of the inoculating culture; and on the environmental (media) conditions on the test plates, $x$. In our case, environment $x$ represents antibiotic concentration, but the model and methods are equally applicable to any other environmental factor(s) being tested.

The "full model" (statistically speaking) involves estimating a separate parameter $p_w$ for each $(\bar{N}, x)$ condition tested in the experiment. In this case, the maximum likelihood estimate of $p_w$ is simply the proportion of replicates that show growth (a standard result for the binomial model). The number of parameters in the full model is equal to the number of test conditions, i.e. $|\{(\bar{N}, x)\}|$.

To further relate population growth to individual cell fates, we make the fundamental assumption that population growth will be observed if and only if at least one individual in the inoculum establishes a surviving lineage. (In the following exposition, we equate "individual" with "cell", but generally an individual unit might be a clump of cells; see Section 10.1 below.) Furthermore assuming that the number of individuals that establish is Poisson-distributed with mean $\alpha$, we can express our model in its most general form as:

$$p_w(\bar{N}, x) = 1 - \mathrm{e}^{-\alpha(\bar{N}, x)} \tag{S3}$$

(corresponding to Eqn. 2 in the main manuscript). In support of this assumption, we find experimentally that the number of colony-forming units on antibiotic-free LB-agar is indeed adequately described by a Poisson distribution (SI Appendix, Fig. S1). Note that Equation

S3 defines a one-to-one mapping between $p_w$ and $\alpha$, and thus the model likelihood can be equivalently described in terms of either parameter. Thus, maximum likelihood estimates and boundaries of confidence intervals on $p_w$ can be transformed to the corresponding results on $\alpha$ using Equation S3. We refer to the full model parameterized in terms of $\{\alpha(\bar{N}, x)\}$ as **Model A** (see Fig. II.1).

It is useful to define the relative establishment probability, $\tilde{p}_c$, in a focal environment $x$, as the mean number of established cells in that environment, normalized by the result in some baseline environment (for our purposes, antibiotic-free media), denoted $x = 0$. From results at mean inoculum size $\bar{N}_i$, we calculate relative establishment probability as:

$$\tilde{p}_c^{(i)}(x) := \frac{\alpha(\bar{N}_i, x)}{\alpha(\bar{N}_i, 0)} \tag{S4}$$

Note that $\tilde{p}_c$ is not a true probability; in particular, it is possible for $\tilde{p}_c$ to exceed one, if $\alpha$ in environment $x$ exceeds that in environment 0. (The interpretation of $\tilde{p}_c$ is further discussed below.) Using definition S4, we can rewrite the full model in terms of the transformed parameters $\{\alpha(\bar{N}_i, 0), \{\tilde{p}_c^{(i)}(x_j)\}_{j=1}^{s}\}_{i=1}^{m}$, where $m = |\{\bar{N}\}|$ and $s = |\{x\}| - 1$. Note that the total number of parameters remains the same, with a one-to-one mapping between the original and transformed parameterizations. We call this rewritten version of the full model **Model A$'$**. In practice however, estimating $\{\tilde{p}_c^{(i)}(x)\}$ in the full model requires that we have tested growth in both baseline and focal environments at the same mean inoculum size $\bar{N}_i$.

Using these transformed parameters, we can also make the reasonable simplifying assumption that the relative establishment probability $\tilde{p}_c(x_j)$ is constant in a given test environment $x_j$, while $\alpha(\bar{N}_i, 0)$ varies arbitrarily with $\bar{N}_i$ (**Model B$'$**, "fixed environmental effect"). That is, we jointly estimate $\{\alpha(\bar{N}_i, 0)\}_{i=1}^{m}$ and $\{\tilde{p}_c(x_j)\}_{j=1}^{s}$ from the results pooled across all test conditions. This model still requires us to have tested growth in every environment at the same inoculum sizes. The number of parameters to be estimated is reduced from $m \cdot (s+1)$ in Model A$'$, to $m + s$ in the nested Model B$'$.

Finally, we introduce our null model of the inoculum size effect, i.e. the relationship between $p_w$ and $\bar{N}$. Here we invoke the key assumption that each individual acts independently, i.e. the outcome of establishing a surviving lineage is not affected by other individuals in the inoculum. (This independence assumption is very common when modelling dynamics at low population density, for instance using branching processes; here we rigorously test the validity of this assumption.) Suppose that the number of individuals in the inoculum is Poisson-distributed with mean $\bar{N}$ and that the fate of each individual (i.e. whether it establishes a surviving lineage) is an independent Bernoulli trial with success probability $p_c$, called the per-cell establishment probability, which depends only on the environment, $x$. Then we arrive at the number of established lineages being Poisson-distributed,* consistent with our earlier assumption, and can write the mean very simply as:

$$\alpha(\bar{N}, x) = \bar{N} \cdot p_c(x) \tag{S5}$$

Substituting Eqn. S5 into Eqn. S3 leads to

$$p_w(\bar{N}, x) = 1 - \mathrm{e}^{-\bar{N} p_c(x)} \tag{S6}$$

(Eqn. 1 in the main manuscript), which we call the null model of the inoculum size effect, relating $p_w$ to $\bar{N}$. Under this model, inoculum size cancels out in the definition of relative establishment probability (Eqn. S4) and we have simply:

$$\tilde{p}_c(x) = \frac{p_c(x)}{p_c(0)} \tag{S7}$$

Note that according to this model, we cannot obtain estimates of absolute establishment probability ($p_c$) since this parameter plays a symmetrical role to inoculum size ($\bar{N}$) and thus their effects cannot be separated. That is, if we observe a higher proportion of established populations in an experiment, we cannot tell whether this was due to higher inoculum size or higher establishment probability. This limitation is not unique to our experimental protocol: quantifying cell density by counting colony-forming units on solid media relies on the implicit assumption that the establishment probability of a "viable cell" is one. Likewise, we will refer to the mean number of established cells in the baseline (here, antibiotic-free) environment, i.e. $\alpha(\bar{N}, 0) = \bar{N} p_c(0)$, as the effective mean inoculum size, and estimate establishment probabilities in all other environments relative to this baseline. The relative establishment probability in environment $x$, $\tilde{p}_c(x)$, generally provides an upper bound on the absolute establishment probability $p_c(x)$, and if $p_c(0)$ is close to one, as we expect in benign conditions, then $\tilde{p}_c(x)$ will be close to $p_c(x)$.

* The number of cells in the inoculum, $N$, is Poisson-distributed with mean $\bar{N}$ and thus has probability generating function (PGF) $g_N(z) = \mathrm{e}^{-\bar{N}(1-z)}$. Each cell independently has probability $p_c$ of establishing a surviving lineage. Letting $Y$ denote the number of established lineages, $Y|N$ is the sum of $N$ independent Bernoulli($p_c$) trials, and thus has a Binomial($N, p_c$) distribution with PGF $g_{Y|N}(z) = (1 - p_c + p_c z)^N$. We can then derive the distribution of $Y$ via its PGF, $g_Y(z)$, as follows:

$$\begin{aligned} g_Y(z) := \mathbb{E}\left[z^Y\right] &= \mathbb{E}_N\left[\mathbb{E}\left[z^Y|N\right]\right] \\ &= \mathbb{E}_N\left[(1 - p_c + p_c z)^N\right] \\ &= g_N(1 - p_c + p_c z) \\ &= \mathrm{e}^{-\bar{N} p_c (1-z)} \end{aligned}$$

This is the PGF of a Poisson random variable with mean $\bar{N} p_c$.

For the purposes of parameter estimation under the null model, we pool results across inoculum sizes by supposing that we do not make any experimental error in culture dilution steps, and so the mean inoculum size is inversely proportional to the dilution factor applied to the inoculating culture. That is, the $i^{\text{th}}$ inoculum size is:

$$\bar{N}_i = \bar{N}^*/(d_i/d^*)$$

and thus

$$\alpha(\bar{N}_i, x) = \alpha(\bar{N}^*, x)/(d_i/d^*)$$

where $\bar{N}^*$ is the mean inoculum size at a chosen normalizing dilution factor $d^*$, and $d_i$ is the $i^{\text{th}}$ dilution factor. This assumption can be applied either to the parameterization in terms of $\{\alpha(\bar{N}^*, x)\}$ (giving **Model C**) or the transformed parameterization in terms of $\alpha(\bar{N}^*, 0)$ and $\{\tilde{p}_c(x)\}$ (giving **Model C′**), resulting either way in $|\{x\}|$ parameters to estimate. Since we have now defined a scaling relationship between inoculum sizes, we are no longer constrained to using the same set of inoculum sizes in every environment in order to estimate $\tilde{p}_c$. Note that any deviations of the data from this model fit could reflect not only lack of independence among individuals (the null model assumption), but also experimental errors in the dilution steps.

##### 10.1 Heterogeneous establishment probability

In the above model, we assumed that establishment probability $p_c$ is the same for every individual. Realistically, however, even genetically identical cells are likely to be in variable physiological states (metabolism, gene expression levels, phases of the cell cycle, etc.) that could affect their division and/or death rates. Furthermore, bacterial cells may aggregate, such that the individual units are actually clumps containing variable numbers of cells. (For *P. aeruginosa*, it appears that even in planktonic cultures, the majority of cells are in aggregates during exponential growth phase, and a minority remain in aggregates during stationary phase[7].) Both of these issues imply that the establishment probability should vary among individual units. Here we show mathematically that these issues do not affect the formulation of the null model (Model C), but simply require the appropriate interpretation of the parameter $p_c$.

The key requirements of this model are that the number of individual units in the inoculum – regardless of whether these are single cells or aggregates – is Poisson-distributed, and that each individual unit then behaves independently. Suppose an individual establishes with probability $P$, now treated as a random variable drawn <u>independently</u> for each individual from an arbitrary distribution with probability density function $f_P$. For instance, the distribution of $P$ could reflect differences in phenotype (e.g. level of antibiotic resistance) based on intracellular state,

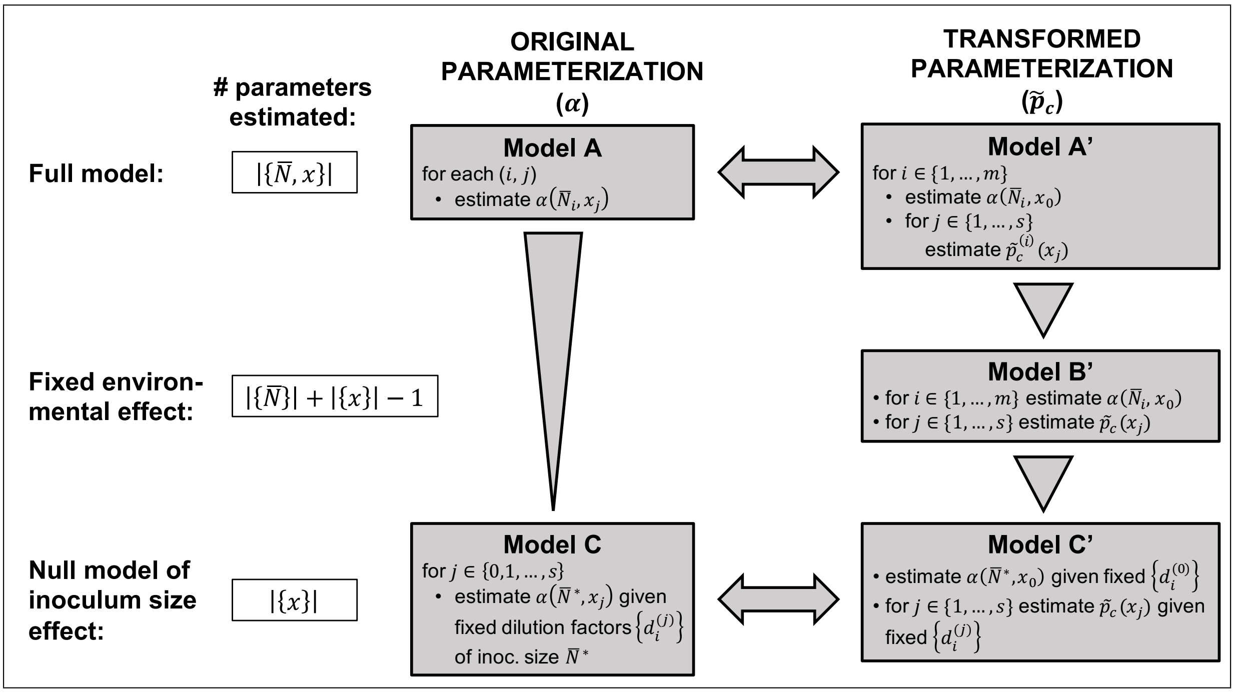


Figure II.1: Flow chart summarizing the relationships among models fit to population growth data across multiple inoculum sizes (indexed by $i$) and environmental conditions (indexed by $j$). For short, we write $m$ for the number of inoculum sizes tested, i.e. $m = |\{\bar{N}\}|$, and $s$ for the number of non-baseline environments, i.e. $s = |\{x\}| - 1$. The baseline environment is $x_0 = 0$. Mathematical equivalence of two models is denoted by a double arrow, while a nesting relationship is indicated by a triangle pointing towards the nested model. Note that fitting Model A$'$ or B$'$ requires that we have tested every inoculum size $\times$ environment combination, while the other models allow different inoculum sizes to be tested in different environments.

or variation in cell aggregate size (with larger aggregates expected to have higher establishment probability).

Let $Y$ be the indicator random variable for the event that an individual unit establishes a surviving lineage. (In the case that the individual units are aggregates of cells, establishment could be due to one or more cells.) Thus $Y|P \sim \text{Bernoulli}(P)$. The unconditional distribution of $Y$ can then be derived as:

$$\begin{aligned}\Pr(Y = 1) &= \int f_P(p) \Pr(Y = 1|P = p)\, \mathrm{d}p \\ &= \int f_P(p)\, p\, \mathrm{d}p \\ &\equiv \mathbb{E}[P]\end{aligned}$$

Thus, $Y$ is overall a Bernoulli trial with success probability equal to the mean establishment probability, and in turn, the sum of $N$ such independent trials yields a binomial distribution. Therefore, the null model outlined above still applies if we simply interpret $p_c$ as the mean establishment probability among individual units behaving independently of one another. In particular, under this model, experimental parameters (e.g. the growth phase from which the inoculating culture was taken) will only affect the outcome if they change the mean establishment probability, whereas the extent of individual variation about a fixed mean does not matter.

##### 10.2 Comparison to previous work

The same or similar theoretical models as presented here have also been used previously. Much earlier, Druett[8] derived the equation (converted to our notation)

$$p_w = (1 - p_c)^N \tag{S8}$$

for the probability of population growth from a fixed inoculum size $N$, assuming independence among individuals. Druett used this equation to model establishment of infection by bacteria invading a host. Meanwhile, our null model of inoculum size effect,

$$p_w = 1 - \mathrm{e}^{-\bar{N} p_c} \tag{S9}$$

arises very generally under the assumptions that the number of individuals is Poisson-distributed (with mean $\bar{N}$) and the fate of each individual is independent[9]. Martin et al.[9] used this general equation to describe the probability of evolutionary rescue[†] in a variety of scenarios (their Equation 3.1). In this case, $p_c$ (or $\theta^*_R$ in their notation) represents "rate of rescue per inoculated

[†]Evolutionary rescue is the situation in which a population, in decline due to a change in environment, recovers due to outgrowth of rare well-adapted genetic variants.

individual" (ref.[9], p. 4), which captures different processes depending on the scenario. For instance, in the case of rescue relying on *de novo* mutations, $\theta_R^*$ accounts for the mutation rate as well as the establishment probability of mutants. In the case of rescue relying on pre-existing (but rare) mutations, $\theta_R^*$ is simply the per-individual establishment probability, as in our interpretation.

Importantly, Martin et al.[9] also fit this equation to experimental data using similar methods to ours (cf. Sections 11-12 below). They likewise assessed the goodness of fit of this model (S9) according to deviance from the full model (our Model C vs. A), but did not include our additional Model B′. Our approach moreover differs in that we estimate effective inoculum size from growth in baseline (antibiotic-free) conditions in parallel with test conditions, rather than treating inoculum size as an independently known value. We thus derive confidence intervals on the estimated establishment probability relative to the baseline conditions, taking into account the uncertainty in both measures.

We are aware of two other empirical studies that, like us, have estimated the establishment probability of single bacterial cells faced with antibiotics, but using different methods.[10;11] These two studies quantified bacterial growth on solid media, and could thus directly visualize the number of established cells (reflecting $\alpha$, in our notation) as colony-forming units. The ratio between CFU counts at any given antibiotic concentration and CFU counts on antibiotic-free plates, sometimes called "plating efficiency",[11] is equivalent to our "relative establishment probability" ($\tilde{p}_c$). In contrast, with liquid cultures, we visualize growth of populations from a random number of cells (reflecting $p_w$), with growth scored as a binary outcome. We then infer the mean number of established cells that yielded this growth ($\alpha$), and hence relative establishment probability, using our stochastic model (Eqn. S3 and S4). One strength of our method is that experiments in liquid culture can readily be conducted on a large scale and are amenable to automation. We are aware of one other study[6] using seeding experiments to assess growth from very few cells in liquid culture; however, this study only assessed the proportion of replicate cultures showing growth ($p_w$), and did not fit these data to a model to estimate the per-cell establishment probability.

#### 11 Likelihood-based parameter estimation and model comparison

**Basic binomial likelihood:** The number of cultures showing growth at a given test condition, $n_{\text{grow}}$, is modelled as binomially distributed with number of trials equal to the total number of replicate cultures, $n_{\text{tot}}$, and success probability $p_w$, the parameter we want to estimate. That is,

$$\Pr(n_{\text{grow}}|(n_{\text{tot}}, p_w)) = \binom{n_{\text{tot}}}{n_{\text{grow}}} p_w^{n_{\text{grow}}} (1-p_w)^{n_{\text{tot}}-n_{\text{grow}}}$$

and so the log likelihood function of $p_w$ given the data $(n_{\text{grow}}, n_{\text{tot}})$ can be written (up to a constant that can be dropped) as:

$$\log \mathcal{L}(p_w|(n_{\text{grow}}, n_{\text{tot}})) = \begin{cases} n_{\text{grow}} \log(p_w) + (n_{\text{tot}} - n_{\text{grow}}) \log(1-p_w)\,, & 0 < n_{\text{grow}} < n_{\text{tot}} \\ n_{\text{tot}} \log(1-p_w)\,, & n_{\text{grow}} = 0 \\ n_{\text{tot}} \log(p_w)\,, & n_{\text{grow}} = n_{\text{tot}} \end{cases} \tag{S10}$$

We have the simple analytical result for the maximum likelihood estimate (MLE):

$$\hat{p}_w = n_{\text{grow}}/n_{\text{tot}}$$

To obtain likelihood-based confidence intervals, we use the test statistic:

$$D(p_w) = 2(\log \mathcal{L}(\hat{p}_w) - \log \mathcal{L}(p_w))$$

i.e. twice the difference in log likelihood between the MLE and any test value of $p_w$, and solve for the boundaries $p_w^*$ such that $D(p_w^*) = D^*$, the critical value for a chosen significance level in the chi-squared test with one degree of freedom (ref.[12], §2.6-2.9 and §9.5). The MLE and confidence interval boundaries for $p_w$ can simply be transformed to those for $\alpha$ using:

$$\alpha = -\log(1-p_w)$$

**Pooling dilution factors:** Recall that under the null model of inoculum size effect, and assuming perfect dilution steps, we have

$$\alpha(\bar{N}_i, x) = \alpha(\bar{N}^*, x)/(d_i/d^*) \tag{S11}$$

where $d_i$ is the $i^{\text{th}}$ dilution factor, normalized by a chosen dilution factor $d^*$, taken as known values. This leaves a single parameter $\alpha^*(x) := \alpha(\bar{N}^*, x)$, the mean number of established cells scaled to the chosen dilution factor, to be estimated in each environment by pooling data across all dilution factors. We thus define the pooled likelihood function in environment $x$:

$$\log \mathcal{L}_{\text{pool}}\left(\alpha^*(x)|\{(n_{\text{grow}}, n_{\text{tot}})\}_{i=1}^m\right) = \sum_{i=1}^m \log \mathcal{L}\left(1 - \mathrm{e}^{-\alpha^*(x)/(d_i/d^*)}|(n_{\text{grow}}, n_{\text{tot}})_i\right) \tag{S12}$$

where $\log \mathcal{L}$ is the binomial log likelihood defined in Equation S10, and $m$ is the number of dilution factors tested in this environment.

**Working with relative establishment probability:** Recall that we can transform the model from a parameterization in terms of $\{\alpha(\bar{N}, x)\}$ in all environments $x$, to one in terms of $\alpha(\bar{N}, 0)$ in the baseline environment ($x = 0$) along with $\{\tilde{p}_c(x)\}$ in environments $x \neq 0$. To obtain MLEs on $\tilde{p}_c$, we could simply substitute MLEs from the original parameterization in terms of $\alpha$: $\hat{\tilde{p}}_c(x) = \hat{\alpha}(x)/\hat{\alpha}(0)$. However, the confidence intervals must take into account the uncertainty in both the numerator and denominator. This requires running the inference jointly across multiple environments.

We write the joint log likelihood across all tested conditions (inoculum sizes and environments), given the data $(n_{\text{grow}}, n_{\text{tot}})$ in each condition, as:

$$\begin{aligned} &\log \mathcal{L}_{\text{joint}} \left( \{\alpha(\bar{N}_i, 0)\}_{i=1}^{m_0}, \{\{\tilde{p}_c^{(i)}(x_j)\}_{i=1}^{m_j}\}_{j=1}^{s}\} | \{(n_{\text{grow}}, n_{\text{tot}})\} \right) \\ &= \sum_{i=1}^{m_0} \log \mathcal{L}(1 - \mathrm{e}^{-\alpha(\bar{N}_i, 0)}) + \sum_{j=1}^{s} \sum_{i=1}^{m_j} \log \mathcal{L} \left( 1 - \mathrm{e}^{-\alpha(\bar{N}_i, 0) \tilde{p}_c^{(i)}(x_j)} \right) \end{aligned} \tag{S13}$$

where $m_j$ is the number of inoculum sizes tested in environment $x_j$. Proceeding further depends on the particular model (cf. Fig. II.1):

- Model A′ (full model): We estimate a distinct value of $\alpha(\bar{N}_i, 0)$ and $\tilde{p}_c^{(i)}(x_j)$ for each inoculum size $\bar{N}_i$, requiring that the same inoculum sizes are tested in each environment. For any given value of $\alpha(\bar{N}_i, 0)$, we could simply rescale all $\tilde{p}_c^{(i)}(x_j)$ to obtain the same optimal likelihood. Therefore the estimate of $\alpha(\bar{N}_i, 0)$ can be computed for the single condition $(\bar{N}_i, 0)$ in isolation. However, estimating each $\tilde{p}_c^{(i)}(x_j)$ requires simultaneous consideration of the baseline environment (0) and the focal environment $(x_j)$ at the $i^{\text{th}}$ inoculum size, implying that the same inoculum size must have been tested in both environments.

- Model B′ (fixed environmental effect): We estimate $\alpha(\bar{N}_i, 0)$ for each $i$, but assume that $\tilde{p}_c(x_j)$ is the same for every $i$. Adjusting the value of $\tilde{p}_c(x_{j^*})$ in any single environment $x_{j^*}$ would affect the optimized values of $\alpha(\bar{N}_i, x_0)$ for all $i$, which in turn affects the optimized values of all other $\tilde{p}_c(x_j)$. Thus, we must conduct joint inference on all parameters using the entire data set (i.e. all inoculum sizes and environments) simultaneously. Again, this requires the same inoculum sizes to have been tested in each environment.

- Model C′ (null model of inoculum size effect): We estimate a single parameter $\alpha_0^* := \alpha(\bar{N}^*, 0)$ in the baseline environment at some normalizing dilution factor $d^*$, assuming Eqn. S11 holds; as well as $\tilde{p}_c(x_j)$ in each non-baseline environment. We can again estimate $\alpha_0^*$ for the baseline environment in isolation, and $\tilde{p}_c(x_j)$ for each environment $x_j$ using only the data from the baseline and focal ($j^{\text{th}}$) environments. Therefore, we do not require the same dilution factors (i.e. inoculum sizes) to be used in each environment. However, since the data in the baseline and focal environments are used simultaneously, the dilution factors should be normalized by the same factor $d^*$ in both cases.

To define confidence intervals (CIs) on a given parameter when the model's likelihood is a function of more than one parameter, in particular for any estimate of $\tilde{p}_c$, we use the concept of profile likelihood (ref.[12], §3.4). The profile likelihood function of a focal parameter is defined as the likelihood when holding this parameter fixed to a given value. The focal parameter's CI is in turn defined by the limits of its fixed values that allow its profile likelihood, optimized over all other parameters, to attain an optimum within a critical difference below the maximum likelihood, as optimized over all parameters including the focal. The critical difference is defined by a chi-squared test with one degree of freedom, since one parameter is fixed in the profile likelihood.

When plotting probability of population growth ($p_w$) at streptomycin concentration $x$ versus effective mean inoculum size ($\bar{N}_{\mathrm{eff}}$) calibrated from $\alpha_0^*$ in streptomycin-free media (main manuscript, Fig. 3b; and SI Appendix, Fig. S5), we plot confidence intervals around the best-fitting (MLE) curve that indicate the uncertainty relative to the x-axis. This accounts for the intuitive idea that a lower value of $\tilde{p}_c(x)$ can only sufficiently explain the data (observed population growth) in association with a higher value of $\alpha_0^*$, and vice versa. More precisely, at the lower limit of the profile likelihood CI on $\tilde{p}_c(x)$, say $\tilde{p}_{c,L}(x)$, we plot the curve $p_w = 1 - \exp(-\tilde{p}_{c,L}(x)\bar{N}_{\mathrm{eff}}\,\hat{\alpha}_{0,L}^*/\hat{\alpha}_0^*)$, where $\hat{\alpha}_0^*$ is the MLE in streptomycin-free media and $\hat{\alpha}_{0,L}^*$ is the optimized value of $\alpha_0^*$ when $\tilde{p}_c(x)$ is fixed to $\tilde{p}_{c,L}(x)$.

**Model selection:** We use the likelihood ratio test (LRT) to compare nested models (ref.[12], p. 168). The test statistic is the deviance ($D$), equal to twice the difference in maximized log likelihood between the more complex (parameter-rich) model and the more parsimonious nested model having fewer parameters. Significance of the deviance is evaluated using a chi-squared test with degrees of freedom equal to the difference in number of model parameters; if non-significant, we do not reject the simpler model. Note that the full model in terms of $p_w$, Model A or Model A′ are all equivalent, simply written in different parameterizations (and similarly for Model C or C′). These equivalent models yield the same likelihood and it does not matter which formulation is used for the LRT.

**Code:** All model fitting was implemented with custom code in R, version 3.3.1 (The R Foundation for Statistical Computing, 2016). At the core are functions to calculate the likelihood in the various models, given model parameters and data as inputs. On top of these are functions to compute parameter estimates (maximum likelihood estimates and confidence intervals) in each model through numerical optimization of the likelihood. At the outer-most layer are functions that take experimental conditions and growth data (in a data frame) as input, and return the entire model fit (all parameter estimates, the optimized log likelihood of the model, and fitted values of $p_w$ in each condition). These outputs can be readily compared across models. Finally, we use scripts to format the input data in particular experimental set-ups, call the model fitting functions, and compare the results across models using the LRT.

### 12 Generalized linear model for analyzing seeding experiment data

For the seeding experiments screening across antibiotic concentrations, with a resistant strain in isolation (Section 4), we additionally fit a generalized linear model (GLM) to the data to assess the significance of experimental variables. Specifically, we treat observed growth in replicate cultures as binomial data and fit a GLM using the built-in R function 'glm'. The response variable, probability of culture growth in a well ($p_w$), is modelled as a function of the following explanatory variables:

- natural logarithm of the dilution factor applied to the inoculating culture (treated as either continuous or categorical)
- antibiotic concentration on the treatment plates (categorical)
- experiment date, when pooling data from more than one experiment (categorical)

We use a complementary log-log (cloglog) link function. The linear predictor then takes the form:

$$\text{cloglog}(p_w) = X\beta$$

where $X$ contains the explanatory variables (i.e. the numerical value of any continuous variable and an indicator for each categorical variable) and $\beta$ contains the coefficients to be fit.

The rationale for choosing the cloglog link function is that it gives a clean relationship to our theoretical model (Section 10). Under the basic assumption that the number of established cells per replicate culture ($\alpha$) is Poisson-distributed, we have

$$p_w = 1 - \mathrm{e}^{-\alpha} \tag{S14}$$

which leads to the simple relationship

$$\text{cloglog}(p_w) := \log(-\log(1 - p_w)) = \log(\alpha) \tag{S15}$$

If we further suppose that the number of cells inoculated per culture is Poisson-distributed with mean $\bar{N}$ and each cell independently has probability $p_c$ of establishing a surviving lineage (i.e. Model C, the null model of inoculum size effect), we can substitute $\alpha = \bar{N}p_c$ and thus

$$\text{cloglog}(p_w) = \log(\bar{N}) + \log(p_c)$$

such that any explanatory variables expected to affect only inoculum size (i.e. the dilution factor applied to the inoculating culture) are separated from those expected to affect only establishment probability (i.e. antibiotic concentration) in the linear predictor. Specifically, suppose that in the $n^{\text{th}}$ observation, $\bar{N}_n = \bar{N}^*/d_n$, where $\bar{N}^*$ is the inoculum size at a particular dilution factor $d^*$, and $d_n$ is the tested dilution factor normalized by $d^*$. Further write $p_{c,n} = p_0 \cdot \tilde{p}_c(x_n)$, where $p_0$ is the per-cell establishment probability in the baseline environment (antibiotic-free media), and $\tilde{p}_c(x_n)$ is the relative establishment probability at the tested antibiotic concentration $x_n$ (cf. Eqn. S7). Then we have

$$\text{cloglog}(p_w(n)) = \log(\bar{N}^*) + \log(p_0) - \log(d_n) + \log(\tilde{p}_c(x_n))$$

This leads to the prediction that, if we treat the logarithm of the dilution factor as a continuous explanatory variable, we expect a fitted coefficient close to -1. The density of the overnight culture used for inoculation (which may vary from experiment to experiment) and the absolute establishment probability in baseline conditions (which is unknown) will be incorporated into the intercept term of the model fit. On these theoretical grounds, we treat the logarithm of dilution factor as a continuous explanatory variable in our main analyses. However, we also check that our conclusions are robust to treating dilution factor as a categorical variable, thus removing any assumptions derived from the theoretical model above (see Sections 14.1.2 and 14.2.2).

#### 12.1 Correspondence between theoretical models used for likelihood inference and GLM

We note that due to the simple form of our theoretical models (in general, Eqn. S14), and hence the natural choice of link function (Eqn. S15), we could have relied entirely on built-in GLM methods to fit our models. More specifically, we have the following correspondences:

- The saturated GLM, with dilution factor as a categorical variable, corresponds to our full model (A or A′).
- The GLM including only main effects of antibiotic and dilution factor (again categorical), but not their interaction, corresponds to our model B′.

- The GLM treating the logarithm of the dilution factor as a continuous variable, and again including only main effects, is similar, but not exactly the same, as our model C or C′. The slight difference arises because the theoretical model assumes perfect dilution steps (fixed values), whereas the GLM fits one coefficient scaling the effect of log(dilfac). Thus, there is one more estimated parameter in the GLM, which according to our theoretical expectations, should be approximately -1. Values closer to -1 reflect more accurate dilution steps in the experiment, and correspond to a closer match between the GLM and theoretical model.

Despite this correspondence, there are several reasons to use our custom-coded model fitting approach. Firstly, it gives likelihood-based confidence intervals that extend to the estimates of relative establishment probabilities based on experimental data in more than one environment. Secondly, it can readily be fit to the data even when none or all of the replicates in a given condition show growth. Finally, likelihood-based fitting approaches can be applied to any stochastic model, so this framework could be extended to more complex theoretical models of population growth, even if a built-in link function is not available. For instance, extensions could include modelling hypothetical mechanisms of interactions between individuals, or using functional forms of the establishment probability derived from an underlying demographic model.

#### Part III

### Supplementary Statistical Results

#### 13 Distribution of the number of colony-forming units

Using the YFP:Rms149 strain, we tested whether the number of CFUs in a small volume of highly diluted culture can be sufficiently described by a Poisson distribution (Section 3). A Poisson distribution was fit to the colony counts per plated $4\mu l$ spot using the sample mean. To conduct a goodness-of-fit test, the data (colony count in each spot) were grouped into categories according to the guideline that there should be an expected number of at least five observations per category (ref.[13] p. 540). Deviation from the Poisson distribution was determined at a 5% significance level in a chi-squared test with degrees of freedom equal to the number of categories minus two (ref.[13] p. 540). The test statistic is $c_1 = \sum_i \frac{(n_i - np_i)^2}{np_i}$ where $n$ is the total number of observations, $n_i$ is the number of observations in category $i$, and $p_i$ is the probability of an observation falling in category $i$ in the fitted distribution (thus, $np_i$ is the expected number of observations in category $i$). We carried out two separate experiments, each with 144 plated spots. In both cases, we could use categories of 0, 1, 2, 3, 4, and 5 or more colonies per spot.

- Experiment 1 (SI Appendix, Fig. S1a): The sample mean is 1.97 (and variance is 1.68). According to the goodness-of-fit test, the deviation from a Poisson distribution is not significant ($c_1 = 7.76$; $\chi^2_4$: $p = 0.10$) and thus we do not reject the null hypothesis that the data are drawn from a Poisson distribution.

- Experiment 2 (SI Appendix, Fig. S1b): The sample mean is 1.96 (and variance is 1.77). Again we do not reject the Poisson distribution ($c_1 = 2.05$; $\chi^2_4$: $p = 0.73$).

### 14 Seeding experiments to estimate establishment probability of the resistant strain in isolation, across antibiotic concentrations

Here we report detailed results of model fitting to data from our seeding experiments with the resistant strain in isolation, screening across antibiotic concentrations (Section 4). For each experiment, we first report the results of fitting our theoretical models of population growth (Sections 10-11) to the data. The model selected by the likelihood ratio test is used to obtain the maximum likelihood estimates and confidence intervals of relative establishment probability ($\tilde{p}_c$), as plotted in the main manuscript, Fig. 2 and 5, and reported in the SI Appendix, Tables S2 and S4, for the Rms149 strain in streptomycin and the PAMBL2 strain in meropenem, respectively. Next, we fit a generalized linear model (GLM) to the growth data (Section 12), which is used to determine significant effects of antibiotic concentration as annotated on Fig. 2 and 5.

#### 14.1 Resistant (Rms149) strain alone, in streptomycin

We first consider the data from two repeat experiments seeding the Rms149-carrying (streptomycin-resistant) strain, in isolation, into media containing various concentrations of streptomycin (0, 1/64, 1/32, 1/16, 1/8 × $\mathrm{MIC}_R$, where $\mathrm{MIC}_R$=2048$\mu$g/ml). These results are presented in the main manuscript, Fig. 2, and SI Appendix, Table S2.

##### 14.1.1 Theoretical model fitting

Recall that Model A (equivalently, Model A′ in the transformed parameterization) is the full model, with the number of estimated parameters equal to the number of tested conditions: (# inoculating dilution factors) × (# streptomycin concentrations) = 15 in these experiments.

Model B$'$ allows a separate estimate of mean number of established cells in streptomycin-free media at each inoculating dilution factor, but assumes relative establishment probability, $\tilde{p}_c(x)$, at each streptomycin concentration $x \neq 0$, is common to all dilution factors. This yields a total number of parameters equal to (# dil. factors) + (# non-zero Strep. conc.) = 7. Finally, Model C (or C$'$), the null model of the inoculum size effect, additionally assumes that the number of established cells scales proportional to inoculum size (inversely proportional to dilution factor), leaving 1 + (# non-zero Strep. conc.) = 5 parameters to estimate. We fit each model to the data, i.e. number of replicate cultures showing growth, using maximum likelihood estimation of the parameters. Since these models are nested, we then use the likelihood ratio test (LRT) to compare their fits to the data. If the deviance ($D$) between the more complex and the simpler nested model is non-significant in a chi-squared test with degrees of freedom (d.f.) equal to the difference in number of estimated parameters, we do not reject the simpler model. (Here we sum the contributions to $D$ from each tested streptomycin concentration, to get an overall model choice for the entire data set.) The simplest model not rejected by the LRT is selected to obtain the reported estimates of relative establishment probability, $\tilde{p}_c$.

**Experiment 1:**

- Model A$'$ vs. B$'$: $D = 3.56$, d.f. = 8, $p = 0.89 \Rightarrow$ do not reject B$'$
- Model B$'$ vs. C$'$: $D = 15.8$, d.f. = 2, $p =$ 3.6e-4 $\Rightarrow$ reject C$'$

Further analysis suggests that the lowest one of the three inoculating dilution factors primarily contributed to the deviation of Model C$'$ in this experiment, suggesting an inaccuracy in preparation of this particular dilution that resulted in lack of proportionality in inoculum sizes as assumed by this model. Indeed, if the data from this dilution factor are excluded from the analysis, Model C$'$ is not rejected ($D = 0.210$, d.f.= 1, $p = 0.65$ compared to Model B$'$). For comparison, dropping either of the other single dilution factors still results in rejection of Model C$'$, suggesting that the test is not simply under-powered on the reduced data set, but rather that the identified dilution factor is the source of error. We report results from the fit of Model B$'$ including data from all dilution factors. However, we obtain similar point estimates for $\tilde{p}_c$ either with or without data from this dilution factor, and fitting either Model B$'$ or C$'$.

**Experiment 2:**

- Model A$'$ vs. B$'$: $D = 8.99$, d.f. = 8, $p = 0.34 \Rightarrow$ do not reject B$'$
- Model B$'$ vs. C$'$: $D = 1.65$, d.f. = 2, $p = 0.44 \Rightarrow$ do not reject C$'$

Thus, in this experiment, we select Model C$'$.

##### 14.1.2 Generalized linear model fitting

In the GLM, the explanatory variables are streptomycin concentration, Strep for short; the natural logarithm of the inoculating dilution factor, log(dilfac) for short; and experiment date, if applicable (when pooling data from two experiments). These results are used to determine the significance of streptomycin effects as annotated on Fig. 2 in the main manuscript.

**Experiment 1:** Fitting a saturated model identified that the log(dilfac) and Strep main effects were significant, but their interaction was not. A reduced model including only the main effects was correspondingly preferred by the Akaike Information Criterion (AIC), as calculated by the built-in 'glm' function in R: AIC=90.2 for the saturated model vs. 83.7 for the reduced model if log(dilfac) is treated as a continuous variable, or 96.1 vs. 83.7 if treated as a categorical variable.

Taking log(dilfac) as continuous, the reduced model fit (Table III.1) indicated that the effects of Strep at $1/16\times\text{MIC}_R$ and $1/8\times\text{MIC}_R$ are significant relative to the streptomycin-free conditions. As expected, the effect of log(dilfac) is also highly significant; furthermore, the fitted coefficient of $-1.30$ is reasonably close to the theoretical prediction of -1 (see Section 12), but presumably skewed by the suspected error in one dilution factor mentioned above. Treating log(dilfac) instead as categorical does not change the conclusions regarding significant effects (lowest vs. highest dilution factor: $p <$ 2e-16; middle vs. highest dilution factor: $p$=3e-10; Strep at $1/16\times\text{MIC}_R$: $p$=0.010; Strep at $1/8\times\text{MIC}_R$: $p$ <2e-16).

Table III.1: GLM results: reduced model[a] fit, seeding experiment 1 with Rms149 in streptomycin

| | Estimated coefficient | Standard error | $z$-value | $p$-value |
|---|---|---|---|---|
| intercept | 0.753 | 0.095 | 7.89 | 3.1e-15 |
| log(dilfac)[b] | -1.299 | 0.081 | -16.0 | <2e-16 |
| 1/64× Strep[c] | -0.024 | 0.120 | -0.201 | 0.84 |
| 1/32× Strep[c] | -0.078 | 0.120 | -0.644 | 0.52 |
| 1/16× Strep[c] | -0.315 | 0.123 | -2.56 | 1.0e-2 |
| 1/8× Strep[c] | -3.024 | 0.273 | -11.1 | <2e-16 |

[a] The model selected based on AIC, containing only the effects listed in the table.

[b] Natural logarithm of the dilution factor, taken as a continuous variable.

[c] Streptomycin concentrations, scaled by $\text{MIC}_R$, taken as categorical variables with streptomycin-free as the baseline.

**Experiment 2:** Taking log(dilfac) as continuous, fitting a saturated model again indicated that the log(dilfac) and Strep main effects were significant, but their interaction was not; the reduced model excluding the interaction term was correspondingly preferred by AIC (saturated: 86.5 vs. reduced: 81.6). The reduced model fit (Table III.2) again indicated that the effects of Strep at 1/16×$\text{MIC}_R$ and at 1/8×$\text{MIC}_R$ are significant. The effect of log(dilfac) is also highly significant, and the fitted coefficient of $-1.01$ is in excellent agreement with the theoretical prediction, consistent with the acceptance of the theoretical Model C′ in the previous analysis.

When treating log(dilfac) as categorical, we cannot fit the saturated model to the complete data set, because zero replicates showed growth at the highest dilution factor and 1/8× Strep, i.e. there is no variation in this category. Excluding this single data point, the saturated model (AIC: 87.0) again yields significant main effects of log(dilfac) and Strep but a non-significant interaction; the reduced model (AIC: 77.0) was correspondingly preferred, and indicated significant effects consistent with the continuous-variable analysis (lowest vs. highest and middle vs. highest dilution factor: both $p <$ 2e-16; Strep at 1/16×$\text{MIC}_R$: $p =$ 2e-7; Strep at 1/8× $\text{MIC}_R$: $p <$ 2e-16).

Table III.2: GLM results: reduced model[a] fit, seeding experiment 2 with Rms149 in streptomycin

| | Estimated coefficient | Standard error | $z$-value | $p$-value |
|---|---|---|---|---|
| intercept | 1.276 | 0.101 | 12.7 | <2e-16 |
| log(dilfac)[b] | -1.009 | 0.076 | -13.3 | <2e-16 |
| 1/64× Strep[c] | -0.029 | 0.115 | -0.247 | 0.81 |
| 1/32× Strep[c] | 0.017 | 0.116 | 0.145 | 0.89 |
| 1/16× Strep[c] | -0.606 | 0.116 | -5.22 | 1.8e-7 |
| 1/8× Strep[c] | -3.585 | 0.264 | -13.6 | <2e-16 |

[a] The model selected based on AIC, containing only the effects listed in the table.

[b] Natural logarithm of the dilution factor, taken as a continuous variable.

[c] Streptomycin concentrations, scaled by $\text{MIC}_R$, taken as categorical variables with streptomycin-free as the baseline.

**Pooling both experiments:** Using experiment date as an additional explanatory variable, a hierarchical search according to minimal AIC (applying the built-in R function 'step' to the saturated model) identified a reduced model in which all main effects and the experiment date

$\times$ log(dilfac) interaction effect are retained, regardless of whether log(dilfac) is treated as continuous or categorical (see Table III.3 for the model fit in the continuous case).

The experiment data $\times$ log(dilfac) interaction presumably arises because of the suspected inaccuracy in the lowest dilution factor only in Experiment 1, as described above. Indeed, the interaction is identified as significant ($p = 0.003$) with log(dilfac) taken as continuous, while with log(dilfac) taken as categorical with the highest dilution factor as the baseline, the interaction between experiment date and the lowest dilution factor is significant ($p = 0.008$), but the interaction with the middle dilution factor is not significant ($p = 0.8$). Taken together, these results again point to error in a single dilution step in Experiment 1 as the source of deviating effects; we do not interpret this result as having biological significance.

More importantly, pooling data from two experiments strengthens the conclusions regarding the effect of streptomycin: specifically, Strep at $1/16\times\text{MIC}_R$ or $1/8\times\text{MIC}_R$ has a highly significant effect relative to the streptomycin-free control ($p$=2e-8 and $p$ <2e-16, respectively, regardless of whether log(dilfac) is treated as continuous or categorical).

Table III.3: GLM results: reduced model[a] fit, pooled seeding experiments with Rms149 in streptomycin

| | Estimated coefficient | Standard error | $z$-value | $p$-value |
|---|---|---|---|---|
| intercept | 0.790 | 0.079 | 10.0 | <2e-16 |
| expt2[b] | 0.441 | 0.089 | 4.95 | 7.4e-7 |
| log(dilfac)[c] | -1.312 | 0.082 | -16.1 | <2e-16 |
| expt2 : log(dilfac) | 0.324 | 0.110 | 2.95 | 3.2e-3 |
| 1/64× Strep[d] | -0.026 | 0.083 | -0.312 | 0.76 |
| 1/32× Strep[d] | -0.028 | 0.083 | -0.336 | 0.74 |
| 1/16× Strep[d] | -0.474 | 0.084 | -5.62 | 1.9e-8 |
| 1/8× Strep[d] | -3.340 | 0.190 | -17.6 | <2e-16 |

[a] The model selected based on AIC, containing only the effects listed in the table.

[b] Experiment identifier, taken as a categorical variable with experiment 1 (expt1) as the baseline.

[c] Natural logarithm of the dilution factor, taken as a continuous variable.

[d] Streptomycin concentrations, scaled by $\text{MIC}_R$, taken as categorical variables with streptomycin-free as the baseline.

#### 14.2 Resistant (PAMBL2) strain alone, in meropenem

We now turn to the seeding experiment data for the PAMBL2-carrying (meropenem-resistant) strain, seeded in isolation into media containing meropenem at various concentrations (0, 1/32, 1/16, 1/8, 1/4 $\times$ $\mathrm{MIC}_R$, where $\mathrm{MIC}_R = 512\mu$g/ml). These results are presented in the main manuscript, Fig. 5, and SI Appendix, Table S4.

##### 14.2.1 Theoretical model fitting

Similarly to the previous experiments, we fit the theoretical models A′, B′ and C′ to the growth data, and compared their fits using the likelihood ratio test (LRT). Note that we used five antibiotic concentrations but only two inoculating dilution factors in this experiment, implying that there are 10 parameters to fit in Model A′, six in Model B′, and five in Model C′. The LRT yields:

- Model A′ vs. B′: $D = 3.17$, d.f.$= 4$, $p = 0.53 \Rightarrow$ do not reject B′
- Model B′ vs. C′: $D = 0.067$, d.f.$= 1$, $p = 0.80 \Rightarrow$ do not reject C′

Thus, we select Model C′.

##### 14.2.2 Generalized linear model fitting

In the GLM, the explanatory variables are meropenem concentration, Mero for short; and the logarithm of the inoculating dilution factor, log(dilfac) for short. The significance of meropenem effects as determined from the GLM fit is annotated on Fig. 5 in the main manuscript.

Since there is no variation in the data for 1/4 $\times$ Mero (zero replicates established at both dilution factors), these data are excluded from the GLM. Fitting to the remaining data (four meropenem concentrations at each of two dilution factors), we find that a reduced model including only main effects is preferred over a saturated model additionally including Mero $\times$ log(dilfac) interactions (AIC: 47.9 for the saturated model vs. 45.0 for the reduced model). The reduced model fit indicates that log(dilfac) has a highly significant effect ($p <$ 2e-16), with the fitted coefficient of -1.02 (when treated as a continuous variable) again very close to the theoretical prediction. Meropenem at 1/16 $\times$ $\mathrm{MIC}_R$ has a marginally significant effect ($p = 0.024$) while at 1/8 $\times$ $\mathrm{MIC}_R$ the result is highly significant ($p <$ 2e-16), regardless of whether log(dilfac) is treated as continuous or categorical (see Table III.4).

Table III.4: GLM results: reduced model[a] fit, seeding experiment with PAMBL2 in meropenem

| | Estimated coefficient | Standard error | $z$-value | $p$-value |
|---|---|---|---|---|
| intercept | 1.284 | 0.129 | 9.94 | <2e-16 |
| log(dilfac)[b] | -1.022 | 0.087 | -11.8 | <2e-16 |
| 1/32× Mero[c] | -0.024 | 0.151 | -0.162 | 0.87 |
| 1/16× Mero[c] | -0.336 | 0.149 | -2.25 | 0.024 |
| 1/8× Mero[c] | -2.986 | 0.252 | -11.9 | <2e-16 |

[a] The model selected based on AIC, containing only the effects listed in the table.

[b] Natural logarithm of the dilution factor, taken as a continuous variable.

[c] Meropenem concentrations, scaled by $MIC_R$, taken as categorical variables with meropenem-free as the baseline.

### 15 Seeding experiments to test the null model of the inoculum size effect

Here we report the complete results of fitting our theoretical models of population growth (Sections 10-11) to experimental data testing the effect of inoculum size at a given streptomycin concentration, using the Rms149 strain (Section 5). Specifically, at each streptomycin concentration, we test whether the null model (Model C or equivalently C′) is not rejected, by the likelihood ratio test, in comparison to the full model (Model A or A′) that allows an arbitrary effect of each inoculum size. The results of the "main experiment" (testing both $1/16\times MIC_R$ and $1/8\times MIC_R$ streptomycin, as well as streptomycin-free conditions) are illustrated in the main manuscript, Fig. 3b, and SI Appendix, Fig. S4; the results of the two "supplementary experiments" (one testing $1/16\times MIC_R$ and one testing $1/8\times MIC_R$ streptomycin, along with a streptomycin-free control in each) are illustrated in SI Appendix, Fig. S5; and estimates of relative establishment probability $\tilde{p}_c$ from all experiments are summarized in SI Appendix, Table S2. In Table III.5 we summarize the results of the likelihood ratio test at each streptomycin concentration in each experiment, based on growth evaluated at either 3d or 5d post-inoculation in the presence of streptomycin. In all but one case, the deviance ($D$) of the null model from the full model (calculated separately at each streptomycin concentration) is non-significant ($p > 0.05$), and thus we do not reject the null model. In supplementary experiment 2, if growth is evaluated at 3d, the deviance becomes marginally significant ($p = 0.48$), but does not remain significant after correcting for multiple testing.

Table III.5: Testing the null model of inoculum size effect: likelihood ratio test results

| Experiment | [Strep] $\times MIC_R$ | # inoc. sizes tested | Deviance[a] (null from full) | $p$-value[b] | Effective density[c] of overnight culture (viable cells/ml) |
|---|---|---|---|---|---|
| main | 0 | 5 | 3.08 | 0.55 | $7.58 \times 10^9$ |
| | 1/16 | 9 | *at 3d:* 9.81<br>*at 5d:* 9.74 | 0.28<br>0.28 | |
| | 1/8 | 6 | *at 3d:* 5.65<br>*at 5d:* 2.91 | 0.34<br>0.71 | |
| suppl. 1 | 0 | 5 | 8.57 | 0.073 | $7.17 \times 10^9$ |
| | 1/16 | 9 | *at 3d:* 1.02<br>*at 5d:* 1.02 | 1.00<br>1.00 | |
| suppl. 2 | 0 | 5 | 0.347 | 0.99 | $9.10 \times 10^9$ |
| | 1/8 | 10 | *at 3d:* 17.0<br>*at 5d:* 13.1 | 0.048<br>0.16 | |

[a] Deviance ($D$) of the null model from the full model, defined as twice the difference in maximal log likelihood of each model (see Section 11).

[b] $p$-value in the likelihood ratio test, i.e. a chi-squared test with $D$ as the test statistic and degrees of freedom = (# parameters in full model) – (# parameters in null model) = (# tested inoculum sizes – 1).

[c] In streptomycin-free conditions, we use the maximum likelihood estimate of $\alpha_0^*$, scaled up by the corresponding dilution factor applied for the inoculation, to estimate an effective viable cell density in the overnight culture used for inoculation. This is equivalent to the "most probable number" method for determining bacterial density using multiple dilution factors[16]. We use this estimate, along with the known dilution factors applied to the inoculating culture, to scale the "effective mean inoculum size" on the x-axis of our plots.

### 16 Seeding experiments to estimate establishment probability of the resistant strain in the presence of the sensitive strain

Here we present detailed analysis of experiments seeding the resistant (Rms149-carrying) strain in the presence of the sensitive strain, across streptomycin concentrations (Section 6; results presented in the main manuscript, Fig. 6, and SI Appendix, Table S6).

#### 16.1 Growth of sensitive strain alone

As controls, we tested growth of the sensitive strain alone, at each streptomycin concentration. Table III.6 reports these growth data. The frequent growth of cultures in $1\times\text{MIC}_S$ streptomycin when inoculated at high density (approx. $5 \times 10^7$ CFU/ml) is consistent with previous observations that MIC often increases with inoculation density[14;15]; recall that the standard $\text{MIC}_S$ was evaluated at inoculation density similar to the "low" density here (approx. $5\times10^5$ CFU/ml). Occasionally, at streptomycin concentrations well above the MIC of the sensitive strain ($4\times\text{MIC}_S$ and higher), replicate cultures inoculated with the sensitive strain alone (see Table III.6) showed growth but low fluorescence. We interpret these cases as probable outgrowth of spontaneous resistant mutants on the DsRed sensitive strain background (or possibly cross-contamination with unrelated, streptomycin-resistant bacteria in the lab). Notably, all of these cases occurred when the sensitive strain was inoculated at high density, providing 100-fold higher initial population size from which mutants could have arisen than the low-density inoculum.

Table III.6: Number of sensitive strain monocultures showing growth, out of 24 replicates per condition

| Experiment | Inoculation density[a] | Streptomycin concentration ($\times\text{MIC}_S$) | | | | | | |
|---|---|---|---|---|---|---|---|---|
| | | 0 | 1/4 | 1 | 4 | 8 | 16 | 32 |
| 1 | low (2.67 $\times10^5$ CFU/ml) | 24 | 24 | 1 | 0 | 0 | N/A | N/A |
| | high (2.67 $\times10^7$ CFU/ml) | 24 | 24 | 23 | 0 | 0 | N/A | N/A |
| 2 | high (4.73 $\times10^7$ CFU/ml) | 24 | N/A | N/A | N/A | 2 | 0 | 1 |

[a] Estimated by plating.

#### 16.2 Seeding data: Theoretical model fitting

When the resistant strain was seeded into the sensitive population, we took the number of replicate cultures showing growth specifically by the resistant strain, as determined by fluorescence (see Section 6), as data for model fitting. The "baseline condition", against which relative establishment probabilities were normalized, was chosen as antibiotic-free media in the absence of the sensitive strain. As before, model selection was based on the likelihood ratio test.

**Experiment 1 (lower streptomycin concentration range):** We tested 15 environmental conditions (5 streptomycin concentrations $\times$ 3 sensitive densities, including the absence of the sensitive), each with 2 inoculating dilution factors of the resistant strain. This yields 30 fitted parameters in the full model (A′), 16 in Model B′, and 15 in Model C′.

- Model A′ vs. B′: $D = 13.6$, d.f.$= 14$, $p = 0.48 \Rightarrow$ do not reject B′

- Model B′ vs. C′: $D = 2.30$, d.f.$= 1$, $p = 0.13 \Rightarrow$ do not reject C′

Thus, in this experiment, we select Model C′.

**Experiment 2 (higher streptomycin concentration range):** We tested 8 environmental conditions (4 streptomycin concentrations $\times$ 2 sensitive densities), each with 2 inoculating dilution factors of the resistant strain. This yields 16 fitted parameters in the full model (A′), 9 in Model B′, and 8 in Model C′.

- Model A′ vs. B′: $D = 2.19$, d.f.$= 7$, $p = 0.95 \Rightarrow$ do not reject B′

- Model B′ vs. C′: $D = 0.319$, d.f.$= 1$, $p = 0.57 \Rightarrow$ do not reject C′

Thus, in this experiment, we select Model C′.

##### 16.3 Seeding data: Assessing significance of experimental conditions

The seeding data showed too many conditions in which zero replicates established for us to effectively fit a GLM to the full dataset. Instead, we focused on comparing selected pairs of conditions that were of particular interest, and in which significance of the effect of the sensitive population on resistant establishment was not immediately clear. For this purpose, we treated each seeding replicate as a Bernoulli trial (with "success" being establishment of resistance in that culture), and compared pairs of experimental conditions using a two-sided Wilcoxon rank-sum test, with a Bonferroni correction for multiple testing. For this purpose, we pooled data across the two inoculated dilution factors of the resistant strain; since the Wilcoxon test is non-parametric, it does not matter that the distribution of the pooled data will be a weighted sum of two binomial distributions, with different success probabilities at different inoculum sizes. In experiment 1, we also pooled data across absence and presence of the sensitive strain at low density, which showed very similar results, to increase the power to detect an effect of the sensitive strain at high density.

Specifically, we tested the following four pairs of conditions, and used a threshold $p$-value of 0.05/4=0.0125 to determine significance after Bonferroni correction. Test statistics and approximate $p$-values were determined using the built-in R function 'wilcox.test'.

- Experiment 1, at $1/16\times\text{MIC}_R$: resistant establishes in 86/240 cultures (no/low-density sensitive) vs. 52/120 cultures (high-density sensitive). Wilcoxon test: $W = 15480$, $p = 0.17$.

- Experiment 1, at $1/8\times\text{MIC}_R$: resistant establishes in 2/240 cultures (no/low-density sensitive) vs. 45/120 cultures (high-density sensitive). Wilcoxon test: $W = 19680$, $p < 2.2 \times 10^{-16}$.

- Experiment 2, at $1/8 \times \text{MIC}_R$: resistant establishes in 0/60 cultures (no sensitive) vs. 45/120 cultures (high-density sensitive). Wilcoxon test: $W = 4950$, $p = 4.8 \times 10^{-8}$.
- Experiment 2, at $1/4 \times \text{MIC}_R$: resistant establishes in 0/60 cultures (no sensitive) vs. 8/120 cultures (high-density sensitive). Wilcoxon test: $W = 3840$, $p = 0.042$.
